# Cancer-Associated Mutations Perturb the Structure and Interactions of the Intrinsically Disordered p53 Transactivation Domain

**DOI:** 10.1101/2020.12.12.422504

**Authors:** Lynn G. Schrag, Indhujah Thevarajan, Xiaorong Liu, Om Prakash, Michal Zolkiewski, Jianhan Chen

## Abstract

Intrinsically disordered proteins (IDPs) are key components of regulatory networks that control crucial aspects of cell decision making. The intrinsically disordered transactivation domain (TAD) of tumor suppressor p53 mediates its interactions with multiple regulatory pathways to control the p53 homeostasis during the cellular response to genotoxic stress. Many cancer-associated mutations have been discovered in p53-TAD, but their structural and functional consequences are poorly understood. Here, by combining atomistic simulations, NMR spectroscopy, and binding assays, we demonstrate that cancer-associated mutations can significantly perturb the balance of p53 interactions with key activation and degradation regulators. Importantly, mutations do not all directly disrupt the known interaction interfaces. Instead, some mutations likely modulate the disordered state of p53-TAD, which affects the interactions. Our work suggests that the disordered conformational ensemble of p53-TAD can serve as a central conduit in regulating the response to various cellular stimuli at the protein-protein interaction level. Understanding how the disordered state of IDPs may be modulated by regulatory signals and/or disease associated perturbations will be essential in the studies on the role of IDPs in biology and diseases.

## Introduction

Intrinsically disordered proteins (IDPs), which lack stable 3D structures under physiological conditions, have challenged the traditional protein structure-function paradigm^1-5^. IDPs tend to contain low-complexity sequences, are enriched with charged residues, and display low overall hydrophobicity^6-7^. Importantly, IDPs do not usually exhibit fully random conformations in the absence of binding partners^8^. Frequently, they display a heterogeneous distribution of partially folded but highly mobile conformations^9^. The inherent thermodynamic instability of IDP conformations arguably provides a robust mechanism for allosteric regulation^10-14^, by allowing the disordered ensemble to efficiently respond to various cellular signals. Combinations of post-translational modifications, ligand or protein binding, changes in environmental conditions (pH, ionic strength, etc.), and mechanical stress are common regulators of IDP conformational ensembles and interaction interfaces^15-20^. Cooperative responses of IDPs, such as binding-induced folding^21^, could naturally integrate multiple signal pathways, thus allowing them to function as hubs within cellular signaling networks^15^. Indeed, IDPs have been estimated to account for nearly half of signaling-associated proteins in eukaryotic cells^22^, and nearly all eukaryotic transcription factors contain one or more disordered regions^23^. Thus, it is not surprising that many human diseases, including neurodegeneration, cancer, and diabetes have been linked to mis-function of IDPs^24-27^. There is thus an important demand and great interest in understanding how intrinsic disorder mediates versatile biological functions and how such mechanisms may fail in human diseases.

A prominent example of a biologically important IDP is the transactivation domain (TAD) of the tumor suppressor and transcription factor p53. As a critical integrator of cellular responses against genotoxic stress, p53 is the most frequently mutated protein associated with human cancers^28-30^. p53 is a multidomain protein comprised of the TAD, proline-rich domain (PRD), DNA-binding domain (DBD), tetramerization domain (TD), and negative regulatory domain (NRD) (**Figure 1A**). Only the DBD is stably folded in the unbound state of p53. In healthy cells, p53 is negatively regulated via targeting for degradation by E3 ubiquitin ligase Human doubleminute2 (HDM2), which binds p53-TAD and promotes poly-ubiquitination of the p53 NRD^31-32^ (see **Figure 1B**). Upon prolonged exposure to genotoxic stressors, p53 accumulates post-translational modifications at multiple positions^33-40^. Key phosphorylation events within TAD, including those at Ser15, Thr18, and Ser20 destabilize the p53-TAD:HDM2 complex, inhibit the HDM2-mediated poly-ubiquitination of p53, and prevent p53 degradation^41-44^. Moreover, multi-site phosphorylation of p53-TAD recruits the histone acetyl transferases and transcriptional co-activators, cyclic-AMP response element-binding (CREB)-binding protein (CBP) and its paralog p300, to p53-TAD^34, 45-50^. These interactions enable site-specific acetylation of p53 NRD, which promotes p53 nuclear localization and further stabilizes the complex with CBP/p300^51-52^ (see **Figure 1B**). These processes eventually lead to the activation of p53-regulated apoptotic genes^53-55^.

**Figure 1.**
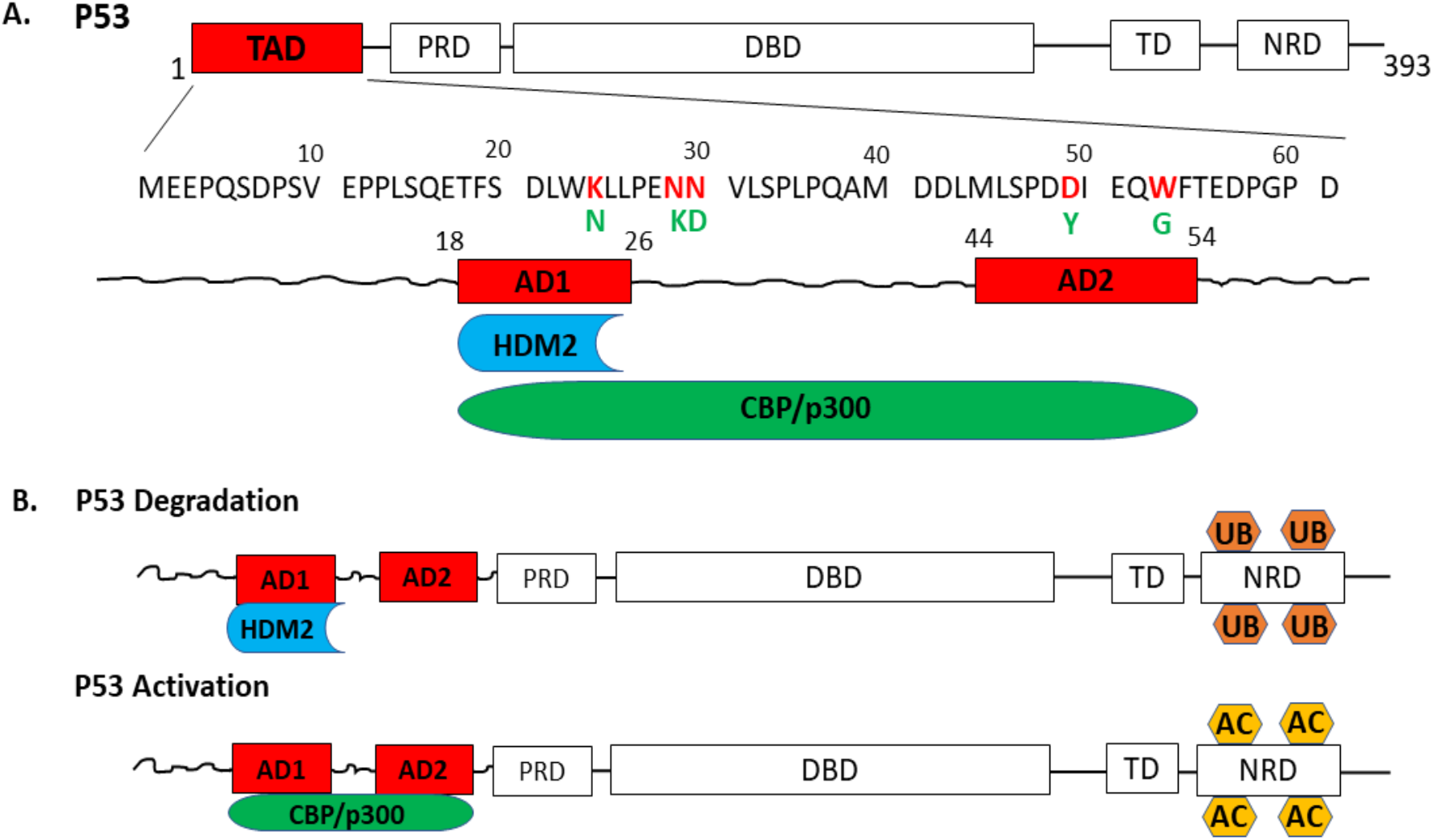
p53-TAD domain structure, interactions and regulation. (A) Domain structure of p53: TAD (transactivation domain), PRD (proline-rich domain), DBD (DNA-binding domain), TD (tetramerization domain), and NRD (negative regulatory domain). The sequence of human p53-TAD is shown with the selected cancer-linked mutation positions marked in red and the amino acid substitutions shown in green. The helical segments within TAD, AD1 and AD2, are shown below with the interacting partners, HDM2 and CBP/p300. (B) p53 regulation is mediated by HDM2 and CBP/p300. Top: In unstressed cells, HDM2 binds to AD1 and negatively regulates p53 stability by polyubiquitination (UB) of NRD. Bottom: Under genotoxic stress, phosphorylation of TAD inhibits HDM2-p53-TAD interaction and promotes the formation of a complex with CBP/p300, which leads to acetylation (AC) of NRD, and to activation and stabilization of p53.

The molecular mechanism of modulation of the balance of p53 interactions with the degradation and activation pathways by multi-site TAD phosphorylation is not fully understood. Many phosphorylation sites are not located at and near the known interaction sites of p53-TAD and thus do not directly disrupt protein-protein interactions^56^. Alternatively, phosphorylation could modulate the unbound conformational ensemble of p53-TAD, which might affect its interactions. Specifically, the overall shape, transient secondary structures and/or long-range contacts in p53-TAD may be perturbed, which could alter the availability of key binding residues and/or change the entropic cost of binding to various regulators^56^. Such conformational modulation mechanisms have been demonstrated in previous studies of CREB/CBP^57^, Connexin43 and 4E-BP2^58-59^, as well as interaction of IDPs with several drug molecules^60-64^. Importantly, although cancer-associated mutations in p53 are predominantly located within its DBD, a large number of clinically relevant residue substitutions occur within the TAD^65^. Many of these cancer-associated mutations occur outside of the known interaction sites of p53-TAD. It is plausible that these mutations could also modulate the disordered ensemble of p53-TAD to perturb the balance of its interactions and further interfere with the regulation of the p53 homeostasis during a cellular response to genotoxic stresses.

Testing the above hypotheses is challenging and requires a detailed analysis of the disordered conformational ensemble of p53-TAD. p53-TAD has been previously characterized using various experimental methods, including NMR spectroscopy^66-70^, small-angle X-ray scattering (SAXS)^66^, and single molecule fluorescence resonance energy transfer (smFRET)^71^. Those studies show that an unbound p53-TAD contains partial helical structures in the regions AD1 and AD2 (see **Figure 1**)^66-67^, with the AD1 region estimated to be up to ∼30% helical. The overall TAD shape appears to be extended with a Stokes radius of ∼2.4 nm (residues 1-73) according to size exclusion chromatography^66, 70-71^. Transient long-range contacts have also been observed within TAD^68, 72^. Critically, these experimental approaches measure ensemble-averaged properties and do not directly reveal the full range of potentially heterogeneous conformational states sampled by TAD. In contrast, molecular dynamics (MD) simulations employing physics-based force fields could provide the atomistic details of the disordered ensemble to assist in understanding the molecular basis of its function^14^. This challenging task has become increasingly feasible in recent years, thanks to a continuous development of more accurate protein force fields^73-77^, highly efficient GPU-enabled MD algorithms^78-84^, and advanced conformational sampling algorithms^85-97^. The simulated ensembles need to be properly validated by a direct comparison with experimental observables and by assessing their ability to recapitulate the effects of various perturbations such as mutations, covalent modifications, and binding^14^. Combined information from atomistic simulation and biophysical characterization can thus provide the most comprehensive molecular image for dissecting the mechanistic link between the disordered conformational ensemble and protein function, and for revealing how such mechanisms may fail in human diseases.

In this study, we investigated four variants of p53-TAD, each with one or two amino-acid substitution(s) found in human cancers^65, 98-100^, namely, K24N, N29K/N30D, D49Y, and W53G. These mutants have been associated with female genital, brain, bladder or breast cancers^65^, and were previously predicted to substantially modulate the level of disorder in unbound p53-TAD^56^. By combining protein-protein interaction assays, NMR spectroscopy, and atomistic simulation with an aggregated enhanced sampling time of over 240 µs, we assessed functional consequences and conformational ensemble changes in p53-TAD resulting from the above missense mutations. The results show that, while the TAD structure and dynamics are only marginally affected, these cancer-associated mutations have profound effects upon the TAD interactions with the partner domains of its key regulators HDM2 and CBP. Our results demonstrate that subtle changes in the conformational ensemble of an IDP may produce distinct molecular responses, which highlights a central role of the ensemble dynamics in regulating the IDP’s response to various cellular stimuli.

## Results

### Cancer mutations perturb p53-TAD’s interactions with key regulators

Although many mutations found in disordered regions of proteins have been associated with diseases^101^, it is not obvious whether these mutations have direct functional consequences at the molecular level. For human p53-TAD (residues 1-73), we examined the functional impact of the four selected TAD variants, K24N, N29K/N30D, D49Y, and W53G, by studying their interactions with the partner domains of the key regulators HDM2 and CBP (see **Figure 1**). For these studies, we produced the recombinant N-terminal region of the E3 ubiquitin ligase HDM2 (residues 17-125), which is sufficient for binding to p53-TAD^102^. CBP contains four domains, TAZ1, KIX, TAZ2, and NCBD that all interact with p53-TAD, but binding affinities for TAZ1 and TAZ2 are significantly higher than those for KIX and NCBD^50^. Therefore, we focused on the interactions of the p53-TAD variants with TAZ1 and TAZ2 domains, because they could best reflect how CBP binding to p53 may be impacted by the cancer-associated mutations.

We conducted a series of *in vitro* binding assays utilizing biolayer interferometry (BLI) with the p53-TAD variants non-covalently immobilized on Ni-NTA biosensor tips. The p53-TAD variants were exposed to 3 different concentrations for each interaction partner and the BLI responses were recorded with 2 experimental replicates (**Figures 2, S1-3)**. As shown in Figure 2, each of the tested partner domains (HDM2, TAZ1, TAZ2) interacted with the biosensor tip even in the absence of p53-TAD (control “non-specific” traces). However, the BLI signal response clearly exceeded the “non-specific” level for each of the partner domains during their interactions with wild type p53-TAD. The association and dissociation phases from all 6 measurements for each interaction were used to calculate the association (*k*_on_) and dissociation (*k*_off_) rate constants and the equilibrium dissociation constant (*K*_D_) (**Table 1**). The *k*_on_ and *k*_off_ values for p53-TAD^WT^ indicate fast association/dissociation kinetics for binding of all three partners, in agreement with the previous NMR measurements^50^. Importantly, the *K*_D_ values from our BLI experiments (see **Table 1**) reproduce those for p53-TAD^WT^ interactions with HDM2, TAZ1, and TAZ2, as previously determined using isothermal titration calorimetry (ITC) and NMR^48, 50, 103^. For example, the *K*_D_ for p53-TAD^WT^-HDM2 interaction determined from BLI (260 ± 61 nM) is in agreement with the values of 230 ± 20 nM^50^ or 200 ± 20 nM^48^ from ITC experiments, despite slight differences in the length of the p53-TAD constructs and buffer conditions between these studies. Among the selected mutants, only the p53-TAD^K24N^-HDM2 interaction has been studied before and showed that the K24N substitution did not significantly affect the binding affinity^69^, again in agreement with the present BLI results (**Table 1**).

**Table 1.** Thermodynamic and kinetic parameters of interactions between p53-TAD variants and the binding partners. Each interaction was measured with two replicates at three unique concentrations of each binding partner using BLI with an Ni-NTA biosensor tip. Concentrations used were 2 µM, 3 µM, and 4 µM for HDM2 (residues 17-125), 20 µM, 30 µM, and 40 µM for TAZ1, and 300 nM, 600 nM, and 900 nM for TAZ2. *N*.*D*. indicates that the interaction was not detectable at this concentration range. The data indicating significant deviations from p53-TAD^WT^ are shown in italics.

| Binding Partner | p53 Variant | $K_D$ (nM) | $k_{on}$ ($10^4 \text{ M}^{-1}\text{s}^{-1}$) | $k_{off}$ ($10^{-3} \text{ s}^{-1}$) |
| --- | --- | --- | --- | --- |
| HDM2 (17-125) | WT | 260±61 | 7.7±1.3 | 19.2±1.1 |
| | K24N | 301±39 | 7.1±0.9 | 21.0±1.0 |
|  | *N29K/N30D* | *N.D.* | *N.D.* | *N.D.* |
| | D49Y | 275±52 | 8.0±1.1 | 21.6±2.6 |
| | W53G | 287±44 | 6.8±0.6 | 19.2±1.4 |
| TAZ1 | WT | 1639±380 | 1.45±0.30 | 22.8±1.9 |
| | K24N | 2159±470 | 1.52±0.21 | 32.0±4.4 |
| | N29K/N30D | 1650±430 | 1.35±0.24 | 21.3±1.8 |
| | D49Y | 2188±630 | 1.45±0.25 | 30.2±2.8 |
|  | *W53G* | *4953±1500* | *0.67±0.27* | *30.0±5.2* |
| TAZ2 | WT | 61.10±0.97 | 8.36±0.13 | 5.26±0.50 |
|  | *K24N* | *210±14* | *4.53±0.31* | *9.5±1.6* |
|  | *N29K/N30D* | *N.D.* | *N.D.* | *N.D.* |
|  | *D49Y* | *N.D.* | *N.D.* | *N.D.* |
|  | *W53G* | *N.D.* | *N.D.* | *N.D.* |

**Figure 2.**
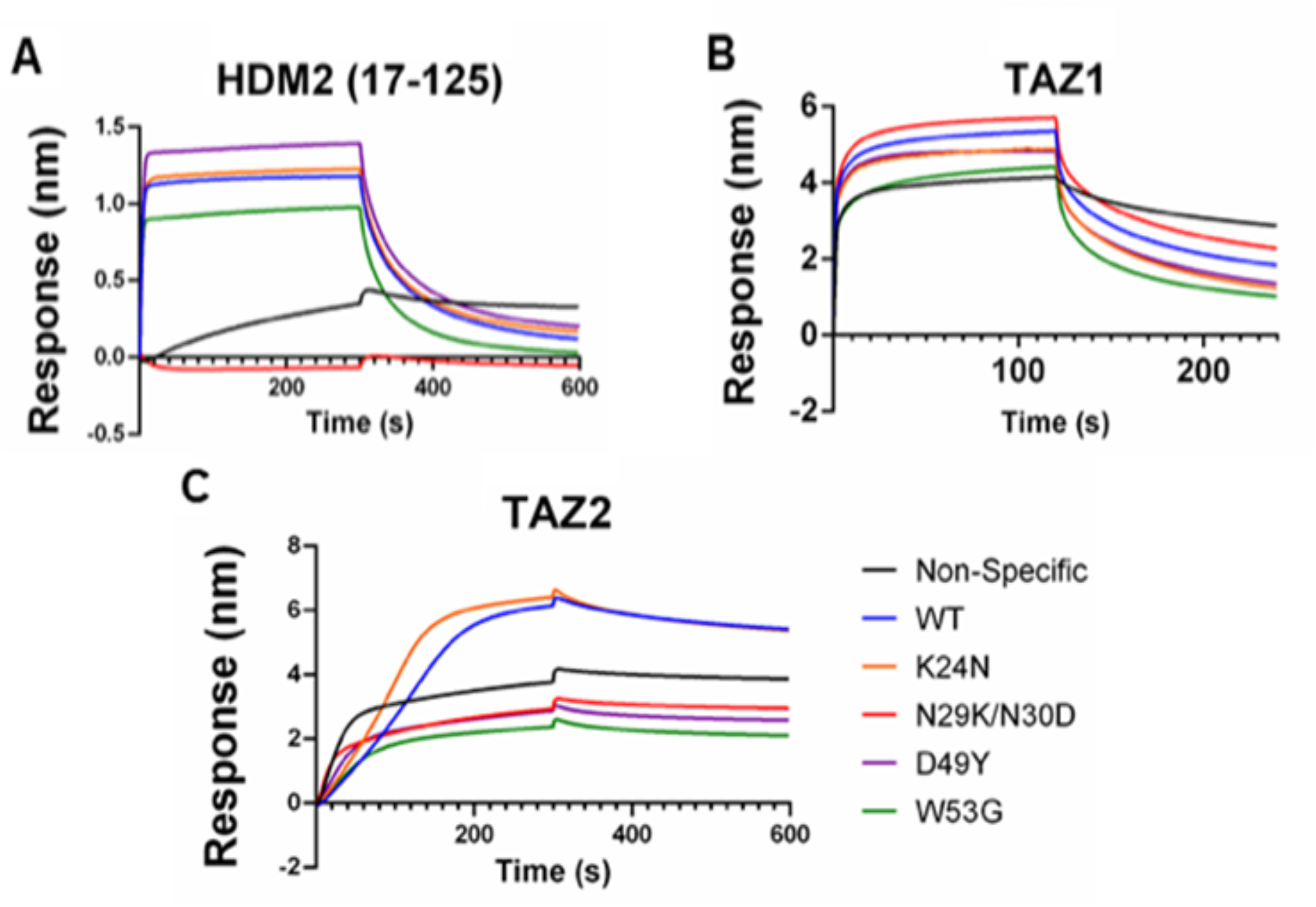
Binding of the p53-TAD variants to HDM2 (residues 17-125), TAZ1, and TAZ2. 100 ng/µL of p53-TAD^WT^ and its variants were applied to Ni-NTA BLI biosensor tip prior to the introduction of 3 µM HDM2(17-125) (A), 30 µM TAZ1 (B), or 300 nM TAZ2 (C). BLI signal response was set at 0 at the beginning of the association phase (time 0 s). The dissociation phase was initiated at 300 s for HDM2 and TAZ2 or at 120 s for TAZ1. Each trace represents the average response from two independent experiments. The control traces labeled “Non-Specific” were obtained without p53-TAD on the biosensor tip.

As summarized in **Table 1** and **Figure 2**, the four selected cancer-associated mutations in p53-TAD induce significant and diverse changes in the binding affinities towards the regulatory partners. Strikingly, the BLI response of HDM2 applied to a biosensor tip with immobilized p53-TAD N29K/N30D falls below the “non-specific” trace (Figure 2A), which indicates a severe loss of affinity of HDM2 towards the N29K/N30D variant. Similarly, the BLI traces for three p53-TAD variants (N29K/N30D, D49Y, and W53G) fall below the “non-specific” signal of TAZ2 binding (Figure 2C), again indicating a severe loss of interaction affinity. The above cases of undetectable interactions were labeled N.D. in Table 1.

Among the interactions that were detected and analyzed, the K24N substitution in p53-TAD leads to a significant reduction of binding affinity for TAZ2, as compared to p53-TAD^WT^ (Table 1). However, the K24N substitution does not affect interactions with TAZ1 or HDM2. Interestingly, N29K/N30D abolishes p53-TAD binding to both HDM2 and TAZ2 without affecting interactions with TAZ1. Both D49Y and W53G lose affinity towards TAZ2, but not towards HDM2. In addition, W53G, but not D49Y, shows a reduced the affinity towards TAZ1. Taken together, these results suggest that mutations in the disordered p53-TAD could significantly perturb the regulatory network of p53, thus likely disrupting its tumor suppressor function.

### Effects of cancer mutations cannot be fully explained by disruption of interfacial interactions

The observed effects of cancer mutations on p53-TAD’s interactions cannot be fully explained by perturbation of interfacial interactions^31, 104-106^. For example, HDM2 binds p53-TAD primarily through the short AD1 helix (residues 19-25)^31^ (**Figure S4**) and yet the N29K/N30D substitutions in the disordered region distinct from AD1 renders p53-TAD’s interaction with HDM2 undetectable (**Table 1**). In the most striking case of the p53-TAD interaction with TAZ2, where all four mutations significantly affect the binding affinity (Table 1), only Trp53 is involved in direct interactions of the complex (**Figure 3**). Lys24, Asn29/30 and Asp49 are all fully solvent-exposed in the p53:TAZ2 complex and are not involved in supporting protein-protein interactions. The Asn29/30 tandem in particular is positioned within a dynamic and disordered loop between the AD1 and AD2 regions that harbor the interaction sites for HDM2 and CBP (**Figure S4**).

**Figure 3.**
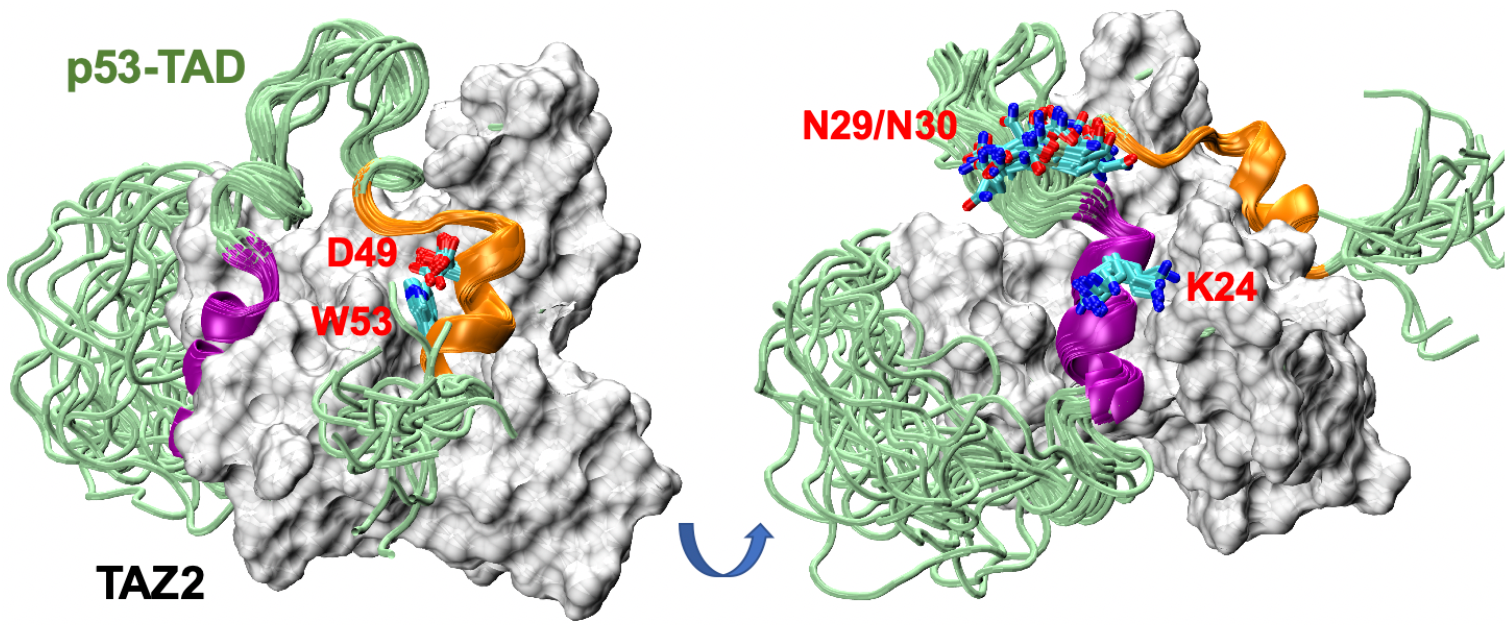
Dynamic binding interface between p53-TAD and TAZ2. All 20 members of the NMR ensemble (PDB: 5HPD^49^) are shown for p53-TAD (green, with AD1 and AD2 colored in purple and orange, respectively), while a single copy of TAZ2 (model 1 of the ensemble) is shown as silver molecular surface. Side chains of p53-TAD residues K24, N29, N30, D49 and W53 are shown in stick representation.

Therefore, the selected mutations likely affect the interaction affinities indirectly, possibly by modulating the unbound conformational ensemble of p53-TAD.

### NMR analysis of local and distal effects of cancer mutations on TAD structure and dynamics

We characterized the effects of mutations on the structure and dynamics of unbound p53-TAD using NMR spectroscopy. The ^15^N-HSQC spectrum of p53-TAD^WT^ was acquired under experimental conditions identical to those used previously by Vise et al^107^ (see **Materials and Methods**). Using the published assignments as the reference, all 60 non-proline residues of p53-TAD^WT^ and p53-TAD^K24N 15^N-HSQC spectra were then assigned to an accuracy within 0.01ppm of the previously reported values (**Figures S5-6**). Additionally, the ^15^N-HSQC spectra of p53-TAD^N29K/N30D^, p53-TAD^D49Y^, and p53-TAD^W53G^ were then acquired and assigned (**Figures S7-S9**). Orphaned cross-peak positions from each spectrum were assigned to the nearest orphaned WT assignment (**Tables S1-S5**). An overlay of all acquired ^15^N-HSQC spectra (**Figure 4A**) for the p53-TAD^WT^ and the four variants reveals that changes in the chemical environment induced by the mutations are observed for fewer than 15% of the observable peaks. Chemical shift perturbation (CSP) analysis of ^1^H and ^15^N chemical shift assignments of the mutants against their WT counterparts was calculated using a rescaled weighted distance function √(25(ΔH)^2^ + (Δ^15^N)^2^) for each residue^108^. The results, summarized in **Figure 4B**, show that CSPs are largely localized near the mutation sites for all four mutants. The lack of distal CSPs implies that the overall structure of the protein is only marginally impacted, especially on the secondary structure level.

**Figure 4.**
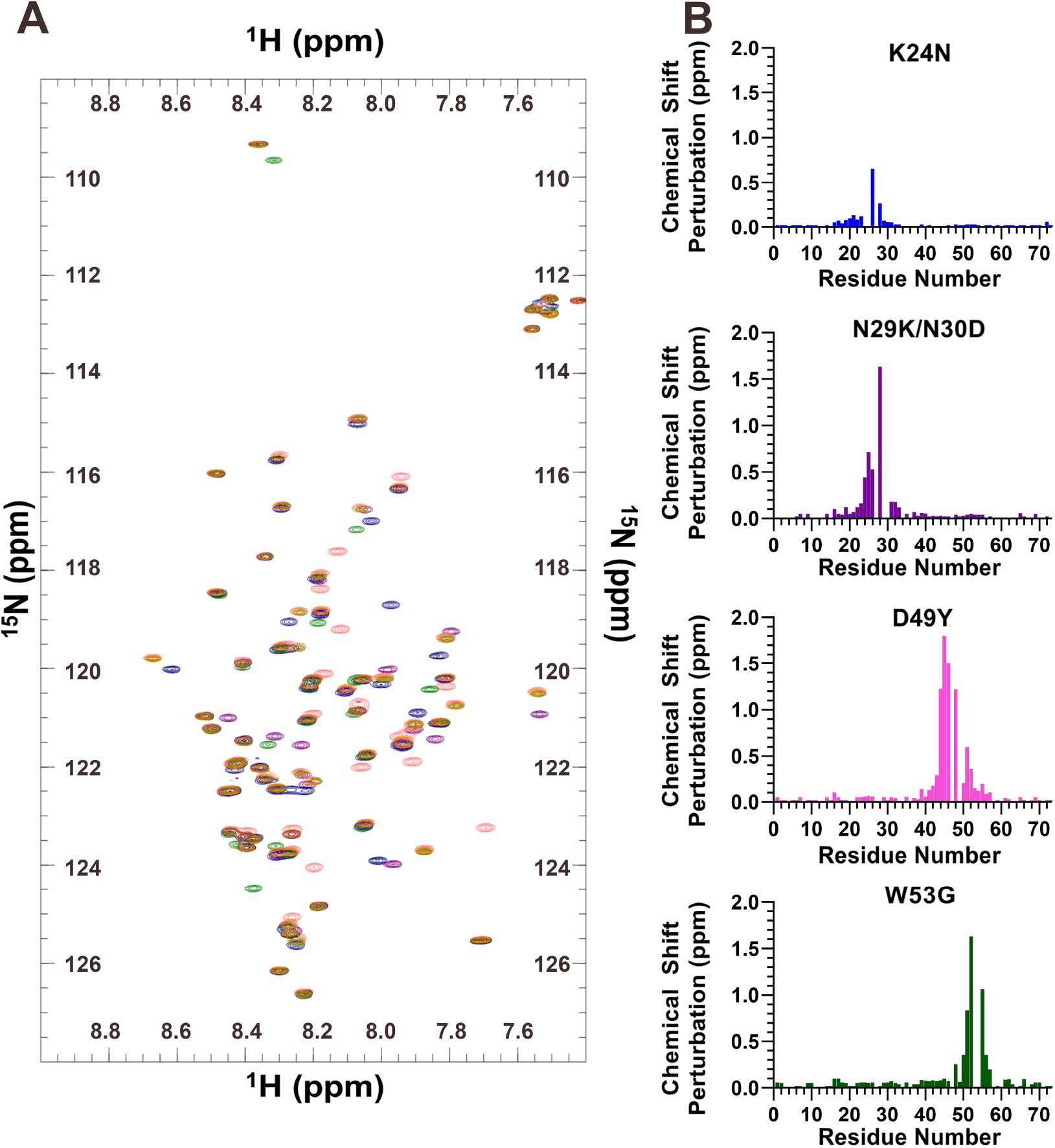
Chemical shift perturbations of p53-TAD variants. (A) Overlay of p53-TAD ^15^N-HSQC spectra for WT (orange), K24N (blue), N29K/N30D (purple), D49Y (pink), and W53G (green). Non-overlapped positions indicate changes in backbone amide chemical environment due to a mutation. (B) Chemical shift perturbation of each assigned p53-TAD residue referenced to their original WT assignment. A stronger chemical shift perturbation indicates a greater impact upon a residue’s chemical environment due to mutation.

A lack of detectable changes in the ensemble-averaged measurements of an IDP does not exclude the possibility of substantial conformational redistributions within the disordered ensemble^14^. For instance, it has been shown that phosphorylation of KID domain of transcription factor CREB at site Ser133 only leads to marginal changes in the overall helicity, even though the population of the underlying conformational substates is significantly perturbed and the accessible conformational space becomes more restricted due to phosphorylation^57^. Redistribution of conformational substates could lead to different relaxation profiles on both short (ps-ns) and long (µs-ms) timescales, which may be qualitatively correlated with a local and/or a long-range structural dynamics. Thus, to detect the effects of mutations on p53-TAD conformational distribution and dynamics, we measured NMR relaxation parameters including spin-lattice relaxation rates (R_1_) (**Tables S6-S10**), spin-spin relaxation rates (R_2_) (**Tables S11-S15**), and hetero-nuclear NOEs (NHNOE/NONOE ratios) (**Tables S16-S20**).

The measured NHNOE/NONOE ratio profiles are negative for all variants (**Figure 5A)**, suggesting that all mutants remain similarly disordered at the secondary structure level. This is consistent with the apparent lack of distal CSPs (**Figure 4B**). Nonetheless, there is a systematic decrease in the absolute values of the mean NHNOE/NONOE ratio for p53-TAD^K24N^ (−0.24±0.21) and p53-TAD^N29K/N30D^ (−0.21±0.23) compared to that for p53-TAD^WT^ (−0.47±0.27), which reflects a reduced flexibility or increased rigidity of the backbone. In contrast, W53G shows a significant reduction of the NHNOE/NONOE ratio in the C-terminal half of the protein, consistent with the expectation that replacing a large hydrophobic residue (W53) with Glycine should promote the backbone mobility. Importantly, analysis of the R_2_/R_1_ ratios reveals that there is a systematic decrease for both p53-TAD^K24N^ and p53-TAD^N29K/N30D^ compared to the WT protein, while there is a systematic increase for p53-TAD^W53G^ and to a lesser degree p53-TAD^D49Y^ (**Figure S10**). That is, p53-TAD^K24N^ and p53-TAD^N29K/N30D^ both have higher tumbling rates compared to the WT protein, suggesting a mutation-induced molecular compaction. In contrast, p53-TAD^W53G^ is more flexible and likely also slightly expanded.

**Figure 5.**
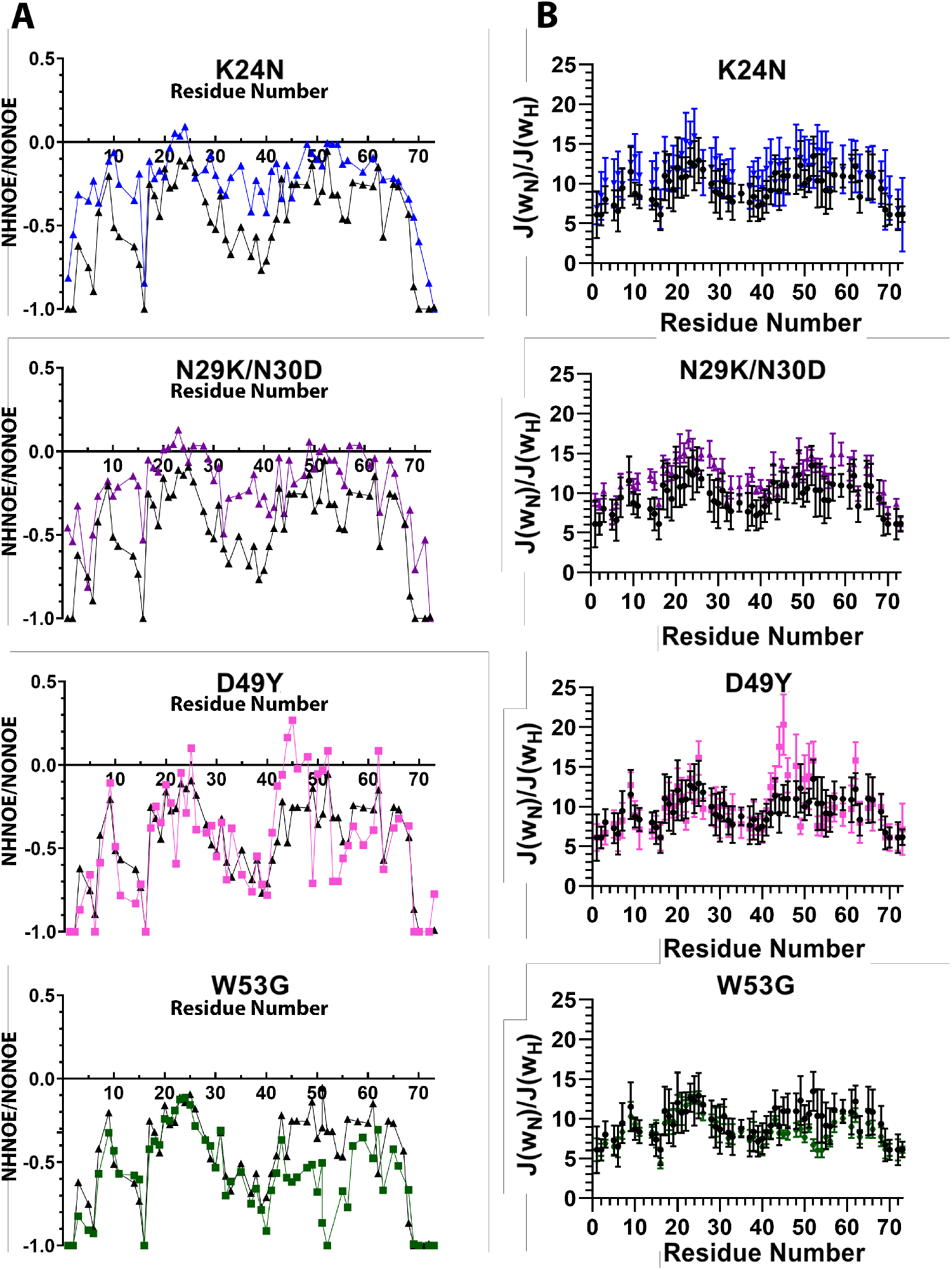
Effects of p53-TAD variants on dynamic properties. **The NHNOE/NONOE Ratio (A) and *J(ω***_**N**_***)/J(ω***_**H**_**) ratio (B)** for each p53-TAD variant was calculated on per residue basis with K24N (blue), N29K/N30D (purple), D49Y (pink) and W53G (green). For each p53-TAD variant, an overlay of the WT data (black) is included as the reference. Error bars represent 95% confidence interval. Higher *J(ω*_N_*)/J(ω*_H_) and NHNOE/NONOE ratios indicate a greater steric restriction, while lower values for both correlate with a greater flexibility.

To further interpret the changes in ensemble dynamics on a per residue basis, reduced spectral density approaches (RSDA)^109-111^ were used to approximate a relaxation power at 0 MHz (*J(0)*), 50.7 MHz (*J(ω*_N_)), and 435 MHz *(J(ω*_H_)) for each p53-TAD construct (**Figures S11-13, and Table S21**). Generally, the spectral density above 200 MHz using RSDA is sensitive to changes in auto-correlation times that reflect contribution from the amide ^1^H bond vector autocorrelation (<500 ps), while the spectral density below 200 MHz is dominated by the global molecular tumbling (>500ps), or the molecular correlation time (*τ*_c_). Consistent with the R_2_/R_1_ ratio analysis, *J*(0) is significantly reduced for both p53-TAD^K24N^ and p53-TAD^N29K/N30D^, to ∼1.45 ns/rad vs ∼ 1.85 ns/rad for p53-TAD^WT^, while remaining similar for the other two mutants. In rigid, uniformly tumbling proteins, the RSDA approximations of *J(0), J(ω*_N_), and *J(ω*_H_) would show agreement in the component *τ*_c_ values for all functions independent of frequency. However, all p53-TAD constructs suggest disparate *τ*_c_ values that vary from one another in a frequency-dependent manner, which suggests that numerous ps-ns motions contribute to the relaxation of the p53-TAD variants. To further explore this effect, we calculated the *J(ω*_N_*)/J(ω*_H_) ratio to evaluate changes in overall contributions from fast internal motions (*J(ω*_*H*_); *τ*_c_ < 500ps; NHNOE/NONOE ratio and R_1_ dependent) and compared them against contributions from segmental motion (*J(ω*_*N*_); *τ*_c >_ 500ps; predominantly R_1_ dependent)^109^. Disproportionate contributions from greater internal motions <500ps timescales will be reflected as lower *J(ω*_N_)/*J(ω*_H_) ratios for specific residue positions, while larger ratios correspond to lower contributions from faster timescale motions. As summarized in **Figure 5B**, we observed that p53-TAD^D49Y^ and p53-TAD^W53G^ display nearly identical *J(ω*_N_)/*J(ω*_H_) ratios as the p53-TAD^WT^, except near their respective mutation sites. In contrast, p53-TAD^K24N^ and p53-TAD^N29K/N30D^ steric restriction profiles are globally elevated with respect to p53-TAD^WT^, in agreement with the restricted flexibility for these protein variants indicated by the NHNOE/NONOE ratios (**Figure 5A**).

### Atomistic simulations of conformational redistributions in p53-TAD mutants

The above NMR analyses suggest that cancer-linked mutations may perturb the disordered ensemble of p53-TAD without significant changes of the local secondary structure propensities. Atomistic simulations were performed to further characterize the modulation of the disordered ensembles of p53-TAD. We recently demonstrated that structural ensembles generated using a combination of enhanced sampling via replica exchange with solute tempering (REST2)^91, 97^, highly accurate protein force field a99SB-disp^74^ and GPU-accelerated MD could accurately recapitulate a wide-range of structural features of large, non-trivial IDPs^112^. In particular, the overall size, secondary structure, and transient long-range contacts derived from the simulated ensemble of p53-TAD^WT^ were in quantitative agreement with a wide range of experimental data, including NMR chemical shifts, paramagnetic relaxation enhancement (PRE), SAXS and smFRET. The same protocol was applied here to generate atomistic ensembles of the four selected p53-TAD cancer-associated variants (see **Materials and Methods**).

We first analyzed key conformational properties to examine the convergence of the simulated ensembles, including radii of gyration (*R*_g_) (**Figure S14**), end-to-end distances (**Figure S15**) and overall helicity (**Figure S16**). Clearly, all p53-TAD variants are highly dynamic and sample numerous conformational states throughout the simulation timescales. By dividing each simulation trajectory into multiple segments, we observed that both the overall dimensions and secondary structures were highly consistent among the segments for all systems (**Figures S17-18**), indicating that all simulated ensembles were well-converged. For example, the residue helicity profiles are converged to within 0.05 for most regions (except around the residues 20-25 for p53-TAD^N29K/N30D^). We note that achieving the level of convergence as demonstrated here is very difficult for a complex IDP like p53-TAD, and this is often a major bottleneck in studies of IDPs using atomistic simulations in explicit solvent. Together with previous benchmark studies showing that a99SB-disp can faithfully describe both local and long-range structural properties of p53-TAD^74, 112^, the simulated ensembles are expected to provide a viable atomistic description of how cancer-associated mutations modulate the disordered state of p53-TAD.

Simulated atomistic ensembles recapitulate the observation from NMR chemical shift analysis (e.g., **Figure 4**) that the four cancer-linked mutations do not substantially perturb local structures of p53-TAD on the ensemble level (**Figure 6A**). Most structural differences between the mutated variants and wild type p53-TAD are localized near the mutation site. For example, p53-TAD^K24N^ slightly reduces the helical propensities near AD1 region. This is qualitatively consistent with a previous NMR secondary chemical shift analysis^69^, even though the helicity reduction observed in the current simulation is smaller (∼30% vs ∼50% reduction). Intriguingly, atomistic ensembles reveal possible distal effects on the residue helicity for some of the protein variants, particularly, p53-TAD^N29K/N30D^, where the residue helicity in the C-terminal AD2 region (residues 45-55) is significantly reduced. Distal effects of the mutations are indicative of transient long-range interactions in the disordered state of p53-TAD, which have been detected by both florescence quenching^72^ and paramagnetic relaxation enhancement NMR experiments^70^.

**Figure 6.**
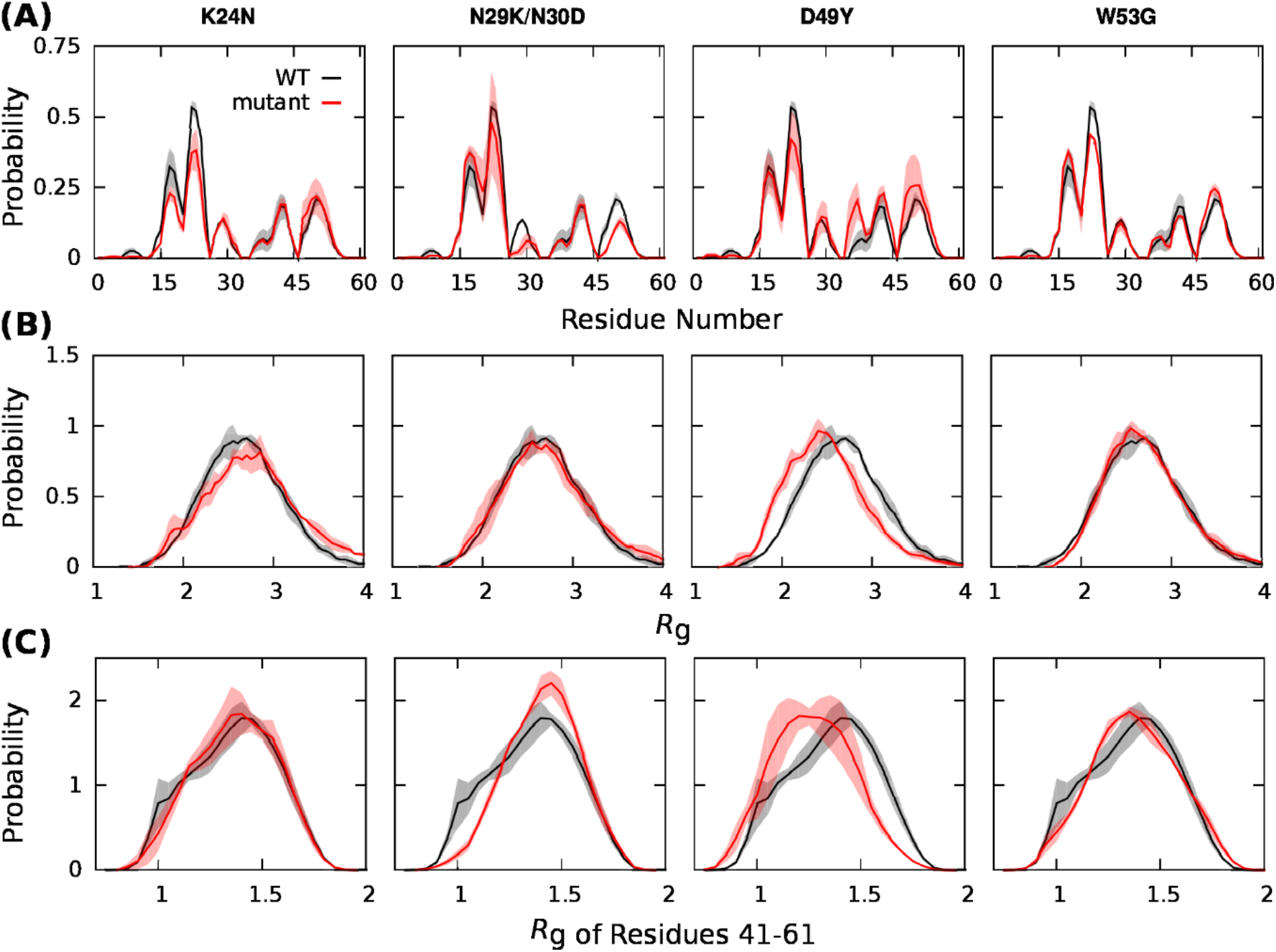
Conformational properties of p53-TAD variants obtained from atomistic simulations. (A) Residue helical propensity of four p53-TAD mutated variants in comparison with WT. Both α helices and 3_10_ helices were included in this calculation, since the probability of forming 3_10_ helices is nonnegligible in force field a99SB-disp^104^. (B) Radius of gyration (*R*_g_) probability distribution of four p53-TAD variants in comparison with WT. (C) Probability distribution of *R*_g_ of p53-TAD residues 41-61 for four protein variants in comparison with WT. The shaded areas indicate uncertainties, which were estimated by dividing each trajectory into three equal segments and calculating the standard deviation of the mean values.

Similarly, p53-TAD^D49Y^ showed little effects on helicity, except for the residues 35-40, where it was slightly increased. Although helical propensity near the mutated residue did not change significantly, we noted that the probability of forming helical turns (definition based on DSSP^113^, *i*.*e*., *i* → *i*+4 hydrogen bond formation) for residues 45-50 was notably increased (**Figure S19**). This observation was consistent with the NMR result that in p53-TAD^D49Y^ the mutation led to more steric restrictions for the same region based on the NHNOE/NONOE and *J(ω*_*N*_)/*J(ω*_*H*_) analysis (**Figure 5**). As for p53-TAD^W53G^, residue helicity did not display significant changes, even though the backbone of residue 53 was more flexible in p53-TAD^W53G^ than in WT (**Figure S20**). The latter agrees with the less restricted motions near the mutation site observed in NHNOE/NONOE and *J(ω*_*N*_)/*J(ω*_*H*_) analysis (**Figure 5**). The overall size of all p53-TAD variants is similar to that of WT, except D49Y, for which the *R*_g_ distribution slightly shifted towards a more compact state (**Figure 6B**). Taken together, the four cancer-associated mutations perturb the unbound ensemble of p53-TAD locally and distally, inducing subtle but non-negligible changes in both the overall dimensions and secondary structure of the protein.

To further analyze the conformational space available to p53-TAD and potential changes induced by the four cancer-associated mutations, we visualized the simulated ensembles through principal component analysis (PCA) (see Materials and Methods for detail). As shown in **Figure 7**, p53-TAD^WT^ mainly samples a broad basin of largely extended conformations, which is shared by all four mutated variants. However, the mutated variants populate additional states with significant probabilities except p53-TAD^W53G^. Analysis of the conformational properties reveal that these new basins mostly correspond to states of increased compaction and/or apparent rigidity. For instance, p53-TAD^D49Y^ shows a slight increase of the population of compact states, as illustrated in **Figure 7**. Through PCA analysis, we found that the p53-TAD^D49Y^ compaction could occur at either N-terminal or C-terminal region of the polypeptide, as confirmed by comparing *R*_g_ of the two regions (**Figures 6C and S21**) and contact probability maps (**Figure S22**) between p53-TAD^D49Y^ and p53-TAD^WT^. Both p53-TAD^K24N^ and p53-TAD^N29K/N30D^ also give rise to more compact conformations (**Figure 7)**, particularly in the N-terminal region **(Figures S21-22**). For p53-TAD^W53G^, the mutation leads to a wider basin of largely extended conformations similar to that of p53-TAD^WT^.

**Figure 7.**
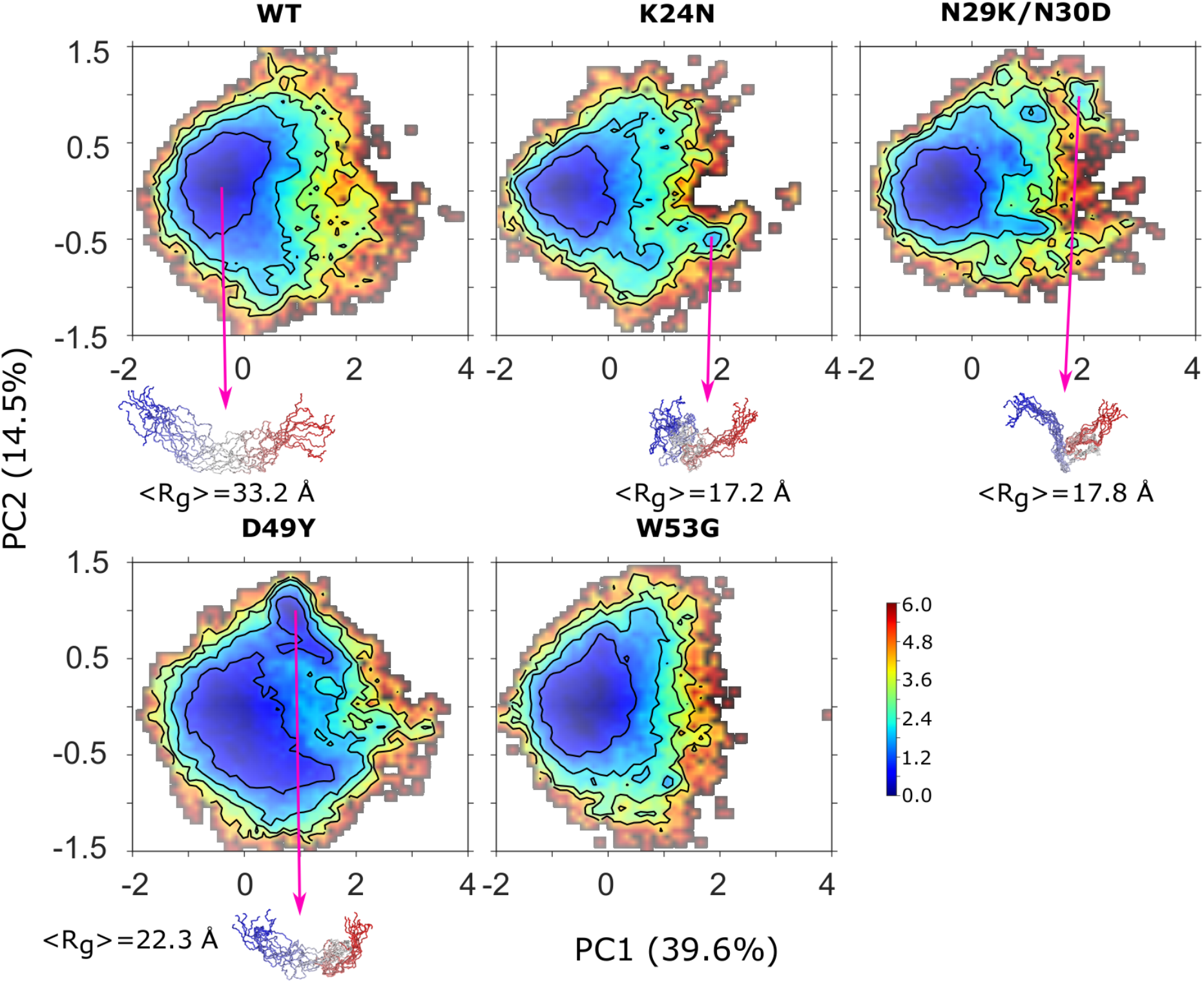
Conformational distributions of p53-TAD^WT^ and its variants. The simulated structural ensembles of p53-TAD were projected onto the first two PCs derived from PCA analysis. Heatmaps show the negative logarithm of the probability distribution. Values in the parenthesis are percentages of variance captured in each direction. Ten representative structures are shown for a few highly populated states, where the peptide is shown in backbone trace, with the color changing from red at the N-terminus to blue at the C-terminus. The average radius of gyration is also shown for each basin illustrated.

## Discussion

We discovered that cancer-associated mutations in intrinsically disordered p53-TAD can disrupt the balance of p53’s interaction with key activation and degradation regulator proteins. Notably, all four mutations selected for the current study significantly decrease p53-TAD’s affinity towards TAZ2, the CBP domain with the strongest binding affinity to WT p53. For three of the mutated variants, N29K/N30D, D49Y, and W53G, the interaction with TAD-TAZ2 was below the limit of detection (see **Figure 2** and **Table 1**), while for K24N the binding affinity was reduced more than three-fold. In contrast, only one mutated variant, N29K/N30D showed a defect in binding to HDM2 and only W53G showed a weaker binding to TAZ1, as compared to p53-TAD^WT^. Biologically active p53 forms a tetramer whose four TADs can simultaneously interact with four CBP domains, namely, TAZ1, TAZ2, KIX, and NCBD^114^. The K_D_ value for the p53-TAD interaction with TAZ2 is at least 30-fold lower than that with TAZ1 (see **Table 1**) and over 400-fold lower than those for binding KIX and NCBD^103^. Thus, among the four CBP domains, the dominant interaction, which regulates the status of the p53 activation, is provided by the TAD-TAZ2 interface. Our results indicate that all four selected cancer-linked variants of p53-TAD may be defective in their interactions with CBP through TAZ2, with N29K/N30D, D49Y, and W53G showing the strongest binding deficiencies. A loss of interaction with CBP implies a likely inhibition of the mechanisms that stabilize and activate p53, which could lead to a loss of the pro-apoptotic tumor suppressor activity and, as a result, contribute to tumor development. Interestingly, the N29K/N30D variant of p53-TAD apparently lost affinity towards both HDM2 and CBP (see **Table 1**). On the one hand, the N29K/N30D targeting for proteasomal degradation could be suppressed due to a less efficient ubiquitination by HDM2. However, even if the cellular stability of the N29K/N30D variant is enhanced, as compared to WT p53, the simultaneous CBP interaction defect could produce a dominant tumor-suppressor loss-of-function phenotype. Altogether, our results help explain why the selected mutations in p53-TAD are associated with cancer occurrence.

Importantly, the localization and orientation of the selected residues in p53-TAD suggests that their mutations do not directly perturb the binding interface of p53-TAD with the affected partners (**Figures 3 and S4**), with the exception of W53G in the complex with TAZ2. Instead, the mutations likely affect the interaction affinities indirectly, possibly by modulating the unbound conformational ensemble of p53-TAD. The defects in binding affinities caused by most of the selected mutations were severe, which precluded accurate determination of the kinetic and thermodynamic parameters for those interactions. However, the BLI data for the K24N interaction with TAZ2 indicate that the loss of binding affinity is due to a reduced association rate and an increased dissociation rate (Table 1). It has been shown previously that the conformational flexibility and residual structures play key roles in IDP association kinetics^115-116^, arguably by allowing rapid folding upon a nonspecific encounter with the target^117-119^. Thus, a reduced association rate for the p53-TAD mutants may result from modified populations of specific sub-states and/or the overall conformational rigidity of the disordered ensemble of unbound p53-TAD.

To characterize the p53-TAD conformational ensembles for WT and the mutated variants, we employed NMR spectroscopy and atomistic simulations. Both these approaches indicate that the selected mutations induce only subtle changes in the ensemble-averaged protein properties, such as secondary structure propensity (Figures 4 and 6). However, a detailed analysis of the NMR results points at systematic mutation-induced changes in the p53-TAD conformational dynamics. Specifically, the selected mutations in the N-terminal segment of TAD (K24N, N29K/N30D) produce a small, but significant compaction of the domain, as shown by a systematic decrease in R2/R1 relaxation parameter (Figure S10), a decrease in the absolute values of NHNOE/NONOE ratio (Figure 5A), and the changes in the spectral density parameters (Figures 5B and S11). In contrast, the mutations in the C-terminal segment of the domain (D49Y, W53G) produce a slight expansion of the polypeptide based on an increase in R2/R1. The NMR results imply mutation-induced changes in the population and/or stability of specific sub-states within the disordered conformational ensemble of p53-TAD for all selected mutations. Interestingly, either an apparent compaction or an expansion of the disordered TAD has pronounced effects in its interactions with the protein partners (Table 1).

The atomistic simulations did not resolve the full details of the mutation-induced effects of the p53-TAD dynamics, as determined by NMR. For example, an apparent compaction of the domain was detected in the simulation of D49Y on the ensemble level, but not for the remaining mutated variants (Figure 6B). Detailed analysis of the conformational space reveals that both K24N and N29K/N30D also lead to increased populations of compacted substates (Figure 7), which can partially explain the results of NMR relaxation analysis. The limitation in atomistic simulations to fully resolve fine features of the disordered ensemble of an IDP reflects the remaining deficiencies in both the protein force field and achievable conformational sampling. We note that the a99SB-disp^74^ force field used in this work is considered one of the most suitable for IDP simulations and that an aggregated enhanced sampling time of over 240 µs is, by far, the most extensive for an IDP of p53-TAD’s size and complexity.

Our observation that even subtle modulation of the disordered conformational ensemble could lead to dramatic changes in binding to key regulators is intriguing. It may reflect a fundamental principle of how IDPs perform versatile functions in biology, that is, the disordered ensemble of IDPs is poised to respond sensitively and rapidly to various cellular stimuli, thus acting as central conduit in cellular signaling and regulation. It further suggests a new conceptual framework to guide the design of novel therapeutic strategies targeting IDPs, where small molecules could modulate the structural ensembles of IDPs through dynamic interactions with an ensemble instead of site-specific ones to alter the interaction profiles and control physiological functions.

## Supporting information

Supporting Information (all figures and tables)

## Acknowledgements

This work was supported by the National Institutes of Health (GM114300). The computing was performed on the Pikes cluster housed in the Massachusetts Green High-Performance Computing Center (MGHPCC).

## Materials and Methods

### Expression, mutagenesis and purification of p53-TAD (1-73)

A pET28a vector with sub-cloned cDNA of *Hs*p53-(1-73) (72R) and the N-terminal His-tag was procured from Addgene, (plasmid #62082), deposition courtesy of Dr. Gary Daughdrill (University of South Florida). p53-(1-73) plasmid was then transformed into New England Biolabs DH10β cells for plasmid stock generation in preparation of mutagenesis. p53/pET28a plasmid was then templated in Quickchange Lightning Site-Directed mutagenesis kit (Agilent Technologies) to create the desired mutations in p53-(1-73) through primer mismatch:

K24N (FP: 5’-TCAGAVCTATGGAATCTACTTCCT-3’; RP: 5’-AGGAAGTAGATTCCATAGGTCTGA-3’), N29K/N30D (FP: 5’-GGAAACTACTTCCTGAAAAAGACGTTCTGTCCCCCTTGCC-3’; RP: 5’-GGCAAGGGGGACAGAACGTCTTTTTCAGGAAGTAGTTTCC-3’), D49Y (FP: 5’-CTGTCCCCGGACTATATTGAA-3’; RP: 5’-TTCAATATCGTCCGGGGACAG-3’), and W53G (FP: 5’-CGGACGATATTGAACAAGGCTTCACTGAAGACCCAGGTCC-3’; RP: 5’-GGACCTGGGTCTTCAGTGAAGCCTTGTTCAATATCGTCCG-3’)

P53-(1-73) variants were then transformed into BL21(DE3) low background strain (LOBSTR, Kerafast, Boston, Massachusetts) chemically competent cells and selected for with LB agar plates with 50 µg/mL Kanamycin and 1%(w/v) glucose. Expression of p53-(1-73) for Biolayer Interferometry (BLI) was accomplished using LB media (**e**.**g**., **Figure S23**). p53-(1-73) variants expressed using LB media were grown on shakers at 30°C in 4L cultures of LB with Kanamycin and 1% glucose to an OD_600_= 0.5-0.7, at such point 1mM Isopropyl-β-D-thiogalactoside (IPTG, Sigma) was added to the culture and grown at 37°C for an additional 6 hrs. Purification of p53-(1-73) from the clarified cell lysate was accomplished using a Nickel Nitrilotriacetic Acid (Ni-NTA, Qiagen) column purification and 50 mM Tris-HCl, 300 mM NaCl at pH 8.0 with either 10 mM or 250 mM imidazole for wash and elution buffers respectively. Eluent fractions of p53-(1-73) variants were then concentrated using Amicon Ultra-15 (3.5 KDa MWCO) centrifugal filters. Concentrates were then subjected to two rounds of G75 Superdex Gel-filtration column chromatography at 4°C with 300 mM NaCl, 50 mM NaH_2_PO_4_, 1 mM EDTA at pH 7.0 buffer and centrifugal concentration. Final concentrates of p53-(1-73) variants were then dialyzed overnight at 4°C in a Slide-A-Lyzer dialysis cassette (3.5 KDa MWCO) into a 20 mM Tris-HCl, 50 mM NaCl, and 10% glycerol at pH 7.5 prior to storage at −20°C.

^15^N labeling of p53-(1-73) variants for NMR studies was grown in 6L cultures of M9 media (2.5 mg/L Thiamine, 2.5 mg/L Choline Chloride, 2.5 mg/L Ca-d-Pantothenate, 2.5 mg/L Nicotinamide Adenine Dinucleotide, 1.25 mg/L Pyridoxal Hydrochloride, 50 mg/L EDTA, 8.3 mg/L FeCl_3_ · 6H_2_O, 840 µg/L ZnCl_2_, 130 µg/L CuCl_2_·2H_2_O, 100 µg/L CoCl_2_ ·6H_2_O, 100 µg/L H_3_BO_3_, and 16 µg/L MnCl_2_ ·6H_2_O; 6 g/L Na_2_HPO_4_, 3 g/L KH_2_PO_4_, 500 mg/L NaCl, 4 g/L Glucose, 120 mg/L MgSO_4_, 33 mg/L CaCl_2_ and 500 mg/L ^15^NH_4_Cl) at 30°C. Induction of expression using 1mM IPTG was conducted OD_600_=0.5-0.7. The cultures were then incubated for an additional 6hrs at 37°C. Initial Ni-NTA of ^15^N labeled p53-(1-73) purification was identical to conditions outlined in LB expressions. Ni-NTA eluents of labelled p53-(1-73) were then concentrated and dialyzed into 50 mM Tris-HCl overnight. Cleavage of N-terminal 7His-tag was accomplished via recombinant thrombin cleavage site using Sigma-Aldrich Clean Cleave thrombin-agarose kit for 6 hrs cleavage at room temperature. Cleaved p53-(1-73) was then concentrated and purified using the Gel-filtration chromatography. Cleaved and gel-filtered p53-(1-73) eluents were passed over Ni-NTA column with flow through and wash fractions containing cleaved p53-(1-73) variants were collected. Finally, successfully purified ^15^N labeled p53-(1-73) variants were concentrated to a final volume of less than 1 ml prior to dialysis into the final NMR buffer of 50 mM NaH_2_PO_4_, 50 mM NaCl, 1 mM DTT, 1 mM EDTA, pH 6.5. A 675 µL volume containing an approximate 300 µM concentration of each p53-(1-73) variant with NMR acquisition buffer was mixed with 35 µL volume of 99% D_2_O to create 5%D_2_O/95%H_2_0. The final 700 µL volume was then loaded into a Wilmad Precision 5 mm probe tube via glass transfer pipette for NMR acquisition.

### Expression and purification of HDM2 (17-125)

A pGEX-6p-2 vector with subcloned cDNA of HDM2 (17-125) with a N-terminal GST tag was a gift from Dr. Gary Daughdrill, (Addgene #62063). The pGEX6P2-HDM2(17-125) plasmid was transformed into New England Biolabs DH10-β *E. coli* competent cells for plasmid stock preparation. The plasmid was then transformed into BL21 (DE3) Rosetta rare tRNA coding chemically competent cells and selected using 25 µg/mL Chloramphenicol and 100 µg/mL of Ampicillin on LB agar plates. HDM2 was expressed using LB media and grown at 37°C with Chloramphenicol and Ampicillin to OD_600_ = 0.8, at such point 1mM IPTG was added to the culture and grown at 25°C for an additional 5 hrs. Culture was then centrifuged at 5000 rpm for 30 mins and pellets were resuspended in Glutathione S-transferase (GST) binding buffer (50 mM Tris-HCl, 300 mM NaCl, 2.5 mM EDTA, 0.02% NaN_3_, 2 mM DTT, pH 7.4). The cells were lysed using French press and centrifuged at 13,000 rpm for 40 mins. HDM2 containing supernatant was loaded onto a Glutathione Sepharose 4B resin (GE Healthcare) column that had been already equilibrated with GST binding buffer. The HDM2 protein fractions were eluted with GST binding buffer with 10 mM reduced glutathione. The GST tag was removed using HRV 3C protease (Thermo Scientific) with an enzyme to substrate ratio of 1:100 at 4°C for 16 hrs. After the removal of the GST tag the HDM2 protein was concentrated using Amicon Ultra-15 (3.5 KDa MWCO) centrifugal filters. Finally, the protein was buffer exchanged into 20 mM Tris-HCl, 150 mM NaCl, 1mM DTT, pH 8.0 with 10% glycerol prior to storage at −20°C (see **Figure S24**).

### Expression and purification of TAZ2 domain of CBP

A pET22b recombinant plasmid coding for a bi-cistronic mRNA from the cDNA of 6His-Gb1-STAT1 (710-750) and CBP (1764-1850) TAZ2, which was provided as a gift from Dr. Peter Wright (Addgene plasmid #99342). The pET22b/STAT1/TAZ2 cDNA encodes a second ribosomal binding site at the start of the TAZ2 cDNA sequence enabling concurrent expression of TAZ2 and binding partner STAT1. pET22b/STAT1/TAZ2 was transformed into BL21(DE3) Rosetta rare tRNA coding chemically competent cells and selected for using 25 µg/mL Chloramphenicol and 100 µg/mL of Ampicillin on LB agar plates with 1% glucose.

STAT1/TAZ2 was expression was accomplished using 2L volume of LB with 1% glucose, Chloramphenicol and Ampicillin via shaker at 37°C. When the culture OD_600_=0.1-0.3 approximately 250 µM of ZnCl_2_ was introduced to facilitate stability of expressed TAZ2. Induction of culture expression with 1 mM IPTG was conducted when OD_600_=0.5-0.7 and then allowed to express for 3hrs. Following pelleting and cell lysis STAT1/TAZ2 was resuspended in 20 mM Tris-HCl, 300 mM NaCl and 10 mM Imidazole at pH 8.0. The clarified cell lysate was then passed over Ni-NTA column and elution fractions containing STAT1/TAZ2 were isolated. The STAT1/TAZ2 fractions were then applied to a sulfopropyl (SP) sepharose cation exchange column with a flow rate of 0.3 mL/min. Using 20 mM Tris-HCl and 10 mM DTT at pH 6.5 buffer, TAZ2 was independently eluted using a linear salt gradient from 50 mM to 1M NaCl. Purified TAZ2 fractions were then quickly concentrated and then dialyzed into 25 mM Na-Acetate, 25 mM NaCl, 0.1 mM DTT, and 0.1 mM ZnCl_2_ at pH=5.8. Appropriate folding of TAZ2 was affirmed through Circular Dichroism (CD) spectroscopy (**Figure S25**) prior to storage at 4°C.

### Expression and purification of TAZ1 domain of CBP

pET21a plasmid containing CBP-(340-439) TAZ1 was provided by Dr. Peter Wright’s lab. pET21a/TAZ1 was chemically transformed, selected, and expressed identically as outlined for STAT1/TAZ2 purification. TAZ1 clarified lysate was suspended in 20 mM Tris-HCl, 50 mM NaCl, and 10 mM DTT at pH 6.9 and was isolated using SP-sepharose linear gradient procedure outline for TAZ2. TAZ1 containing eluent was then concentrated and injected onto a G75 Superdex with 20 mM Tris-HCl, 300 mM NaCl, and 10 mM DTT at pH 6.8. Fractions containing TAZ1 were then concentrated and dialyzed into 20 mM Tris-HCl, 50 mM NaCl, and 1 mM DTT at pH 6.8. TAZ1 folding, as with TAZ2, was assessed using CD (**Figure S26**) prior to storage at 4°C.

### Biolayer interferometry binding assays

Protein-protein interaction experiments were performed with a BLItz biolayer interferometer (ForteBio; Freemont, California) at room temperature. p53-TAD (WT or each of the variants) was diluted into buffer K (20 mM Tris-HCl, 50 mM NaCl, 0.01% (W/V) BSA, and 0.002% (V/V) Tween-20) to a final concentration of 100 ng/µL. Aliquots of HDM2 (17-125), TAZ2, and TAZ1 were diluted into 20 mM Tris-HCl, 50 mM NaCl, and 1 mM DTT at pH 7.5 (6.8 for TAZ1) to appropriate target protein concentrations (HDM2 (17-125): 2 µM, 3µM, and 4 µM; TAZ1: 20 µM, 30 µM, and 40 µM; TAZ2: 300 nM, 600 nM and 900 nM). Ni-NTA BLItz biosensor tips were hydrated for 10 minutes in buffer K prior to the experiments. The data acquisition was conducted using BLItz Pro 1.2 software. The experiments included the following steps: baseline 1, biosensor tip was equilibrated for 30 s in buffer K; analyte binding, p53-TAD or buffer K (control) was applied to the tip for 180 s; baseline 2, the tip was washed with buffer K for 120 s; ligand association, HDM2 (17-125), TAZ1, or TAZ2 was applied to the tip for 300 s (120 s for TAZ1); ligand dissociation, the tip was washed in buffer K for 300 s (120 s for TAZ1). The concentration of p53-TAD used for immobilization on the biosensor tips was optimized to achieve the strongest response during the analyte binding stage with the signal remaining stable during a subsequent washing step, which indicated a specific immobilization of the His-tagged p53-TAD on the Ni-NTA tips. The ligand association and dissociation phases were analyzed with nonlinear least-squares fitting of the kinetic equations built in the GraphPad Prism 8.1.0 software to obtain the apparent values of the binding on-rate (*k*_on_), off-rate (*k*_off_), and the dissociation constant (*K*_D_).

### Heteronuclear NMR experiments of p53-TAD variants

All 2D heteronuclear ^15^N-^1^H NMR experiments were acquired on a 11.75 Tesla Varian 500 MHz VNMRs system (Agilent Technologies Inc., Palo Alto, CA) with an operational frequency of 499.84 MHz. All NMR experiments were accomplished at the NMR Core Facility in the Department of Biochemistry and Molecular Biophysics at Kansas State University, utilizing a 5 mm cryogenic inverse detection pulse field gradient probe operating at a temperature of 25*τ*0.1°C. Purified p53-TAD variants were prepared in 95% triple distilled H_2_O and 5% D_2_O NMR acquisition buffer to an effective concentration of 300 µM for initial ^15^N-HSQC acquisition (Bodenhausen, 1980). 2D ^15^N-Heteronuclear Single Quantum Coherence (^15^N-HSQC) experiments of p53-TAD (1-73) variants were acquired with a spectral width of 6009 Hz and 256 complex points t_2_ and data in the ^15^N dimension were acquired using a sweep width of 1944 Hz and 256 complex t_1_ points. 100% of all HSQC cross-peak resonance assignments for p53-TAD^WT^ (1-73) were assigned based upon reported chemical shift values from Vise et al, 2005. Assignments of p53-TAD (1-73) for K24N, N29K/N30D, D49Y, and W53G variants were assigned in the same manner as WT with missing cross-peaks being tentatively assigned to the most proximal orphan peaks. All NMR experiments were conducted with a pre-saturation pulse scheme for suppression of the solvent peak (HOD) at 4.78 ppm. The residual water peak was then used as an internal reference for chemical shift assignments^120^. The processing of NMR free induction decay data was conducted using VNMRJ3.2a (Agilent Technologies Inc., Palo Alto, CA). Spectral analysis was then conducted using graphical assignment utilities CCPNMR and Sparky-NMRFAM^121-122^.

### Relaxation data acquisition and analysis of p53-TAD variants

Relaxation experiments for uniformly ^15^N-labeled p53-TAD (1-73) variants were acquired at 298 K and a concentration of 0.5 mM in NMR buffer. Spin-lattice relaxation rates (R_1_), spin-spin relaxation rates (R_2_) and ^1^H-^15^N NOEs ratios were measured by inverse-detected 2D NMR methodologies^123^. Spin-lattice relaxation rates were measured by collected 10 2D spectra with relaxation delays of 10, 50, 110, 190, 310, 500, 650, 1000, 1500, and 1900 ms. Spin-spin relaxation rates were also measured by collecting 10 2D spectra with relaxation delays of 10, 30, 50, 90, 110, 150, 190, 210, 230 and 250 ms. Peak heights of each series of relaxation experiments were then fitted to a single decaying exponential function to determine their respective relaxation times. Measurement of ^1^H-^15^N NOEs was accomplished with 2 separate spectral acquisitions, one with a 3s mixing for NOE buildup (NHNOE) and a second with a 3s recycle delay as reference (NONOE). In general, fitting errors were within 10% of calculated relaxation rates.

Analysis of relaxation rates was integrated using a reduced spectral density mapping approach^107, 110-111, 124-125^. The influences upon ^15^N nuclei relaxation are largely contingent upon the ^15^N chemical shift anisotropy, c, and the dipolar coupling interaction, d, between amide ^15^N and bound ^1^H^126^. The values for R_1_, R_2_, and NOE between amide protons and nitrogen nuclei can then be related in terms of spectral density functions: 

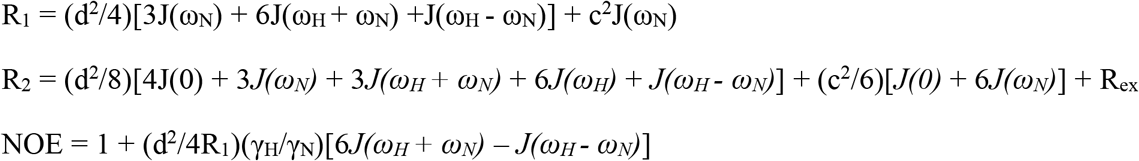

*J(ω)* is the power spectral density function that defines the reorientation of the ^1^H-^15^N bond in both stochastic (global) and intramolecular motions as a function of frequency. The reduced spectral density mapping approach assumes that linear combinations of *J(ω*_*H*_ *+ ω*_*N*_*), J(ω*_*H*_ *-ω*_*N*_), and *J(ω*_*H*_) are treated equally as an average value of *J(ω*_*H*_) (or *J(0*.*87ω*_*H*_)). Using this approximation, the aforementioned relationships can be rewritten in terms of *J(0), J(ω*_*H*_), and *J(ω*_*N*_): 

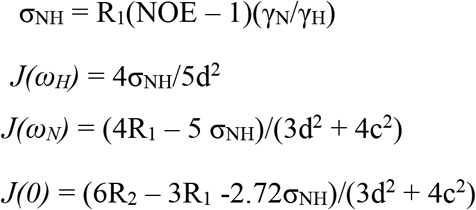

Where σ_NH_ is the spectral density, γ_N_ and γ_H_ are the gyromagnetic ratios for ^15^N and ^1^H nuclei, d = (µ_0_hγ_H_γ_N_/8π^2^)(r_NH-3_) is the dipole-dipole relaxation coefficient, µ_0_ is the magnetic permeability of a vacuum, h is Plank’s constant, r_NH_ is bond-length between amide nitrogen pairs (∼1.02Å), and c = ω_N_Δσ/√3 is the chemical shift relaxation coefficient, and Δσ is the chemical shift anisotropy for ^15^N which is (−160ppm). Using this approach, a single field acquisition of R_1_, R_2_, and HETNOE rates enables effective approximation *J(0), J(ω*_*H*_), and *J(ω*_*N*_) values.

### Atomistic simulation and analysis

p53-TAD^WT^ and four cancer-associated mutants (residues 1-61, see **Figure 1**) were simulated in this study. Both N- and C-termini of the peptide were capped, using an acetyl group and N-methyl amide, respectively. Force field a99SB-disp^74^ was used in all simulation, which has been shown to be one of the most accurate force fields in describing both local and long-range structural features of large IDPs^112^. For each system, the starting structure was generated by solvating a fully extended conformation in a truncated octahedron box using ∼24,000 water molecules. Also, proper number of counter ions (14 Na^+^ in all systems, except for 15 Na^+^ in K24N and 13 Na^+^ in D49Y) were added to neutralize each system. The final volume of simulation boxes was ∼710 nm^3^.

All simulations were carried out using GROMACS 2018^80, 127^ patched with PLUMED2.4.3^128^. Energy minimization using steepest descent algorithm was first performed to remove potential steric clashes in the system. Each system was then equilibrated for 100 ps at 298 K under constant temperature (NVT) conditions, with positions of protein heavy atoms in x, y, and z directions restrained by harmonic potentials. The force constant of restraining harmonic potentials was 1000 kJ/mol/nm2. A 1-ns NPT simulation was also performed at 298 K and 1 atm to further equilibrate the system, keeping the positions of protein heavy atoms restrained. Finally, the system was equilibrated under the same NPT conditions for 1 ns, during which all positional restraints on the peptide were removed. For the last equilibrium simulation, averaged volume of simulation box was computed, and the conformation whose volume was closest to the averaged volume was selected as the starting conformation of production run. All production simulations were performed at 298 K under NVT conditions using the replica exchange with solute tempering (REST2) technique^91, 97^. In comparison with temperature replica exchange, REST2 allowed for accelerated simulation convergence and reduced computational cost at the same time^91^. In REST2 simulations, only the region of interest (e.g., solute) is subject to effective tempering, which can be achieved by scaling the solute-solute and solute-solvent interactions by λ and 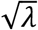
, respectively. In the present work, 16 replicas were used in each REST2 simulation. The effective temperatures of peptide in these 16 replicas ranged from 298 K to 500 K spaced exponentially, which corresponded to λ values of 1.00, 0.97, 0.93, 0.90, 0.87, 0.84, 0.81, 0.79, 0.76, 0.73, 0.71, 0.68, 0.66, 0.64, 0.62 and 0.60, respectively. Exchange attempt frequency between neighboring replicas was every 2 ps, and the mean acceptance ratio was ∼25%. In all simulations, van der Waals interactions were truncated at 1.2 nm, and long-range dispersion corrections were applied. Short-range electrostatic interactions were also cut off at 1.2 nm, with Particle Mesh Ewald (PME)^129^ applied to treat long-range electrostatic interactions. Lengths of bonds between heavy atoms and hydrogen atoms were constrained using LINCS algorithm^130^. Equation of motion was integrated with a time step of 2 fs, and the REST2 simulations lasted for 3.4, 2.2, 2.3, 4.2, and 3.0 µs/replica in WT, K24N, N29K/N30D, D49Y and W53G, respectively. To our best knowledge, this is probably one of the most extensive simulation studies of p53-TAD using explicit solvent all-atom force fields.

Structural ensembles of p53-TAD^WT^ and four cancer-associated mutants were constructed by collecting conformations at λ = 0 from REST2 simulations. Snapshots from the first 500 ns simulations were excluded since the systems were still under equilibration at this stage. All analyses were performed using GROMACS toolset^80, 127^ in combination with in-house scripts. To visualize conformational space available to p53-TAD as well as potential changes induced by cancer-associated mutations, PCA was performed on the peptide backbone conformations. Trajectories of p53-TAD^WT^ and four mutants were first combined, and distribution of reciprocal of interatomic distances (DRID)^131^ was computed for each snapshot using MSMBuilder^132^. Since it is well known that root-mean-square deviation (RMSD) in Cartesian coordinate space is not an optimal distance metric for large, disordered systems, we instead performed PCA analysis using DRID as input.

