## Supporting Information (all figures and tables) for "Cancer-Associated Mutations Perturb the Structure and Interactions of the Intrinsically Disordered p53 Transactivation Domain"

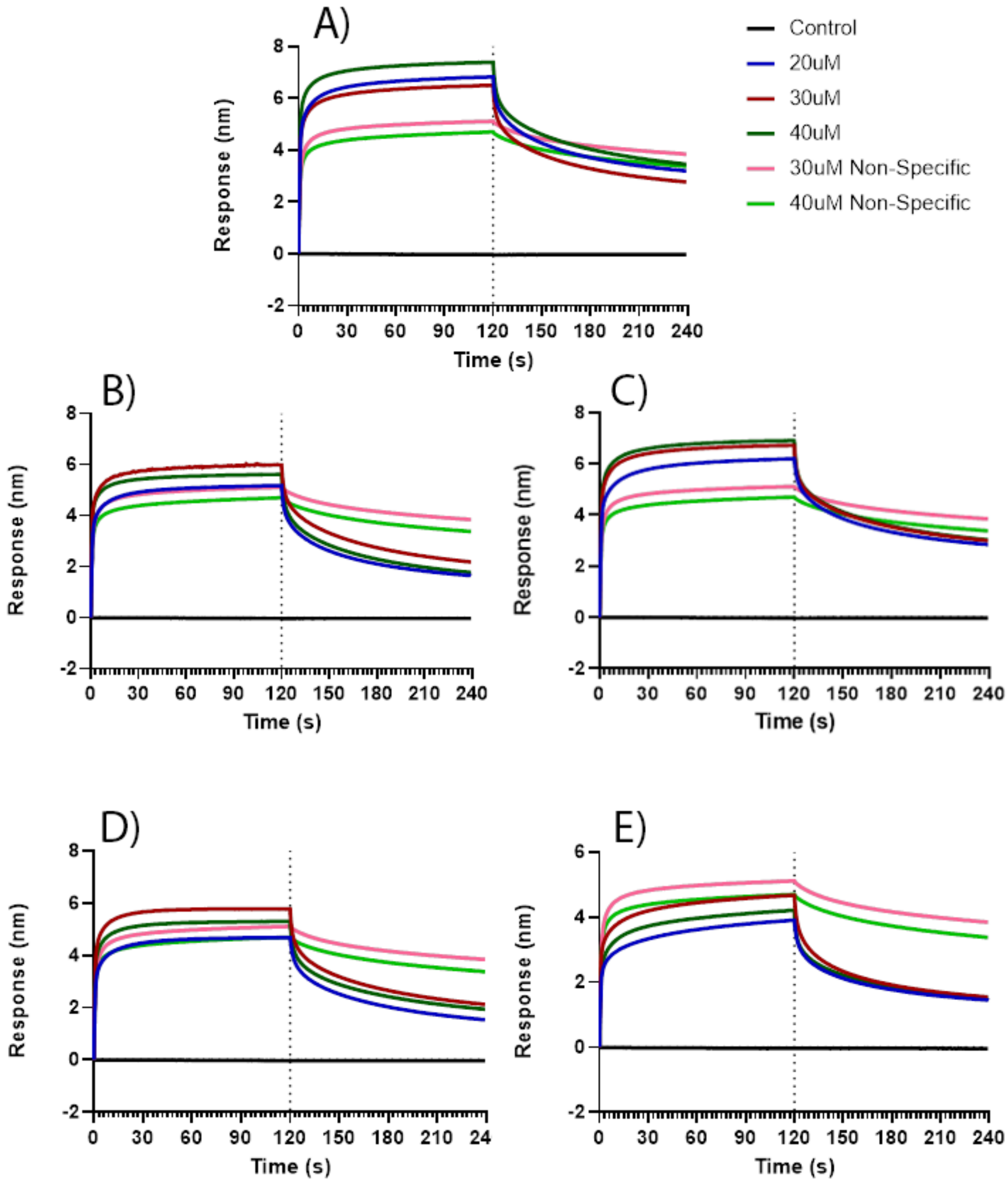


**Figure S1. TAZ1 binding to p53-TAD variants.** p53-TAD variants immobilized on the Ni-NTA biosensor tip (A, WT; B, K24N; C, N29K/N30D; D, D49Y; E, W53G) were subjected to 3 different concentrations of TAZ1, as indicated in the figure. Non-specific binding of TAZ1 to the Ni-NTA tip in the absence of p53-TAD was tested at 2 concentrations denoted as “Non-Specific.” The trace denoted as “Control” was obtained with each p53-TAD variant bound on the tip without including TAZ1 and demonstrates that p53-TAD variants were not depleted during BLI runs. Vertical line at 120 s denotes transition from the association phase to the dissociation phase. The traces shown are averages from 2 independent experiments.

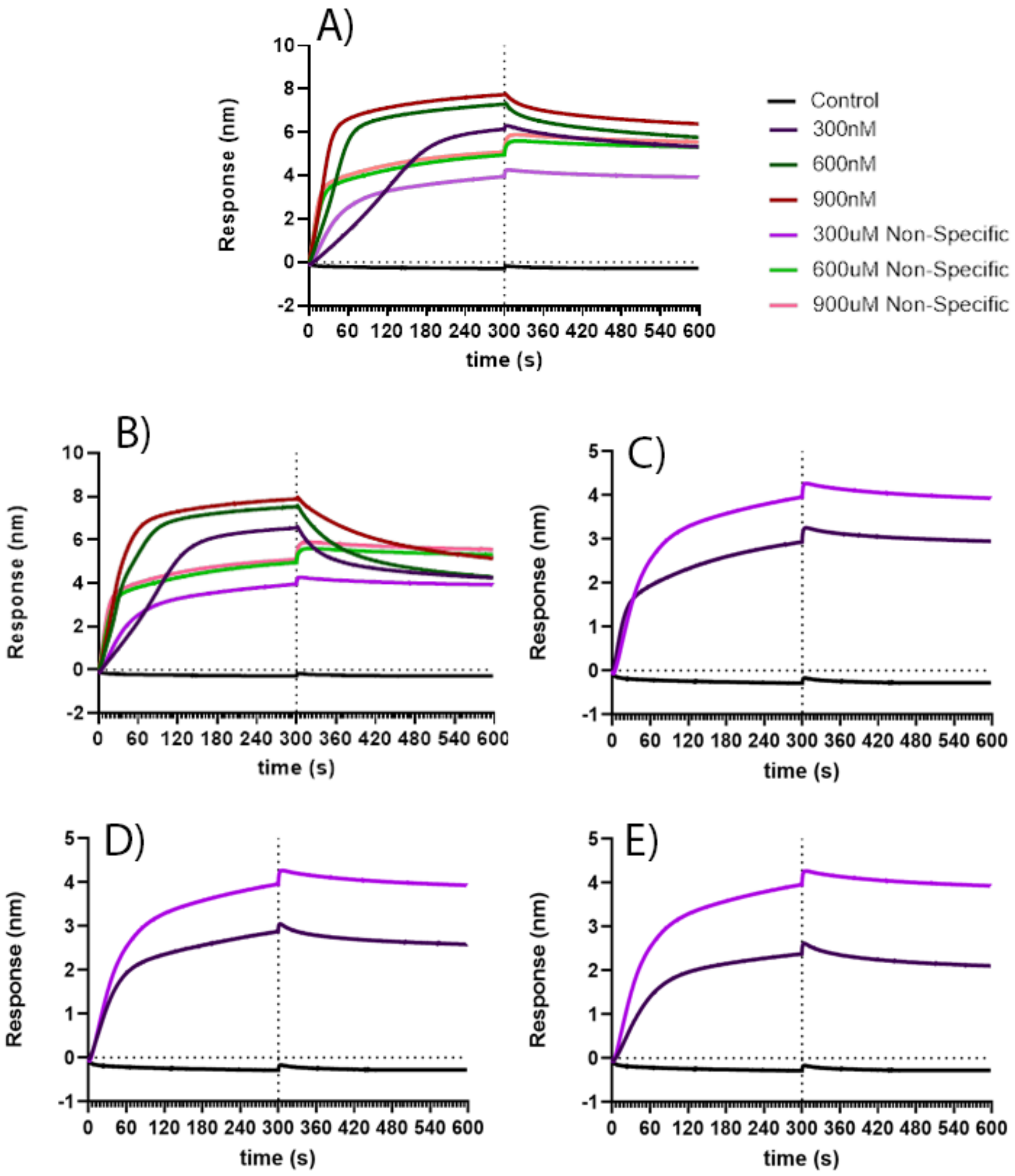


**Figure S2. TAZ2 binding to p53-TAD variants.** p53-TAD variants (A, WT; B, K24N; C, N29K/N30D; D, D49Y; E, W53G) were subjected to different concentrations of TAZ2. The traces for non-specific binding of TAZ2 to the Ni-NTA biosensor tip in the absence of p53-TAD are denoted as "Non-Specific." The control traces were obtained with p53-TAD on the tip without including TAZ2 and demonstrate that the p53-TAD variants were not depleted during BLI runs. Vertical line 300 s denotes transition from the association phase to the dissociation phase. The traces for TAZ2 with N29K/N30D, D49Y and W53G did not show specific protein-protein interactions and were recorded at the lowest TAZ2 concentration to minimize non-specific binding. The traces shown are averages from 2 independent experiments.

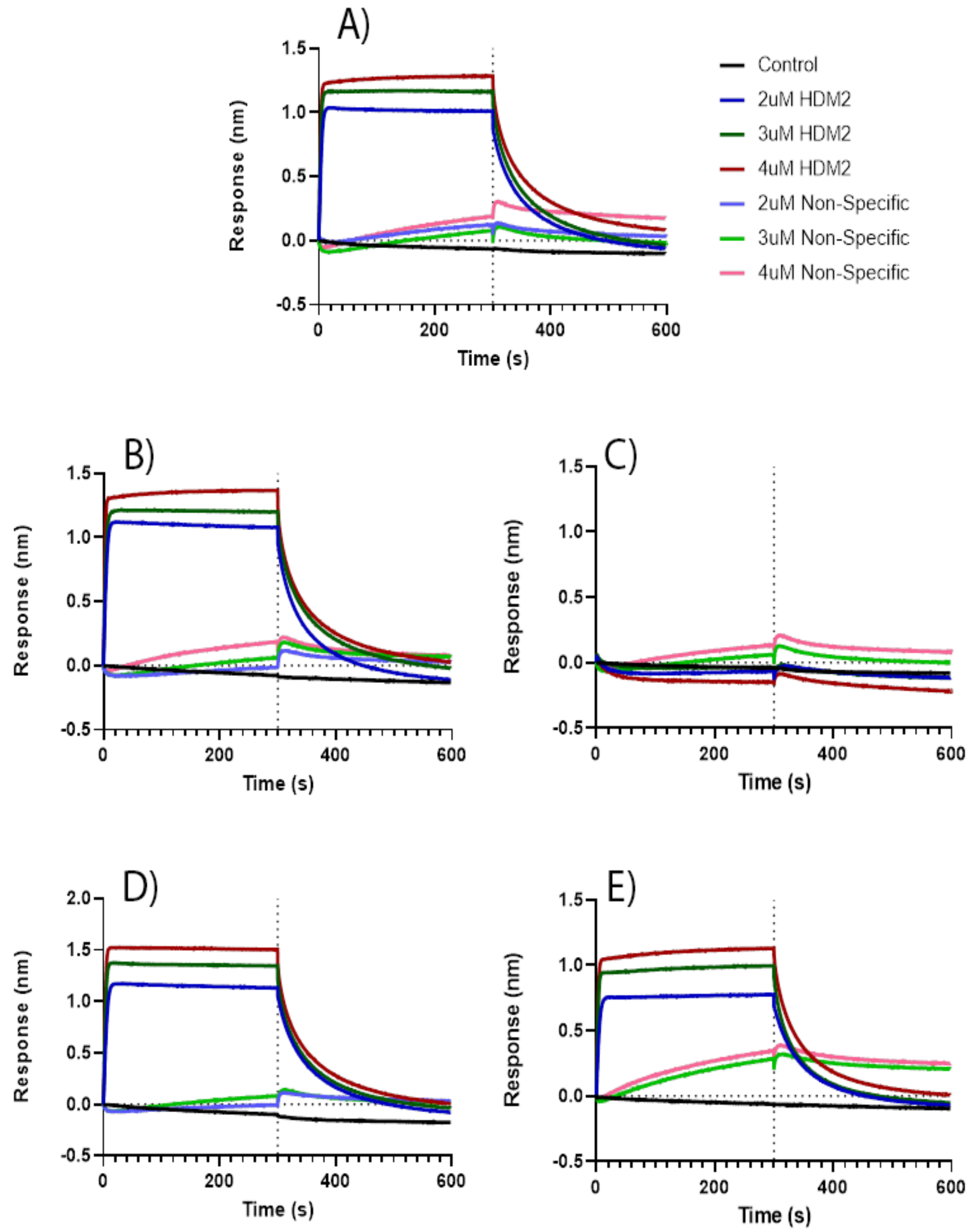


**Figure S3. HDM2 binding to p53-TAD variants.** p53-TAD variants (A, WT; B, K24N; C, N29K/N30D; D, D49Y; E, W53G) were subjected to 3 different concentrations of HDM2. The traces for non-specific binding of HDM2 to the Ni-NTA tip in the absence of p53-TAD are denoted as "Non-Specific." The control traces were obtained with each p53-TAD variant on the tip without including HDM2 and demonstrate that the p53-TAD variants were not depleted during BLI runs. Vertical line 300 s denotes transition from the association phase to the dissociation phase. The traces shown are averages from 2 independent experiments.

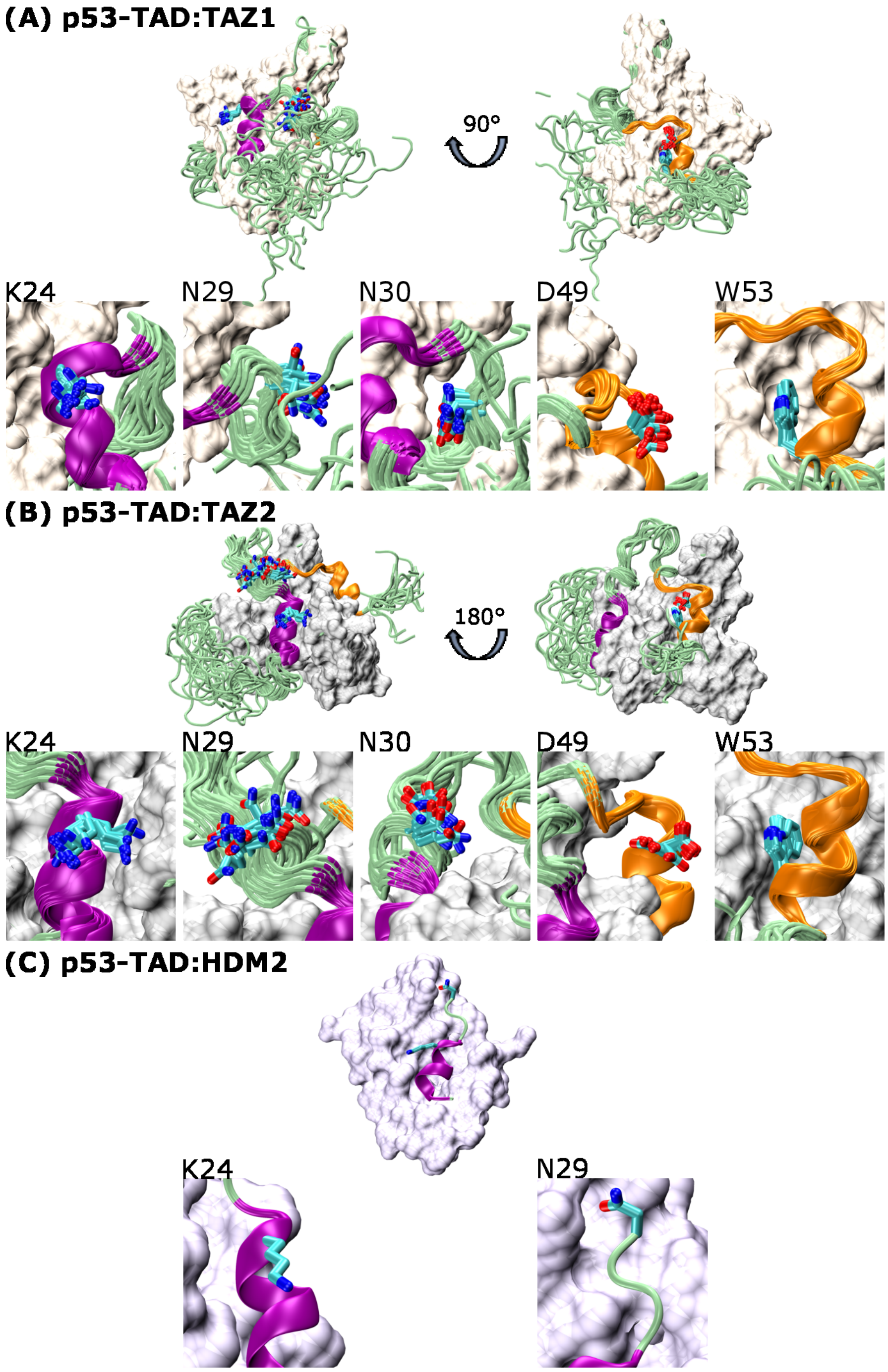


**Figure S4. Binding interface between p53-TAD and TAZ1 (A), TAZ2 (B), or HDM2 (C).** For each panel, the first row shows an overview of the protein complex, where p53-TAD is shown in a cartoon representation and its binding partner is shown as a molecular surface. The second row in each panel illustrates the orientation of p53-TAD residues K24, N29, N30, D49 and W53 at the binding interface. P53-TAD subdomain AD1 is in purple, AD2 is in orange, and the residues subjected to mutations in this study are shown in licorice. PDB IDs are 5HOU[49], 5HPD[49] and 1YCR[31] in panels A, B, and C, respectively. For panels A and B, all 20 models of p53-TAD were shown to illustrate a contrast between stably folded and disordered regions.

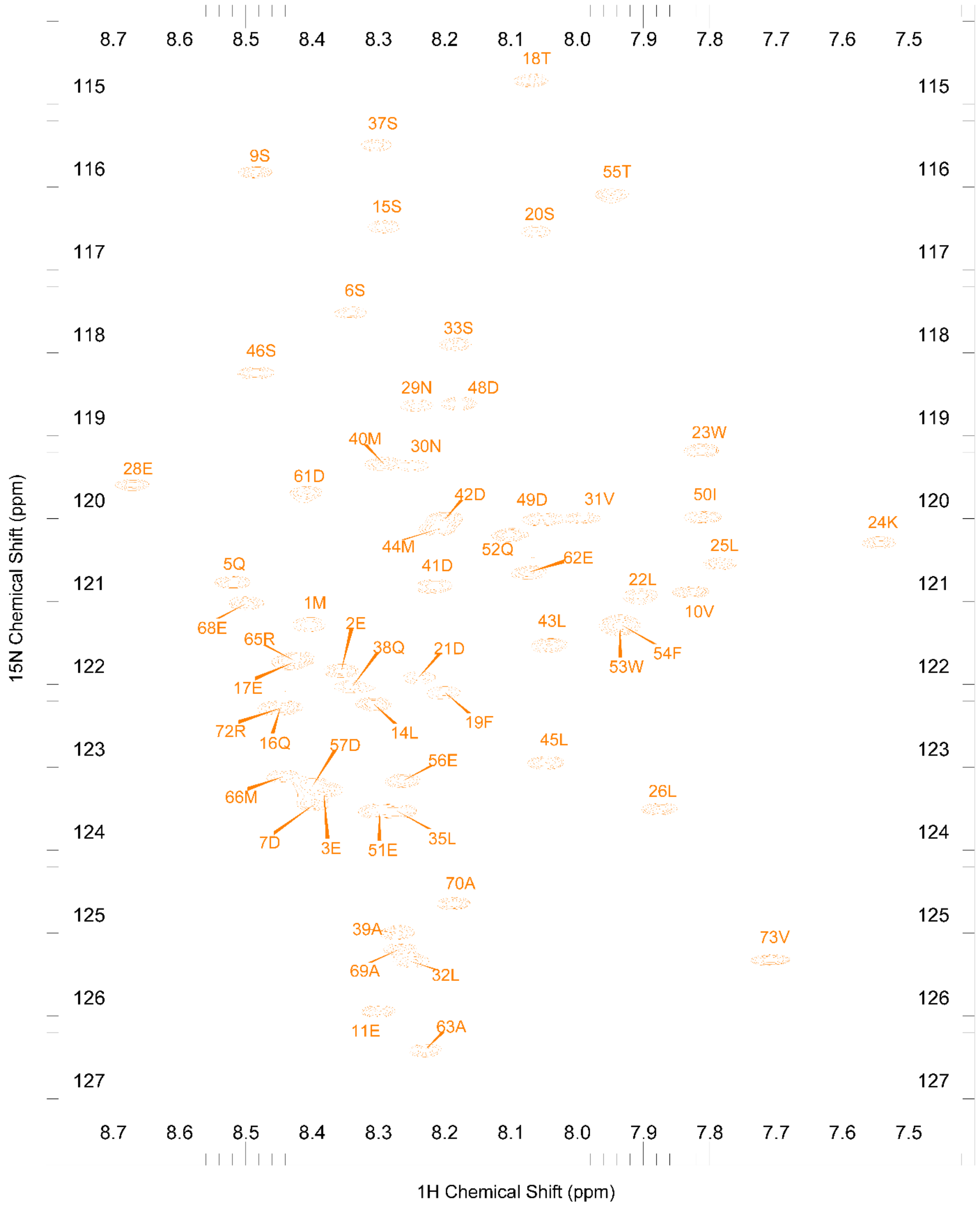


**Figure S5. $^{15}$N-HSQC of p53-TAD$^{WT}$.** The $^{15}$N-HSQC spectrum of p53-TAD$^{WT}$ with resonance assignments derived from Vise et al, 2005.

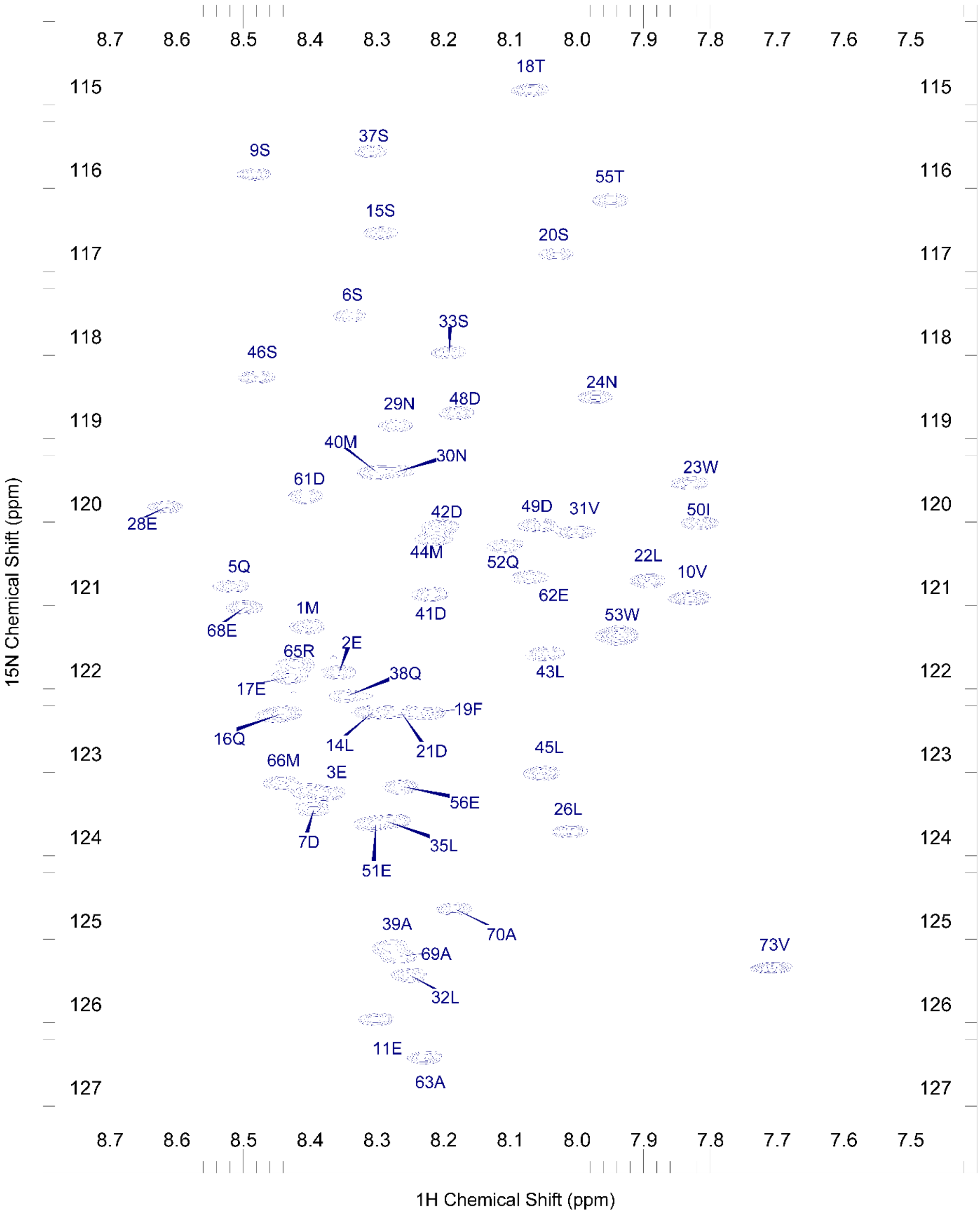


**Figure S6. $^{15}$N-HSQC of p53-TAD$^{K24N}$.** The $^{15}$N-HSQC spectrum of p53-TAD$^{K24N}$ with assignments derived from Vise et al, 2005. Orphaned peaks within the $^{15}$N-HSQC spectrum of p53-TAD$^{K24N}$ were assigned based upon proximity to unassigned positions derived from $^{15}$N-HSQC spectrum of p53-TAD$^{WT}$.

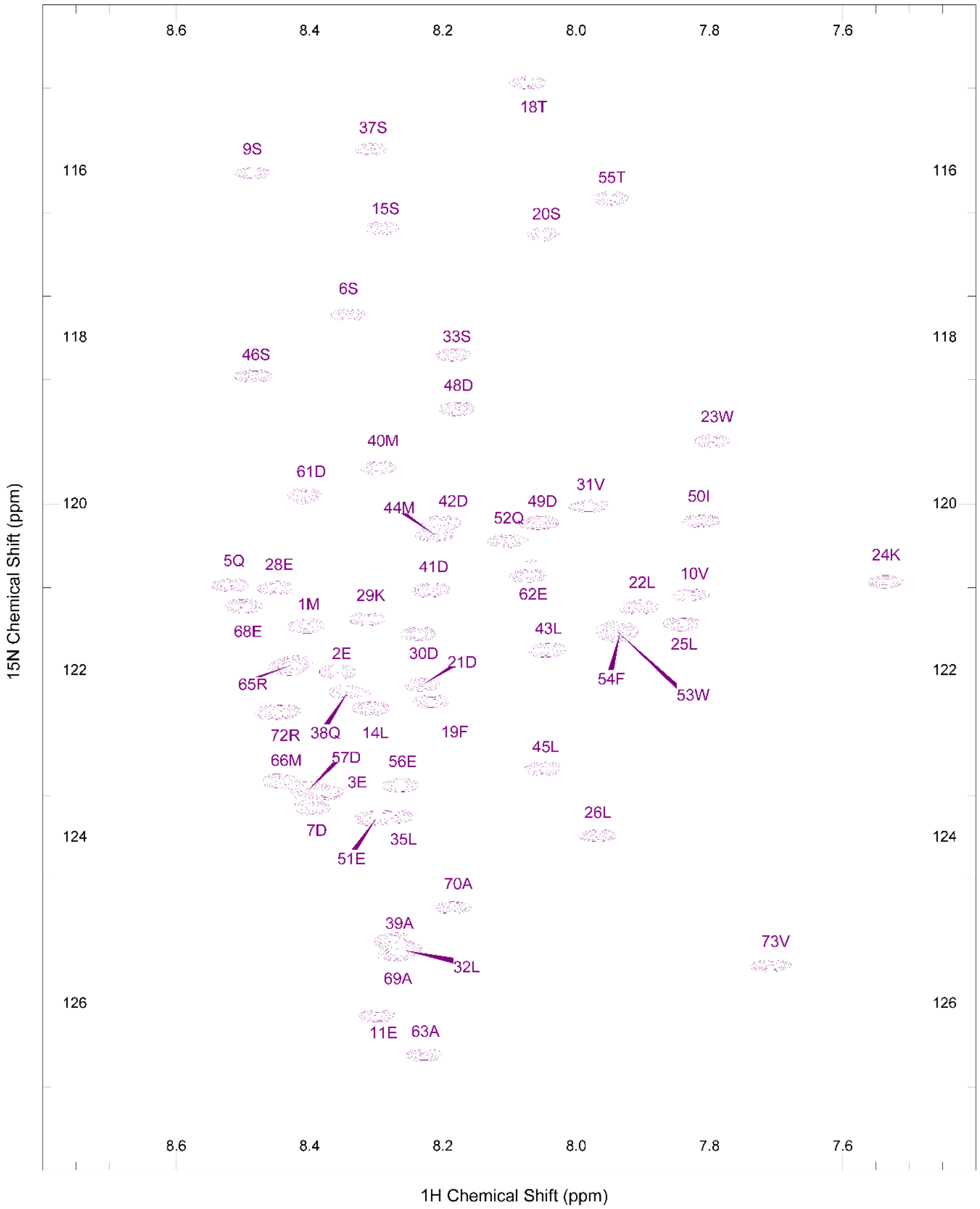


**Figure S7. $^{15}$N-HSQC of p53-TAD$^{N29K/N30D}$.** The $^{15}$N-HSQC spectrum of p53-TAD$^{N29K/N30D}$ with assignments derived from Vise et al, 2005. Orphaned peaks within the $^{15}$N-HSQC spectrum of p53-TAD$^{N29K/N30D}$ were assigned based upon proximity to unassigned positions derived from $^{15}$N-HSQC spectrum of p53-TAD$^{WT}$.

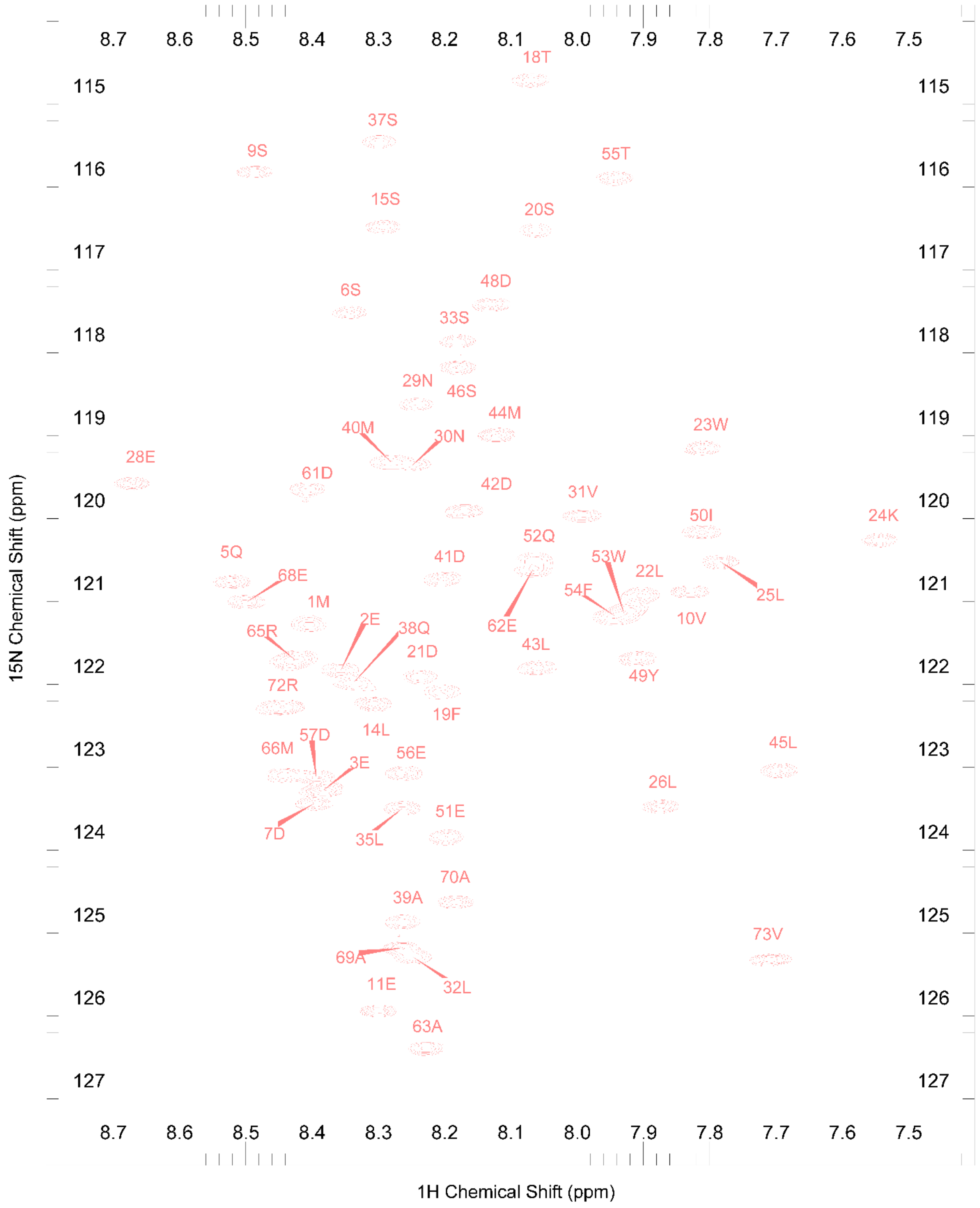


**Figure S8. $^{15}$N-HSQC of p53-TAD$^{D49Y}$.** The $^{15}$N-HSQC spectrum of p53-TAD$^{D49Y}$ with assignments derived from Vise et al, 2005. Orphaned peaks within the $^{15}$N-HSQC spectrum of p53-TAD$^{D49Y}$ were assigned based upon proximity to unassigned positions derived from $^{15}$N-HSQC spectrum of p53-TAD$^{WT}$.

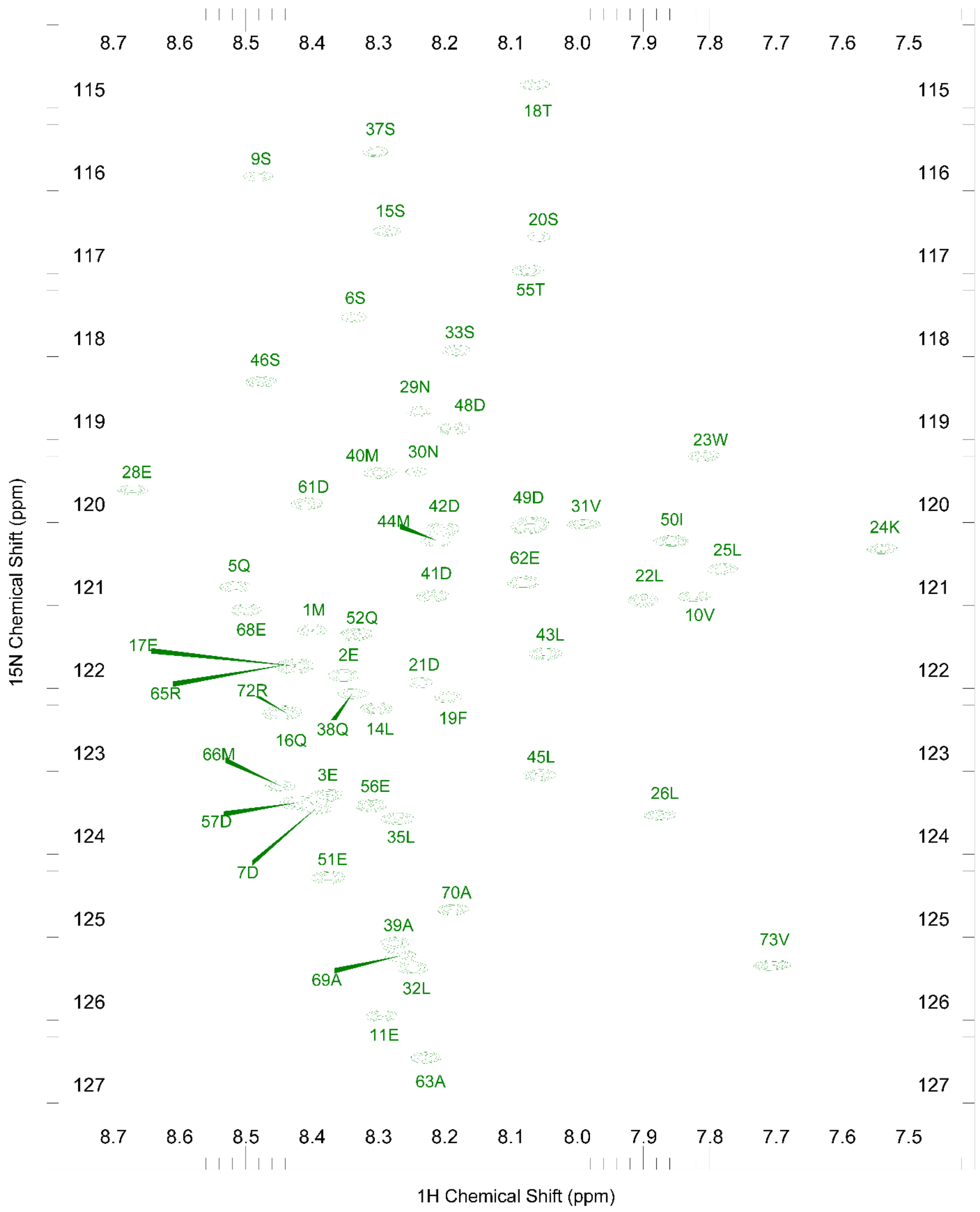


**Figure S9. $^{15}$N-HSQC of p53-TAD$^{W53G}$.** The $^{15}$N-HSQC spectrum of p53-TAD$^{W53G}$ with assignments derived from Vise et al, 2005. Orphaned peaks within the $^{15}$N-HSQC spectrum of p53-TAD$^{D49Y}$ were assigned based upon proximity to unassigned positions derived from $^{15}$N-HSQC spectrum of p53-TAD$^{WT}$.

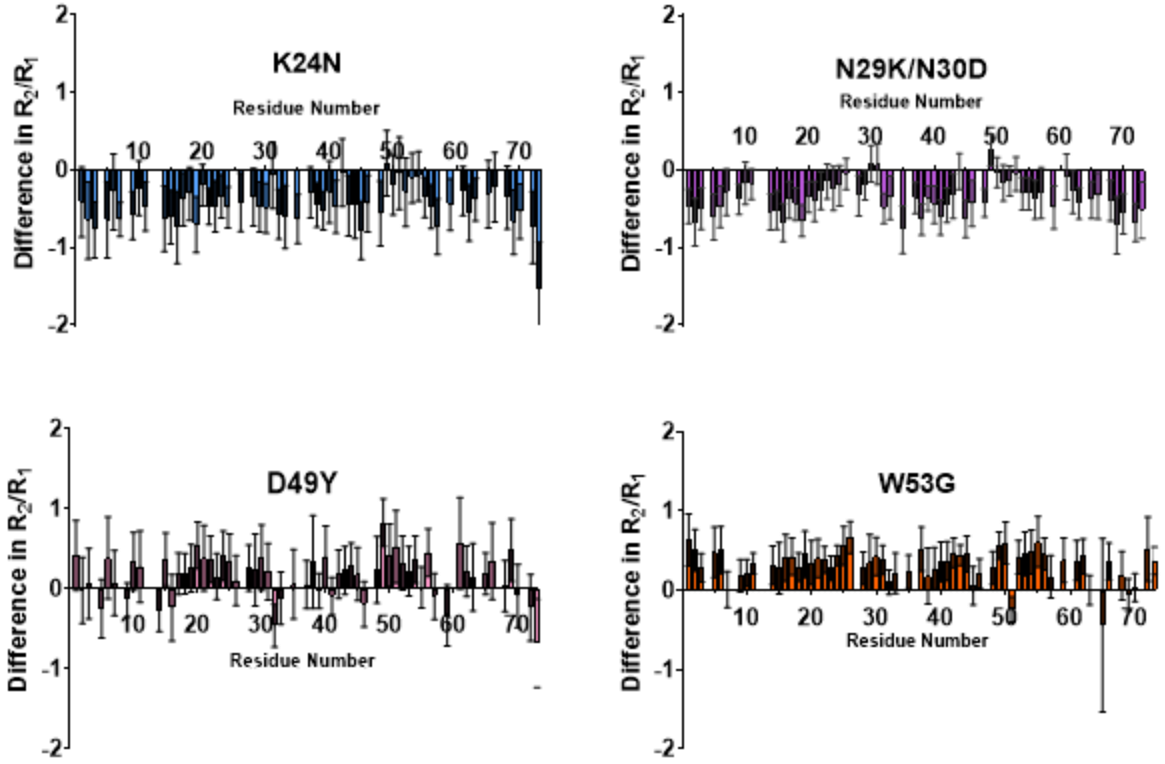


**Figure S10. Difference in R2/R1 between p53-TAD variants and WT.** Error bars reflect 95% confidence interval derived from R1 and R2 values.

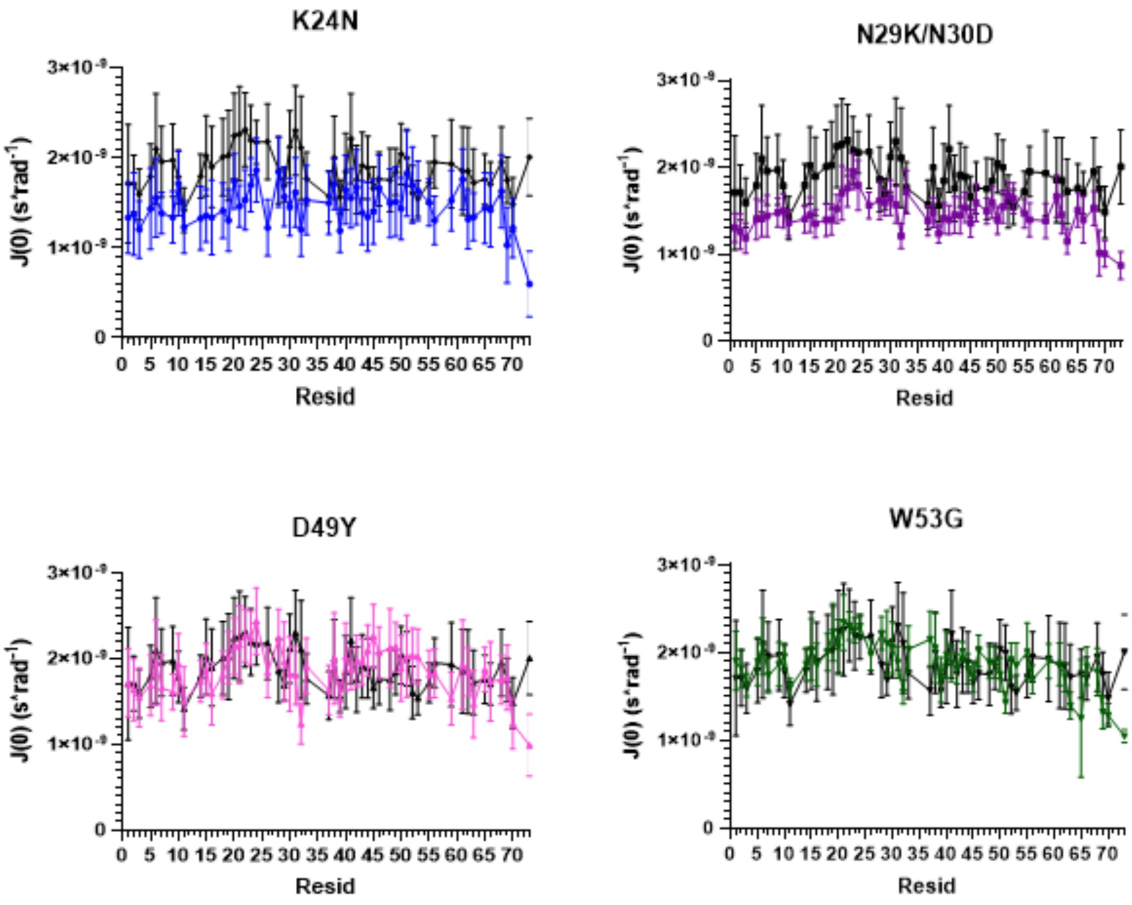


**Figure S11. Reduced spectral density approximation of *J(0)* for p53-TAD variants.** An approximation of relaxation power at a frequency of 0Hz. Color represents the corresponding variant while black represents WT values. Error bars represent the 95% confidence interval derived from $R_1$ and $R_2$ exponential decay fitting.

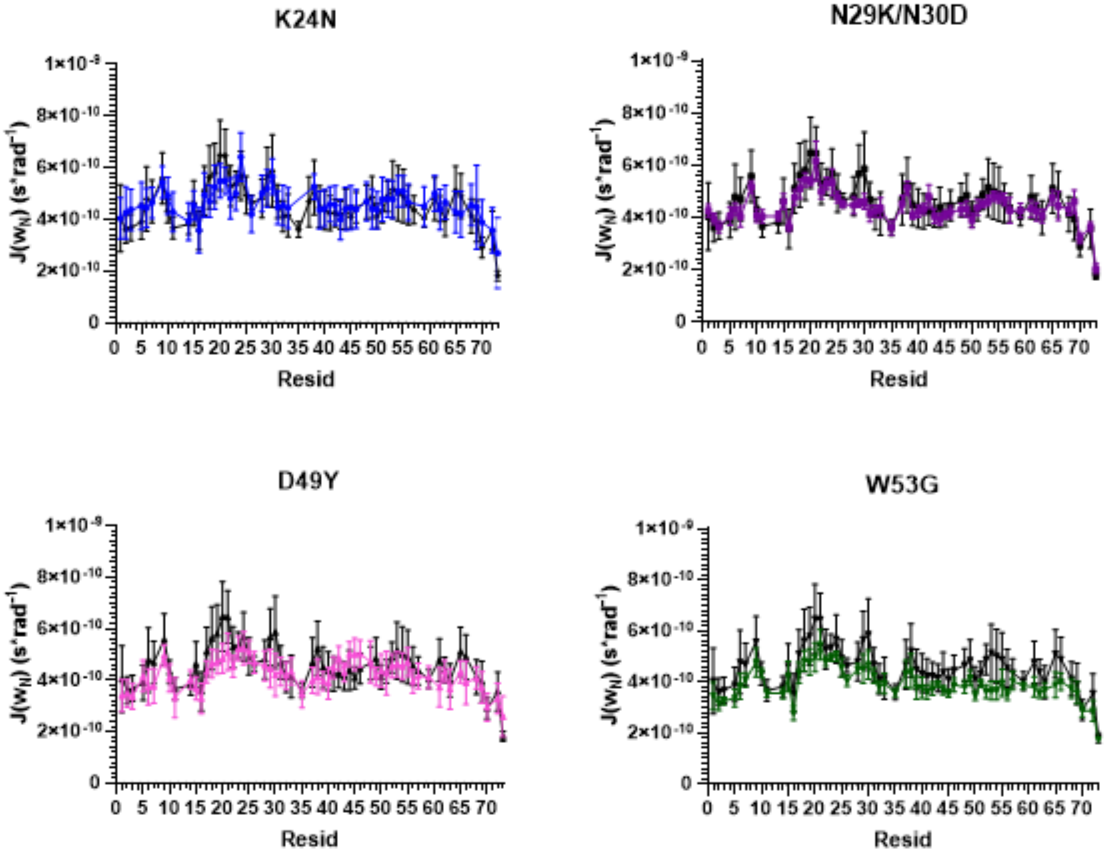


**Figure S12. Reduced spectral density approximation of *J(ω_N)* for p53-TAD variants.** An approximation of relaxation power at a frequency of 50.7MHz, Larmor frequency of $^{15}N$ in 11.74T field. Color represents the corresponding mutant variant while black represents WT values. Error bars represent the 95% confidence interval derived from $R_1$ and $R_2$ exponential decay fitting.

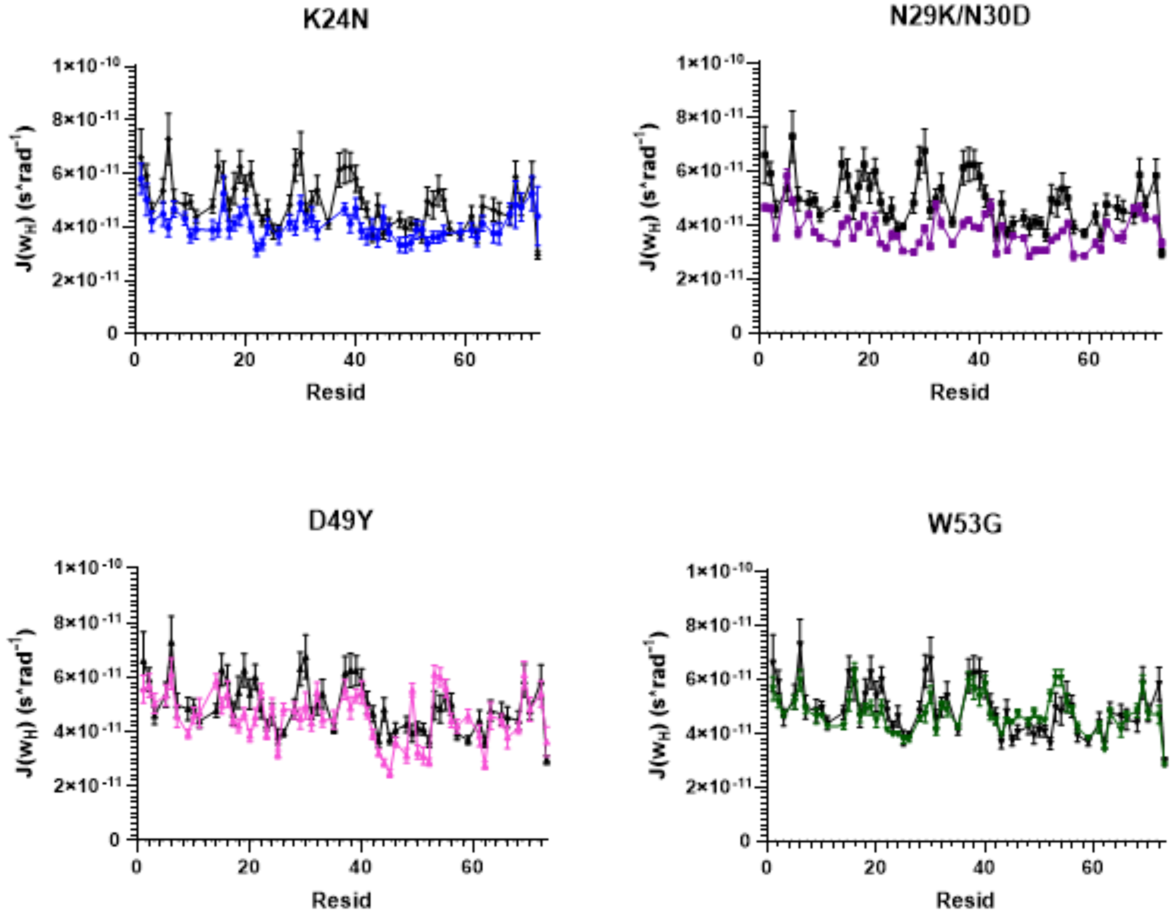


**Figure S13. Reduced spectral density approximation of $J(\omega_H)$ for p53-TAD variants.** An approximation of relaxation power at a frequency of 435MHz, Larmor frequency of $^1$H in 11.74T field. Color represents the corresponding mutant variant while black represents WT values. Error bars represent the 95% confidence interval derived from $R_1$ and $R_2$ exponential decay fitting.

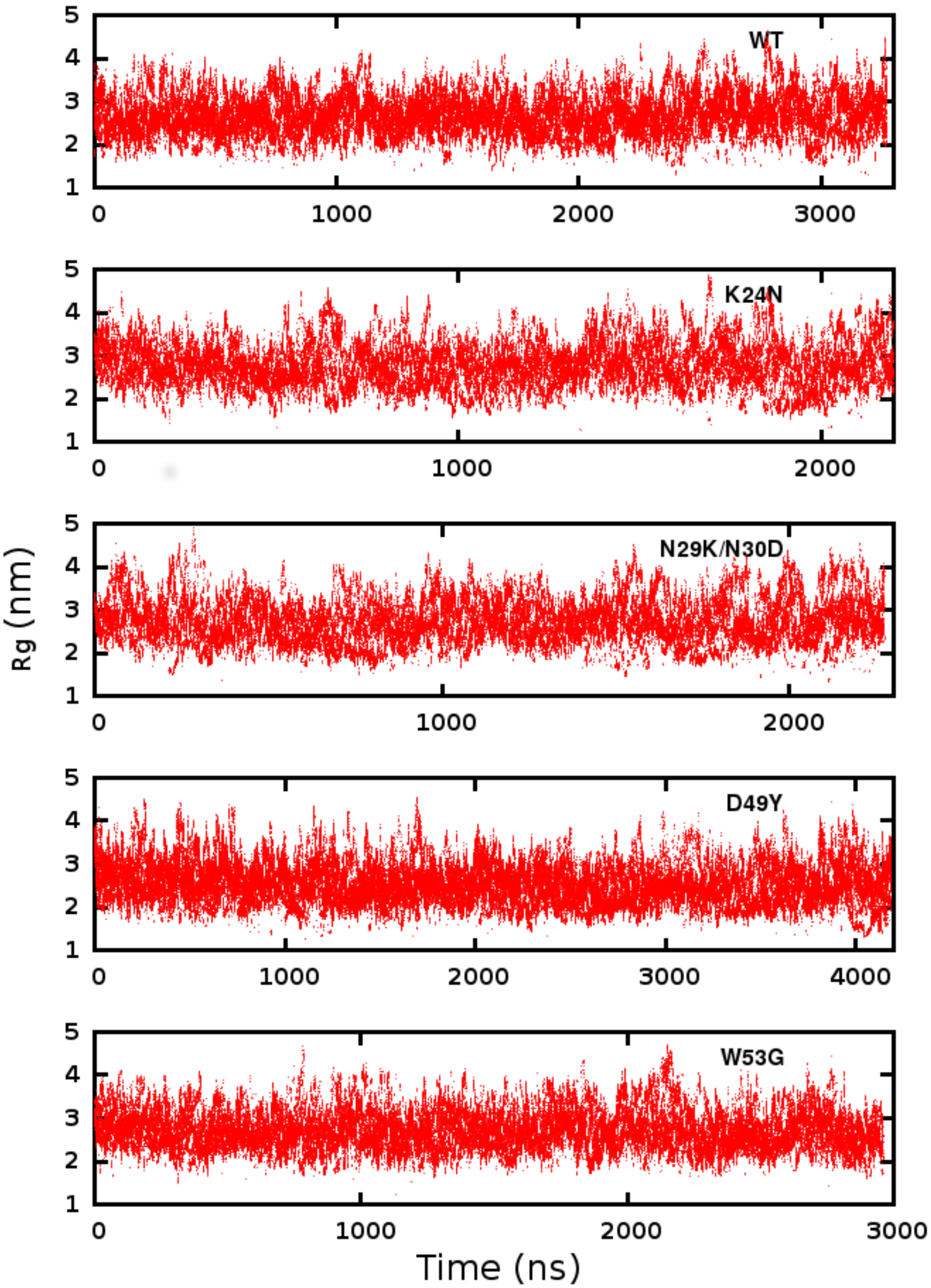


**Figure S14.** ***$R_g$*** **of p53-TAD variants as a function of simulation time at 298 K.**

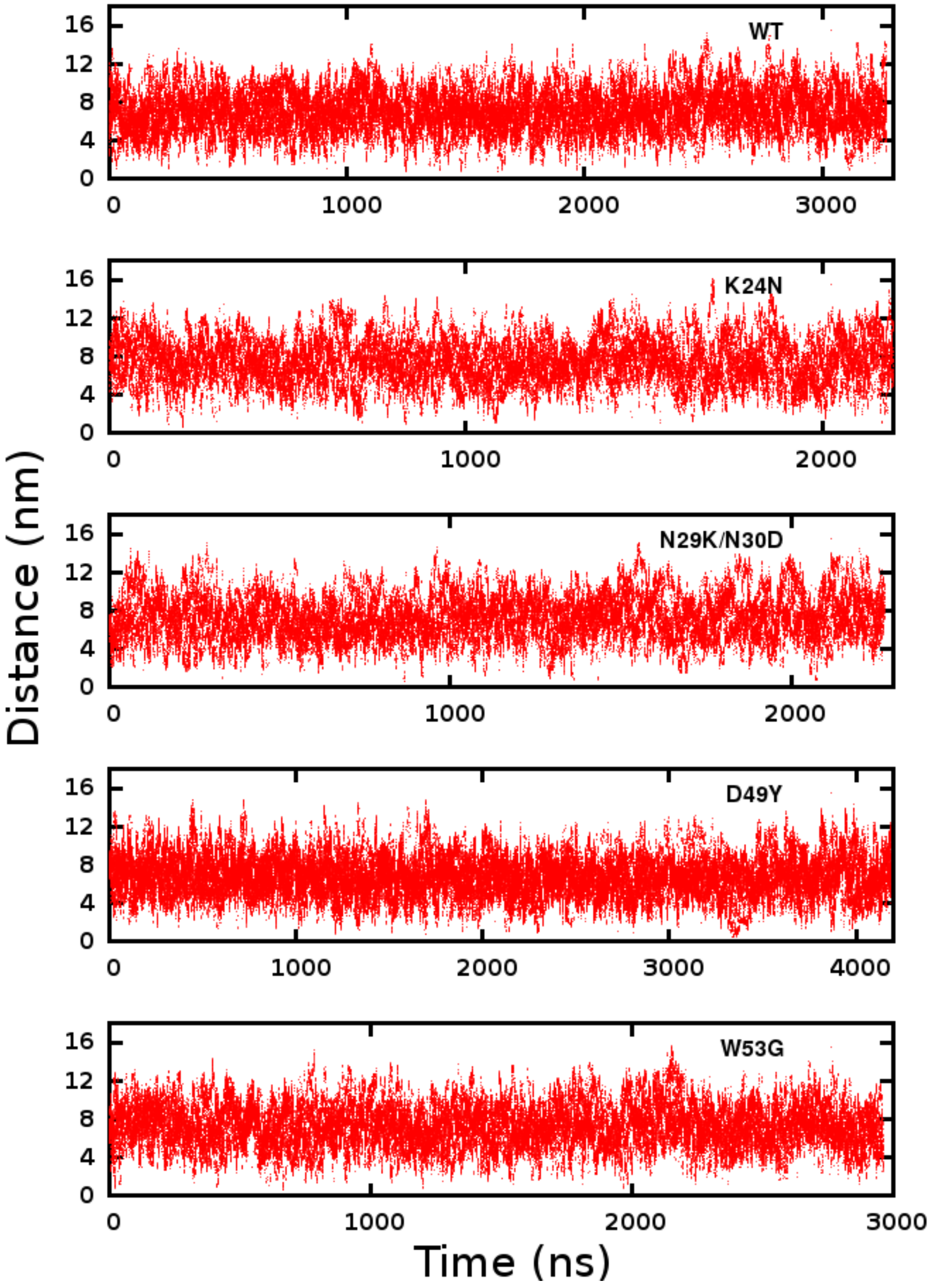


**Figure S15. End-to-end distance of p53-TAD variants as a function of simulation time at 298 K.**

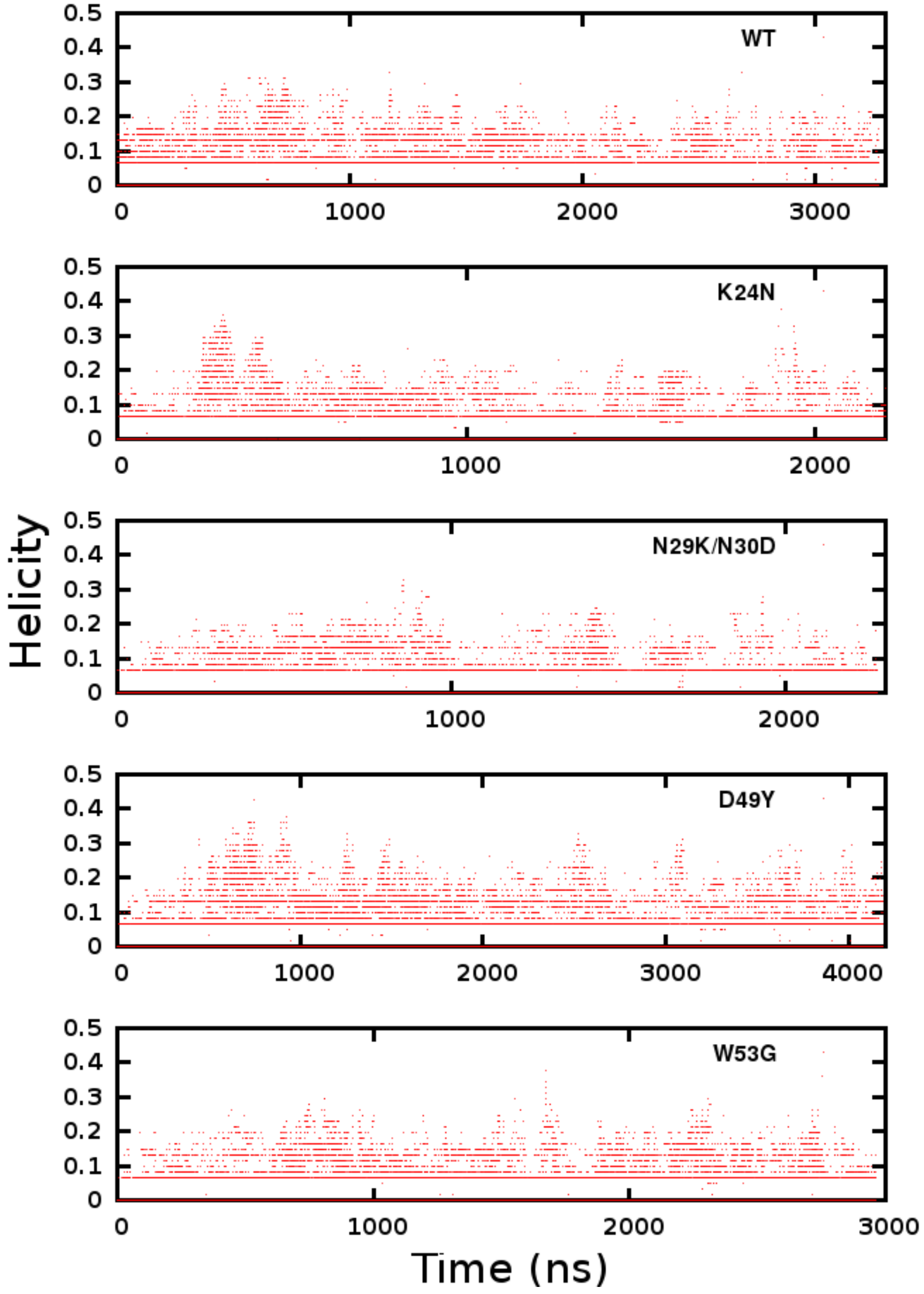


**Figure S16. Overall helicity of p53-TAD variants as a function of simulation time at 298 K.**

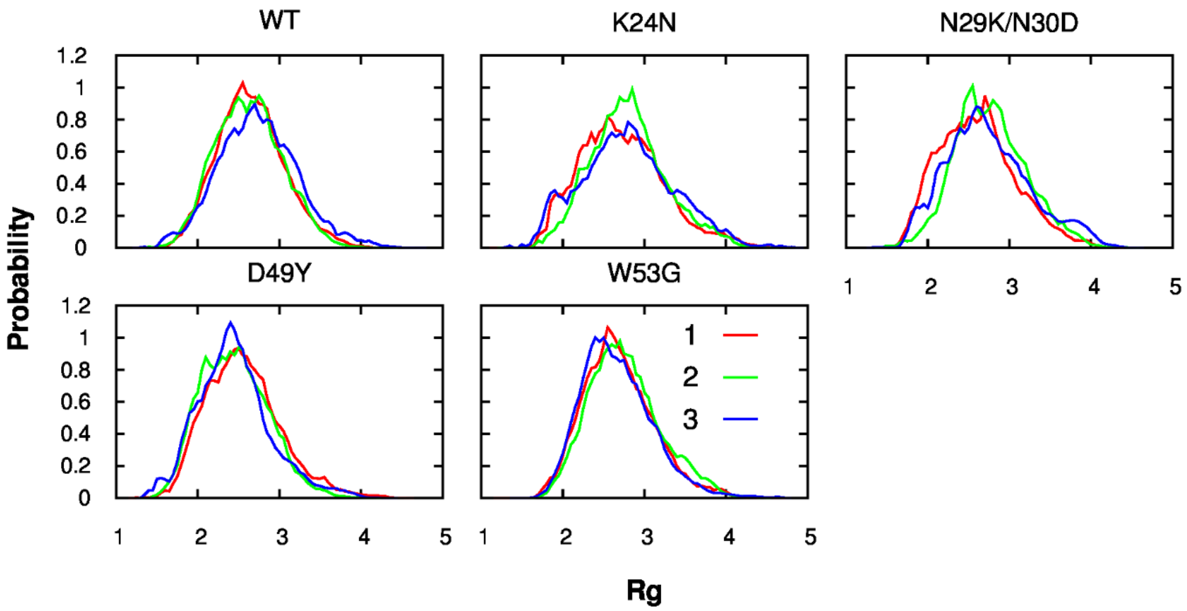


**Figure S17. Probability distribution of peptide $R_g$ for three equally divided trajectory segments.**

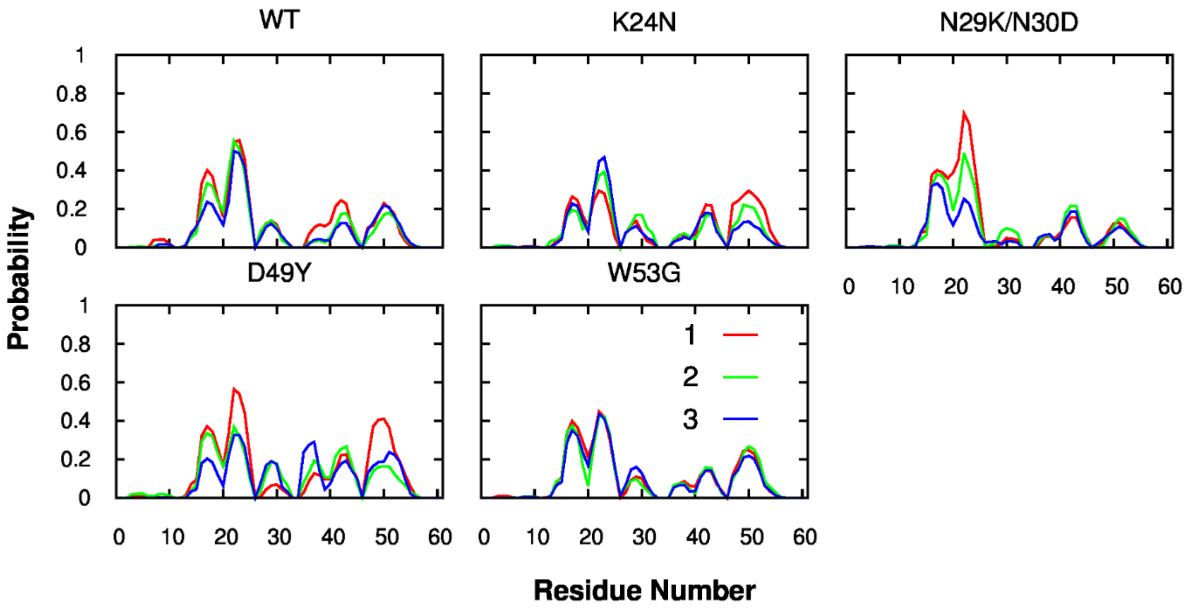


**Figure S18. Residue helical propensity (including both α helices and $3_{10}$ helices) for three equally divided trajectory segments.**

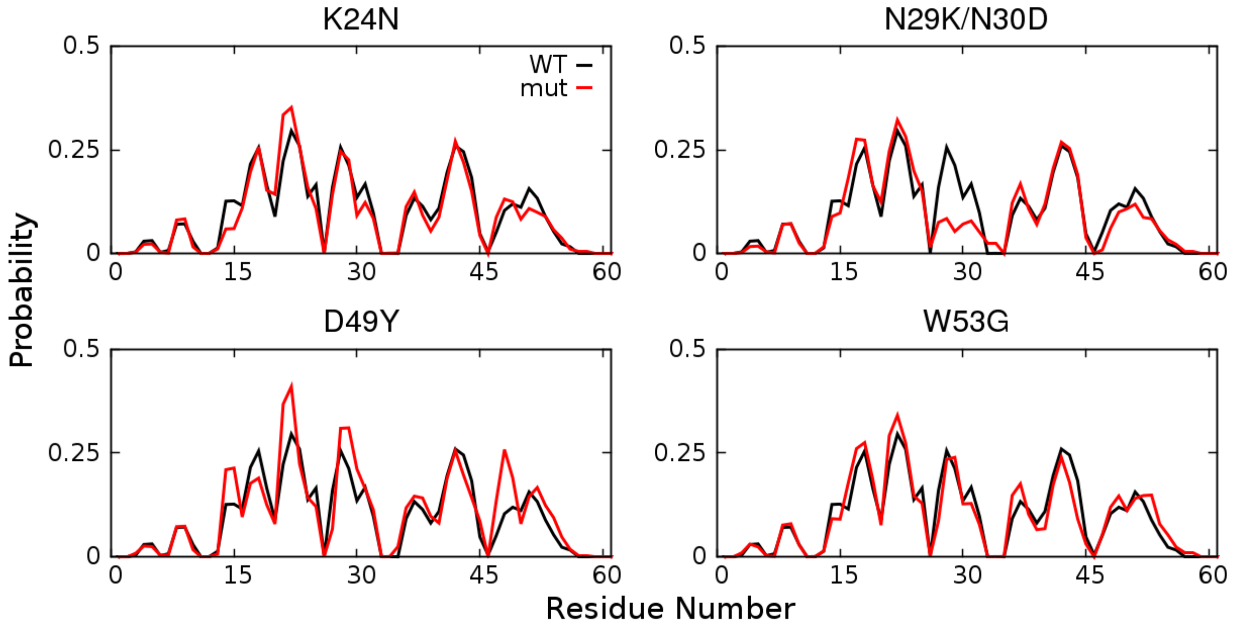


**Figure S19. Probability of forming helical turns in p53-TAD mutants in comparison with WT.**

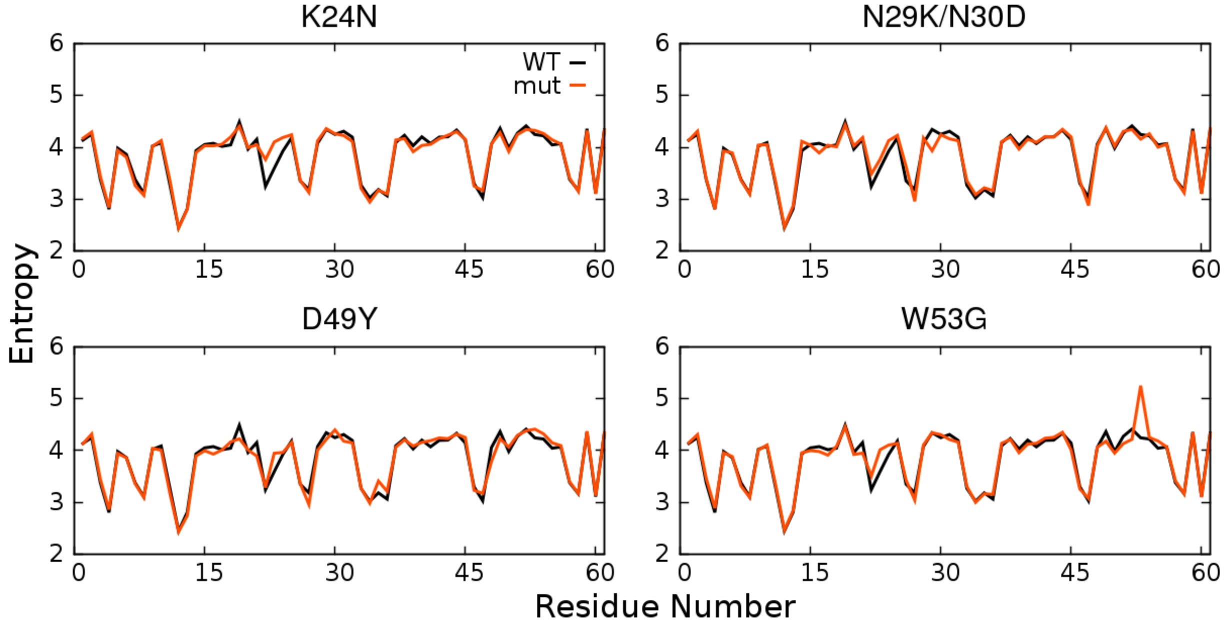


**Figure S20. Residue backbone entropy of p53-TAD mutants in comparison with WT.**

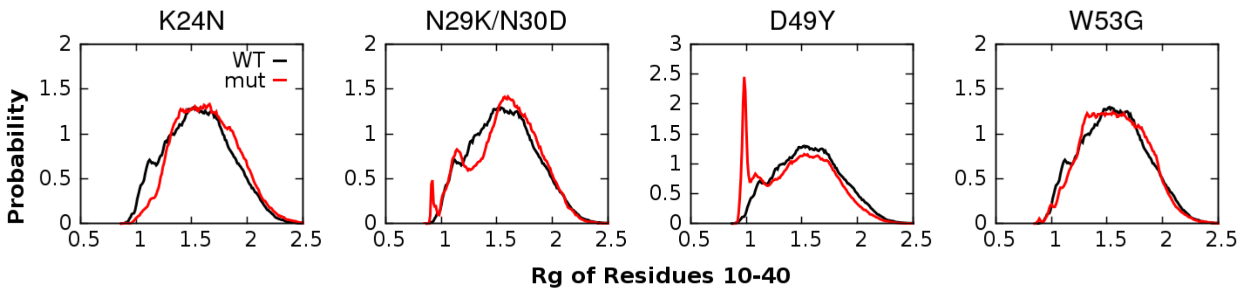


**Figure S21. Probability distributions of $R_g$ of p53-TAD residues 10-40 for four mutants in comparison with the WT results.**

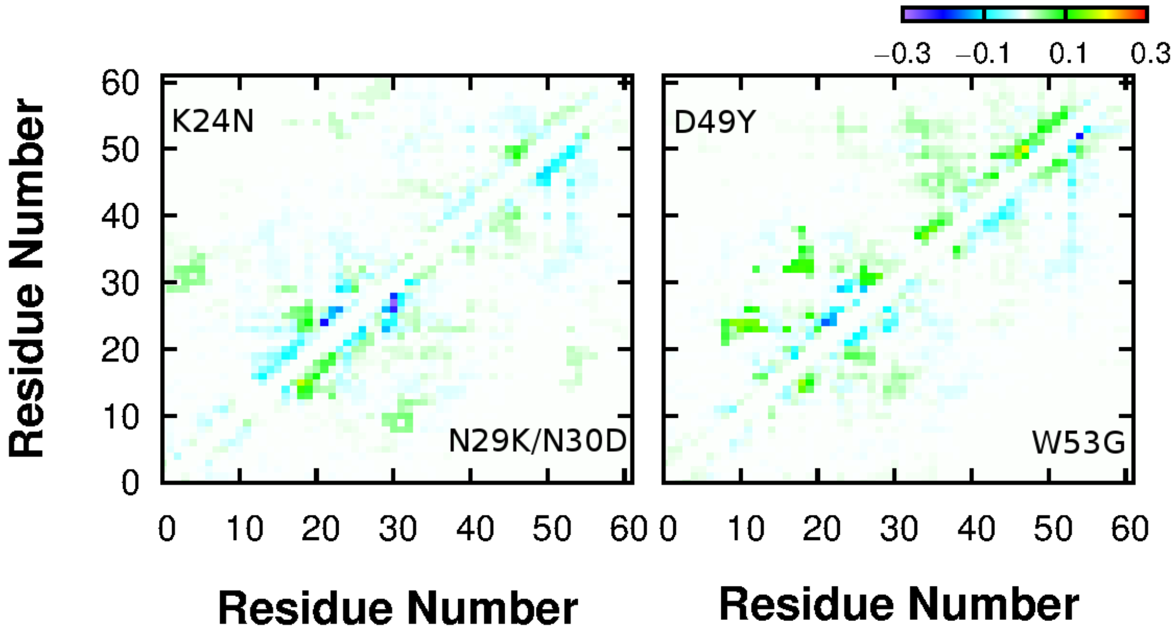


**Figure S22. Difference of contact probabilities between mutants and WT.**

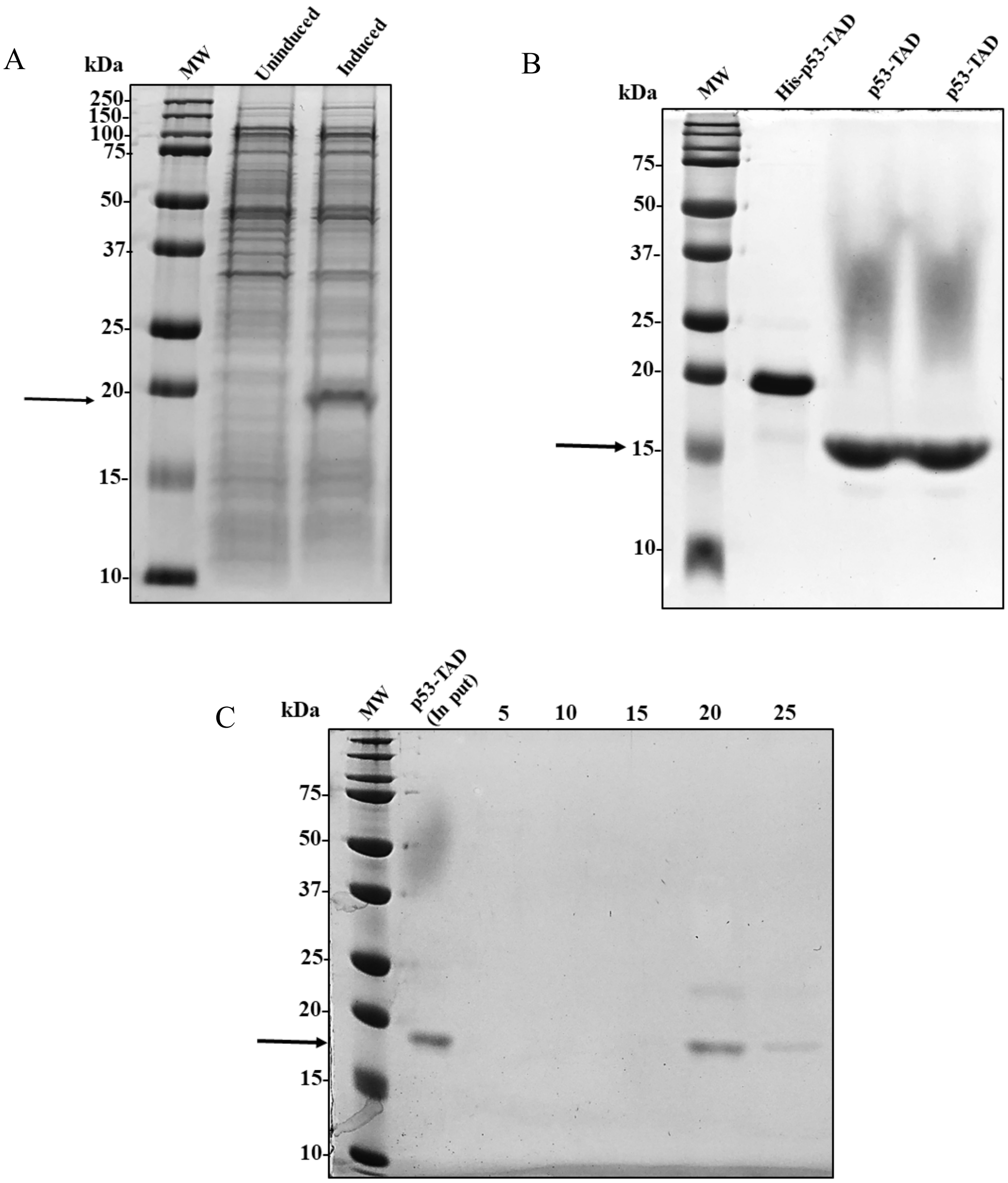


**Figure S23: His-p53-TAD (1-73) wild type expression and purification.** A) The p53-TAD wild type LOBSTR-BL21(DE3) *E. coli* cells were grown in LB media with 50 µg/ml kanamycin and 1% glucose at 30°C until $OD_{600}$ = 0.5-0.7. The culture was induced with 1 mM IPTG and grown at 37°C for an additional 6 hrs. B) The N-terminal His-tag was cleaved using Thrombin CleanCleave kit for 6 hrs at room temperature. (The His-tag was removed for NMR experiments only). The cleaved p53-TAD protein was eluted using column recovery method, according to manufacturer's protocol. C) The final purification was done using gel-filtration chromatography and every 5th fraction was analyzed as shown in the figure.

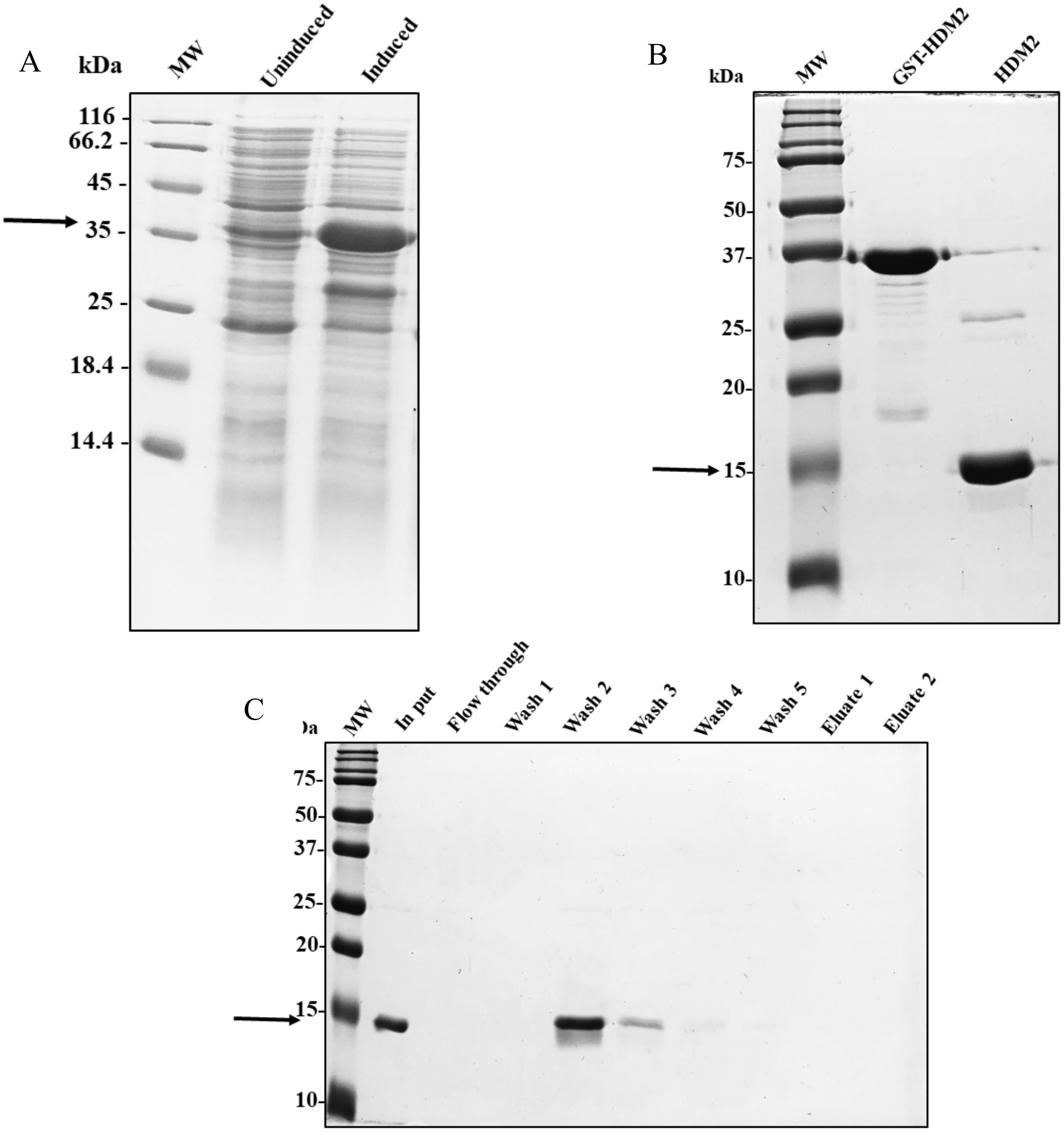


**Figure S24: GST-HDM2 (17-125) expression and purification .** A) Rosetta BL21 (DE3) *E. coli* cells containing GST-HDM2 plasmid were grown in LB media with 25 µg/ml Chloramphenicol and 100 µg/ml Ampicillin at 37°C until $OD_{600}$ = 0.8. The culture was induced with 1mM IPTG and grown at 25°C for an additional 5 hrs. B) The GST tag was removed using HRV 3C protease with an enzyme to substrate ratio of 1:100 at 4°C for 16 hrs. C) The cleaved HDM2 protein was eluted in the wash fraction using a Glutathione Sepharose 4B resin column as a final step of purification to remove the trace amount of GST tagged HRV 3C protease.

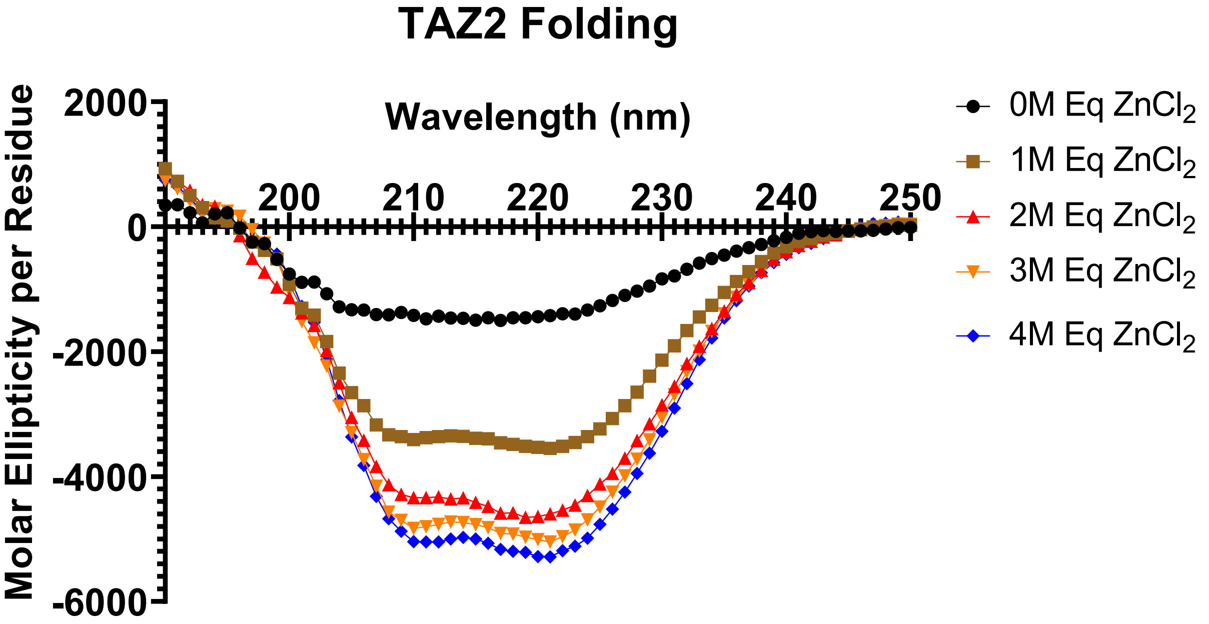


**Figure S25. Folding of TAZ2 observed by Circular Dichropolarimetry (CD).** 160uM of TAZ2 suspended in 20mMTris, 50mM NaCl, 10mM DTT at pH=7.5. TAZ2 folding was monitored using CD operating with a range of 190-250nm, scan speed 1nm/s, and 10 cumulant averages represented after buffer subtraction. Molar equivalent additions of $ZnCl_2$ were then applied to TAZ2 solution in a stepwise manner and allowed 10 minutes for folding at room temperature.

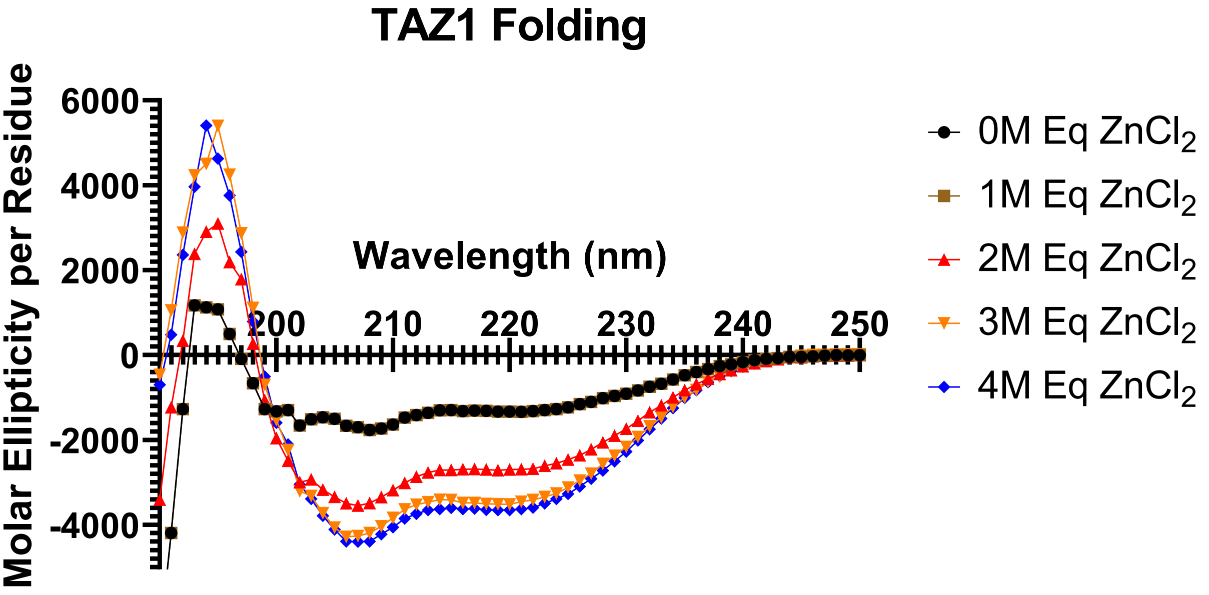


**Figure S26. Folding of TAZ1 observed by Circular Dichropolarimetry (CD).** 200uM of TAZ1 suspended in 20mMTris, 50mM NaCl, 10mM DTT at pH=6.9. TAZ1 folding was monitored using CD operating with a range of 190-250nm, scan speed 1nm/s, and 10 cumulant averages represented after buffer subtraction. Molar equivalent additions of $ZnCl_2$ were then applied to TAZ1 solution in a stepwise manner and allowed 10 minutes for folding at room temperature.

**Table S1. $^{15}$N-HSQC assignments for p53-TAD$^{WT}$.**

| p53-TAD$^{WT}$ | | | | | |
|---|---|---|---|---|---|
| Residue | N (ppm) | HN (ppm) | Residue | N (ppm) | HN (ppm) |
| 1Met | 121.47 | 8.41 | 38Gln | 122.22 | 8.34 |
| 2Glu | 122.03 | 8.36 | 39Ala | 125.2 | 8.27 |
| 3Glu | 123.47 | 8.38 | 40Met | 119.54 | 8.29 |
| 5Gln | 120.96 | 8.52 | 41Asp | 121.01 | 8.22 |
| 6Ser | 117.71 | 8.34 | 42Asp | 120.2 | 8.2 |
| 7Asp | 123.65 | 8.39 | 43Leu | 121.73 | 8.04 |
| 9Ser | 116.02 | 8.49 | 44Met | 120.34 | 8.21 |
| 10Val | 121.08 | 7.83 | 45Leu | 123.15 | 8.05 |
| 11Glu | 126.13 | 8.3 | 46Ser | 118.44 | 8.48 |
| 14Leu | 122.44 | 8.3 | 48Asp | 118.81 | 8.18 |
| 15Ser | 116.67 | 8.29 | 49Asp | 120.21 | 8.06 |
| 16Gln | 122.49 | 8.46 | 50Ile | 120.17 | 7.81 |
| 17Glu | 121.92 | 8.44 | 51Glu | 123.72 | 8.3 |
| 18Thr | 114.9 | 8.07 | 52Gln | 120.39 | 8.1 |
| 19Phe | 122.3 | 8.2 | 53Trp | 121.49 | 7.94 |
| 20Ser | 116.74 | 8.06 | 54Phe | 121.49 | 7.94 |
| 21Asp | 122.12 | 8.24 | 55Thr | 116.28 | 7.95 |
| 22Leu | 121.13 | 7.91 | 56Glu | 123.36 | 8.26 |
| 23Trp | 119.37 | 7.82 | 57Asp | 123.41 | 8.4 |
| 24Lys | 120.49 | 7.55 | 59Gly | 109.33 | 8.36 |
| 25Leu | 120.76 | 7.79 | 61Asp | 119.88 | 8.41 |
| 26Leu | 123.7 | 7.88 | 62Glu | 120.84 | 8.07 |
| 28Glu | 119.8 | 8.67 | 63Ala | 126.61 | 8.23 |
| 29Asn | 118.84 | 8.25 | 65Arg | 121.9 | 8.42 |
| 30Asn | 119.56 | 8.25 | 66Met | 123.3 | 8.44 |
| 31Val | 120.18 | 8 | 68Glu | 121.22 | 8.5 |
| 32Leu | 125.53 | 8.25 | 69Ala | 125.39 | 8.26 |
| 33Ser | 118.09 | 8.18 | 70Ala | 124.84 | 8.18 |
| 35Leu | 123.72 | 8.27 | 72Arg | 122.47 | 8.44 |
| 37Ser | 115.69 | 8.3 | 73Val | 125.53 | 7.71 |

**Table S2.** $^{15}$N-HSQC assignments for p53-TAD$^{K24N}$.

| p53-TAD$^{K24N}$ | | | | | |
|---|---|---|---|---|---|
| Residue | N (ppm) | HN (ppm) | Residue | N (ppm) | HN (ppm) |
| 1Met | 121.47 | 8.41 | 38Gln | 122.24 | 8.34 |
| 2Glu | 122.02 | 8.36 | 39Ala | 125.23 | 8.27 |
| 3Glu | 123.47 | 8.38 | 40Met | 119.56 | 8.29 |
| 5Gln | 120.96 | 8.52 | 41Asp | 121.01 | 8.22 |
| 6Ser | 117.72 | 8.34 | 42Asp | 120.22 | 8.2 |
| 7Asp | 123.65 | 8.39 | 43Leu | 121.75 | 8.04 |
| 9Ser | 116.02 | 8.49 | 44Met | 120.36 | 8.21 |
| 10Val | 121.08 | 7.83 | 45Leu | 123.17 | 8.05 |
| 11Glu | 126.14 | 8.3 | 46Ser | 118.45 | 8.48 |
| 14Leu | 122.45 | 8.3 | 48Asp | 118.84 | 8.18 |
| 15Ser | 116.69 | 8.29 | 49Asp | 120.22 | 8.06 |
| 16Gln | 122.5 | 8.45 | 50Ile | 120.18 | 7.81 |
| 17Glu | 121.99 | 8.44 | 51Glu | 123.75 | 8.3 |
| 18Thr | 114.94 | 8.07 | 52Gln | 120.42 | 8.1 |
| 19Phe | 122.36 | 8.21 | 53Trp | 121.52 | 7.94 |
| 20Ser | 116.82 | 8.05 | 54Phe | 121.49 | 7.94 |
| 21Asp | 122.24 | 8.25 | 55Thr | 116.3 | 7.95 |
| 22Leu | 121.06 | 7.9 | 56Glu | 123.36 | 8.26 |
| 23Trp | 119.49 | 7.82 | 57Asp | 123.41 | 8.4 |
| 24Asn | 118.71 | 7.97 | 59Gly | 109.33 | 8.36 |
| 25Leu | - | - | 61Asp | 119.88 | 8.41 |
| 26Leu | 123.77 | 8.01 | 62Glu | 120.85 | 8.07 |
| 28Glu | 119.88 | 8.62 | 63Ala | 126.61 | 8.23 |
| 29Asn | 118.91 | 8.25 | 65Arg | 121.9 | 8.42 |
| 30Asn | 119.58 | 8.26 | 66Met | 123.31 | 8.44 |
| 31Val | 120.23 | 8 | 68Glu | 121.22 | 8.5 |
| 32Leu | 125.56 | 8.25 | 69Ala | 125.39 | 8.26 |
| 33Ser | 118.12 | 8.18 | 70Ala | 124.84 | 8.18 |
| 35Leu | 123.74 | 8.27 | 72Arg | 122.24 | 8.34 |
| 37Ser | 115.71 | 8.3 | 73Val | 125.53 | 7.71 |

Table S3. $^{15}$N-HSQC assignments for p53-TAD$^{N29K/N30D}$.

| P53-TAD$^{N29K/N30D}$ | | | | | |
|---|---|---|---|---|---|
| Residue | N (ppm) | HN (ppm) | Residue | N (ppm) | HN (ppm) |
| 1Met | 121.47 | 8.41 | 38Gln | 122.25 | 8.34 |
| 2Glu | 122.02 | 8.36 | 39Ala | 125.26 | 8.27 |
| 3Glu | 123.47 | 8.38 | 40Met | 119.56 | 8.3 |
| 5Gln | 120.97 | 8.52 | 41Asp | 121.03 | 8.22 |
| 6Ser | 117.73 | 8.34 | 42Asp | 120.23 | 8.2 |
| 7Asp | 123.65 | 8.4 | 43Leu | 121.75 | 8.04 |
| 9Ser | 116.02 | 8.48 | 44Met | 120.37 | 8.21 |
| 10Val | 121.09 | 7.83 | 45Leu | 123.17 | 8.05 |
| 11Glu | 126.14 | 8.3 | 46Ser | 118.46 | 8.48 |
| 14Leu | 122.45 | 8.31 | 48Asp | 118.85 | 8.18 |
| 15Ser | 116.68 | 8.29 | 49Asp | 120.23 | 8.06 |
| 16Gln | 122.49 | 8.44 | 50Ile | 120.2 | 7.81 |
| 17Glu | 121.93 | 8.43 | 51Glu | 123.76 | 8.3 |
| 18Thr | 114.94 | 8.07 | 52Gln | 120.44 | 8.1 |
| 19Phe | 122.37 | 8.22 | 53Trp | 121.53 | 7.94 |
| 20Ser | 116.75 | 8.05 | 54Phe | 121.53 | 7.94 |
| 21Asp | 122.17 | 8.23 | 55Thr | 116.32 | 7.95 |
| 22Leu | 121.24 | 7.9 | 56Glu | 123.37 | 8.26 |
| 23Trp | 119.24 | 7.8 | 57Asp | 123.43 | 8.4 |
| 24Lys | 120.93 | 7.54 | 59Gly | 109.33 | 8.36 |
| 25Leu | 121.43 | 7.84 | 61Asp | 119.89 | 8.41 |
| 26Leu | 123.98 | 7.97 | 62Glu | 120.85 | 8.07 |
| 28Glu | 121.01 | 8.45 | 63Ala | 126.62 | 8.23 |
| 29Lys | 121.38 | 8.31 | 65Arg | 121.93 | 8.43 |
| 30Asp | 121.56 | 8.24 | 66Met | 123.32 | 8.44 |
| 31Val | 120.03 | 7.98 | 68Glu | 121.23 | 8.5 |
| 32Leu | 125.36 | 8.26 | 69Ala | 125.39 | 8.27 |
| 33Ser | 118.21 | 8.18 | 70Ala | 124.84 | 8.18 |
| 35Leu | 123.77 | 8.27 | 72Arg | 122.49 | 8.44 |
| 37Ser | 115.74 | 8.31 | 73Val | 125.54 | 7.71 |

**Table S4. $^{15}$N-HSQC assignments for p53-TAD$^{D49Y}$.**

| p53-TAD$^{D49Y}$ | | | | | |
|---|---|---|---|---|---|
| **Residue** | **N (ppm)** | **HN (ppm)** | **Residue** | **N (ppm)** | **HN (ppm)** |
| **1Met** | 121.48 | 8.4 | **38Gln** | 122.19 | 8.34 |
| **2Glu** | 122.03 | 8.36 | **39Ala** | 125.07 | 8.26 |
| **3Glu** | 123.48 | 8.38 | **40Met** | 119.51 | 8.28 |
| **5Gln** | 120.97 | 8.52 | **41Asp** | 120.93 | 8.2 |
| **6Ser** | 117.71 | 8.34 | **42Asp** | 120.11 | 8.17 |
| **7Asp** | 123.64 | 8.4 | **43Leu** | 122 | 8.06 |
| **9Ser** | 116.02 | 8.49 | **44Met** | 119.2 | 8.12 |
| **10Val** | 121.08 | 7.83 | **45Leu** | 123.24 | 7.69 |
| **11Glu** | 126.14 | 8.3 | **46Ser** | 118.37 | 8.18 |
| **14Leu** | 122.43 | 8.31 | **48Asp** | 117.62 | 8.13 |
| **15Ser** | 116.68 | 8.29 | **49Tyr** | 121.89 | 7.91 |
| **16Gln** | 122.49 | 8.44 | **50Ile** | 120.37 | 7.81 |
| **17Glu** | 121.93 | 8.43 | **51Glu** | 124.05 | 8.2 |
| **18Thr** | 114.91 | 8.07 | **52Gln** | 120.72 | 8.07 |
| **19Phe** | 122.29 | 8.2 | **53Trp** | 121.35 | 7.93 |
| **20Ser** | 116.72 | 8.06 | **54Phe** | 121.37 | 7.94 |
| **21Asp** | 122.1 | 8.24 | **55Thr** | 116.09 | 7.94 |
| **22Leu** | 121.13 | 7.9 | **56Glu** | 123.27 | 8.26 |
| **23Trp** | 119.36 | 7.81 | **57Asp** | 123.32 | 8.39 |
| **24Lys** | 120.46 | 7.54 | **59Gly** | 109.33 | 8.36 |
| **25Leu** | 120.72 | 7.78 | **61Asp** | 119.84 | 8.41 |
| **26Leu** | 123.67 | 7.87 | **62Glu** | 120.81 | 8.07 |
| **28Glu** | 119.78 | 8.67 | **63Ala** | 126.59 | 8.23 |
| **29Asn** | 118.83 | 8.24 | **65Arg** | 121.9 | 8.43 |
| **30Asn** | 119.56 | 8.25 | **66Met** | 123.3 | 8.44 |
| **31Val** | 120.17 | 7.99 | **68Glu** | 121.21 | 8.5 |
| **32Leu** | 125.49 | 8.25 | **69Ala** | 125.38 | 8.27 |
| **33Ser** | 118.07 | 8.18 | **70Ala** | 124.83 | 8.18 |
| **35Leu** | 123.7 | 8.26 | **72Arg** | 122.48 | 8.44 |
| **37Ser** | 115.65 | 8.3 | **73Val** | 125.52 | 7.71 |

**Table S5. $^{15}$N-HSQC assignments for p53-TAD$^{W53G}$.**

| p53-TAD$^{W53G}$ | | | | | |
|---|---|---|---|---|---|
| Residue | N (ppm) | HN (ppm) | Residue | N (ppm) | HN (ppm) |
| 1Met | 121.5 | 8.4 | 38Gln | 122.26 | 8.34 |
| 2Glu | 122.04 | 8.35 | 39Ala | 125.27 | 8.28 |
| 3Glu | 123.49 | 8.38 | 40Met | 119.6 | 8.3 |
| 5Gln | 120.98 | 8.52 | 41Asp | 121.08 | 8.22 |
| 6Ser | 117.72 | 8.34 | 42Asp | 120.28 | 8.2 |
| 7Asp | 123.65 | 8.39 | 43Leu | 121.78 | 8.05 |
| 9Ser | 116.03 | 8.48 | 44Met | 120.42 | 8.21 |
| 10Val | 121.09 | 7.82 | 45Leu | 123.24 | 8.06 |
| 11Glu | 126.15 | 8.3 | 46Ser | 118.51 | 8.48 |
| 14Leu | 122.44 | 8.3 | 48Asp | 119.06 | 8.19 |
| 15Ser | 116.68 | 8.29 | 49Asp | 120.25 | 8.07 |
| 16Gln | 122.49 | 8.44 | 50Ile | 120.42 | 7.86 |
| 17Glu | 121.93 | 8.42 | 51Glu | 124.48 | 8.37 |
| 18Thr | 114.93 | 8.06 | 52Gln | 121.55 | 8.33 |
| 19Phe | 122.31 | 8.19 | 53Gly | 109.66 | 8.32 |
| 20Ser | 116.75 | 8.06 | 54Phe | - | - |
| 21Asp | 122.13 | 8.24 | 55Thr | 117.16 | 8.07 |
| 22Leu | 121.14 | 7.9 | 56Glu | 123.61 | 8.31 |
| 23Trp | 119.4 | 7.81 | 57Asp | 123.58 | 8.42 |
| 24Lys | 120.52 | 7.54 | 59Gly | 109.33 | 8.36 |
| 25Leu | 120.76 | 7.78 | 61Asp | 119.97 | 8.41 |
| 26Leu | 123.73 | 7.87 | 62Glu | 120.92 | 8.08 |
| 28Glu | 119.81 | 8.67 | 63Ala | 126.65 | 8.23 |
| 29Asn | 118.86 | 8.24 | 65Arg | 121.9 | 8.42 |
| 30Asn | 119.59 | 8.24 | 66Met | 123.38 | 8.45 |
| 31Val | 120.23 | 7.99 | 68Glu | 121.26 | 8.5 |
| 32Leu | 125.58 | 8.25 | 69Ala | 125.42 | 8.27 |
| 33Ser | 118.13 | 8.18 | 70Ala | 124.87 | 8.19 |
| 35Leu | 123.78 | 8.27 | 72Arg | 122.47 | 8.44 |
| 37Ser | 115.73 | 8.3 | 73Val | 125.54 | 7.71 |

#### Table S6. $T_1$ relaxation for p53-TAD$^{WT}$.

| Assign F2 | Time Constant | TC error | Fit error | Number Peaks | Function | Fit Parameter A | Fit parameter B | Error Paramater A | Error Parameter B | (1/s) R1 | (1/s) R1 Error |
|---|---|---|---|---|---|---|---|---|---|---|---|
| 1MetN | 529.7195 | 87.29158 | 1.45459 | 10 | A exp(-Bx) | 39566.8 | 0.00189 | 2600.43579 | 0.000303068 | 2600435.79 | 0.303068 |
| 2GluN | 590.47655 | 43.79016 | 0.91594 | 10 | A exp(-Bx) | 53569.8 | 0.00169 | 1589.54907 | 0.000124911 | 1589549.07 | 0.124911 |
| 3GluN | 616.12102 | 42.82463 | 1.36508 | 10 | A exp(-Bx) | 84750 | 0.00162 | 2341.96265 | 0.000112274 | 2341962.65 | 0.112274 |
| 5GlnN | 571.27924 | 49.80767 | 1.02228 | 10 | A exp(-Bx) | 51145.2 | 0.00175 | 1790.09424 | 0.000151473 | 1790094.24 | 0.151473 |
| 6SerN | 454.39542 | 60.35737 | 1.22984 | 10 | A exp(-Bx) | 41930.4 | 0.0022 | 2294.80029 | 0.000287339 | 2294800.29 | 0.287339 |
| 7AspN | 500.92293 | 44.61594 | 1.13272 | 10 | A exp(-Bx) | 56352.5 | 0.002 | 2056.62012 | 0.000176418 | 2056620.12 | 0.176418 |
| 9SerN | 434.05601 | 38.87684 | 1.00753 | 10 | A exp(-Bx) | 50786.4 | 0.0023 | 1904.55359 | 0.000204718 | 1904553.59 | 0.204718 |
| 10ValN | 534.14078 | 28.68708 | 0.78061 | 10 | A exp(-Bx) | 63591.8 | 0.00187 | 1392.33215 | 0.00010026 | 1392332.15 | 0.10026 |
| 11GluN | 624.6789 | 36.45155 | 0.755 | 10 | A exp(-Bx) | 55676.7 | 0.0016 | 1290.48804 | 0.000093096 | 1290488.04 | 0.093096 |
| 14LeuN | 593.74398 | 31.25202 | 0.78413 | 10 | A exp(-Bx) | 64402.9 | 0.00168 | 1358.75623 | 8.84059E-05 | 1358756.23 | 0.0884059 |
| 15SerN | 483.50692 | 46.07591 | 0.91351 | 10 | A exp(-Bx) | 42712.7 | 0.00207 | 1675.0719 | 0.000195334 | 1675071.9 | 0.195334 |
| 16GlnN | 598.13721 | 63.83366 | 1.99326 | 10 | A exp(-Bx) | 81365.2 | 0.00167 | 3447.09131 | 0.000176435 | 3447091.31 | 0.176435 |
| 17GluN | 471.14883 | 44.13762 | 1.98212 | 10 | A exp(-Bx) | 94558 | 0.00212 | 3660.99854 | 0.00019712 | 3660998.54 | 0.19712 |
| 18ThrN | 424.40112 | 47.01177 | 1.23604 | 10 | A exp(-Bx) | 50743.1 | 0.00236 | 2351.52856 | 0.000257881 | 2351528.56 | 0.257881 |
| 19PheN | 403.18201 | 39.97622 | 0.8666 | 10 | A exp(-Bx) | 39943.6 | 0.00248 | 1673.13757 | 0.000243552 | 1673137.57 | 0.243552 |
| 20SerN | 377.6844 | 40.08395 | 0.75165 | 10 | A exp(-Bx) | 32730.5 | 0.00265 | 1478.97083 | 0.000277908 | 1478970.83 | 0.277908 |
| 21AspN | 372.18391 | 29.52968 | 0.62087 | 10 | A exp(-Bx) | 36071.4 | 0.00269 | 1226.901 | 0.000211853 | 1226901 | 0.211853 |
| 22LeuN | 453.85326 | 33.54258 | 1.22913 | 10 | A exp(-Bx) | 74446.2 | 0.0022 | 2294.21997 | 0.000161962 | 2294219.97 | 0.161962 |
| 23TrpN | 457.69119 | 21.77217 | 0.64513 | 10 | A exp(-Bx) | 60449.8 | 0.00218 | 1201.32385 | 0.0001037 | 1201323.85 | 0.1037 |
| 24LysN | 430.75717 | 36.46475 | 1.20428 | 10 | A exp(-Bx) | 64240 | 0.00232 | 2281.38306 | 0.000195132 | 2281383.06 | 0.195132 |
| 25LeuN | 500.31345 | 37.05282 | 0.83284 | 10 | A exp(-Bx) | 49717.8 | 0.002 | 1512.64673 | 0.000147223 | 1512646.73 | 0.147223 |
| 26LeuN | 523.75121 | 15.80886 | 0.47309 | 10 | A exp(-Bx) | 68599.7 | 0.00191 | 848.41309 | 5.75778E-05 | 848413.09 | 0.0575778 |
| 28GluN | 492.65704 | 38.71398 | 1.10699 | 10 | A exp(-Bx) | 62441.7 | 0.00203 | 2019.22632 | 0.000158534 | 2019226.32 | 0.158534 |
| 29AsnN | 409.84538 | 40.30215 | 0.75555 | 10 | A exp(-Bx) | 35025.4 | 0.00244 | 1451.85791 | 0.000237656 | 1451857.91 | 0.237656 |
| 30AsnN | 393.3032 | 47.49229 | 0.75347 | 10 | A exp(-Bx) | 28752.3 | 0.00254 | 1465.17688 | 0.00030267 | 1465176.88 | 0.30267 |
| 31ValN | 508.80859 | 35.80639 | 0.94621 | 10 | A exp(-Bx) | 59291.1 | 0.00197 | 1710.53906 | 0.000137631 | 1710539.06 | 0.137631 |
| 32LeuN | 556.55335 | 40.59134 | 0.99355 | 10 | A exp(-Bx) | 59462.7 | 0.0018 | 1752.2417 | 0.000130355 | 1752241.7 | 0.130355 |
| 33SerN | 543.90706 | 54.803 | 0.9755 | 10 | A exp(-Bx) | 42572 | 0.00184 | 1731.29272 | 0.000183405 | 1731292.72 | 0.183405 |
| 35LeuN | 640.30087 | 26.84963 | 0.65881 | 10 | A exp(-Bx) | 67333.9 | 0.00156 | 1118.67944 | 6.53745E-05 | 1118679.44 | 0.0653745 |
| 37SerN | 479.62235 | 49.9691 | 0.87517 | 10 | A exp(-Bx) | 37531.4 | 0.00208 | 1608.38501 | 0.000214913 | 1608385.01 | 0.214913 |
| 38GlnN | 438.87321 | 44.67999 | 0.79521 | 10 | A exp(-Bx) | 35292.1 | 0.00228 | 1498.49231 | 0.000229616 | 1498492.31 | 0.229616 |
| 39AlaN | 495.61339 | 44.61729 | 1.03768 | 10 | A exp(-Bx) | 51151.6 | 0.00202 | 1889.66541 | 0.000180194 | 1889665.41 | 0.180194 |
| 40MetN | 514.20064 | 44.73301 | 1.01165 | 10 | A exp(-Bx) | 51345.7 | 0.00194 | 1823.50647 | 0.000167924 | 1823506.47 | 0.167924 |
| 41AspN | 539.91346 | 43.44431 | 1.0959 | 10 | A exp(-Bx) | 59724.4 | 0.00185 | 1948.92151 | 0.000148081 | 1948921.51 | 0.148081 |
| 42AspN | 549.66572 | 34.79131 | 1.08852 | 10 | A exp(-Bx) | 75076.2 | 0.00182 | 1926.30603 | 0.000114695 | 1926306.03 | 0.114695 |
| 43LeuN | 582.10384 | 38.35729 | 0.92927 | 10 | A exp(-Bx) | 61123.6 | 0.00172 | 1617.13135 | 0.000112713 | 1617131.35 | 0.112713 |
| 44MetN | 532.30342 | 47.42483 | 1.64333 | 10 | A exp(-Bx) | 81135.6 | 0.00188 | 2933.90723 | 0.000166066 | 2933907.23 | 0.166066 |
| 45LeuN | 586.87373 | 31.29549 | 0.73839 | 10 | A exp(-Bx) | 59940.4 | 0.0017 | 1283.54395 | 9.06072E-05 | 1283543.95 | 0.0906072 |
| 46SerN | 539.92503 | 36.14529 | 0.94138 | 10 | A exp(-Bx) | 61543.8 | 0.00185 | 1674.11804 | 0.000123439 | 1674118.04 | 0.123439 |
| 48AspN | 514.82075 | 35.45106 | 0.91086 | 10 | A exp(-Bx) | 58234.4 | 0.00194 | 1641.28418 | 0.000133129 | 1641284.18 | 0.133129 |
| 49AspN | 503.8615 | 42.09421 | 1.14872 | 10 | A exp(-Bx) | 60825.9 | 0.00198 | 2082.2478 | 0.000164665 | 2082247.8 | 0.164665 |
| 50IleN | 573.07446 | 32.76447 | 0.9608 | 10 | A exp(-Bx) | 72965.7 | 0.00174 | 1680.98792 | 9.94417E-05 | 1680987.92 | 0.0994417 |
| 51GluN | 551.98672 | 29.08977 | 0.85003 | 10 | A exp(-Bx) | 70295 | 0.00181 | 1502.52942 | 9.52098E-05 | 1502529.42 | 0.0952098 |
| 52GlnN | 508.63493 | 31.06562 | 0.87658 | 10 | A exp(-Bx) | 63214.7 | 0.00197 | 1584.80798 | 0.000119635 | 1584807.98 | 0.119635 |
| 53TrpN | 462.81242 | 50.49004 | 2.6312 | 10 | A exp(-Bx) | 108366 | 0.00216 | 4884.30615 | 0.000232979 | 4884306.15 | 0.232979 |
| 55ThrN | 476.39332 | 51.39203 | 1.25053 | 10 | A exp(-Bx) | 51876.6 | 0.0021 | 2302.57544 | 0.00022387 | 2302575.44 | 0.22387 |
| 56GluN | 513.16791 | 42.58486 | 1.14794 | 10 | A exp(-Bx) | 61053.2 | 0.00195 | 2070.32153 | 0.000160611 | 2070321.53 | 0.160611 |
| 57AspN | 553.41084 | 34.747 | 1.0416 | 10 | A exp(-Bx) | 72361.2 | 0.00181 | 1839.84802 | 0.000113011 | 1839848.02 | 0.113011 |
| 59GlyN | 595.87439 | 26.24381 | 0.68787 | 10 | A exp(-Bx) | 67437.5 | 0.00168 | 1190.78589 | 7.37696E-05 | 1190785.89 | 0.0737696 |
| 61AspN | 503.37266 | 43.74795 | 1.15533 | 10 | A exp(-Bx) | 58847.3 | 0.00199 | 2094.83008 | 0.00017137 | 2094830.08 | 0.17137 |
| 62GluN | 556.61412 | 31.13079 | 0.74197 | 10 | A exp(-Bx) | 57781.2 | 0.0018 | 1308.51025 | 0.000100168 | 1308510.25 | 0.100168 |
| 63AlaN | 575.65479 | 46.57795 | 1.15266 | 10 | A exp(-Bx) | 62022.8 | 0.00174 | 2014.20618 | 0.00013965 | 2014206.18 | 0.13965 |
| 65ArgN | 471.14883 | 44.13762 | 1.98212 | 10 | A exp(-Bx) | 94558 | 0.00212 | 3660.99854 | 0.00019712 | 3660998.54 | 0.19712 |
| 66MetN | 490.89829 | 43.26331 | 0.89059 | 10 | A exp(-Bx) | 44882.7 | 0.00204 | 1626.13354 | 0.000178157 | 1626133.54 | 0.178157 |
| 68GluN | 567.7597 | 46.10141 | 1.02303 | 10 | A exp(-Bx) | 54940.4 | 0.00176 | 1794.43103 | 0.000142085 | 1794431.03 | 0.142085 |
| 69AlaN | 554.62498 | 57.68256 | 1.09277 | 10 | A exp(-Bx) | 46128.6 | 0.0018 | 1929.07263 | 0.000185534 | 1929072.63 | 0.185534 |
| 70AlaN | 736.51304 | 51.38667 | 1.01133 | 10 | A exp(-Bx) | 61586 | 0.00136 | 1655.5199 | 9.42737E-05 | 1655519.9 | 0.0942737 |
| 72ArgN | 598.13721 | 63.83366 | 1.99326 | 10 | A exp(-Bx) | 81365.2 | 0.00167 | 3447.09131 | 0.000176435 | 3447091.31 | 0.176435 |
| 73ValN | 1177.65832 | 62.14104 | 1.18782 | 10 | A exp(-Bx) | 96081.9 | 0.000849143 | 1746.16272 | 4.46823E-05 | 1746162.72 | 0.0446823 |

#### Table S7. $T_1$ relaxation for p53-TAD$^{K24N}$.

| Assign F1 | Time Constant | TC Error | Fit Error | Num Peaks | Function | Fit Param A | Fit Param B | Param Error A | Param Error B | (1/s) R1 | (1/s) R1 Errror |
|---|---|---|---|---|---|---|---|---|---|---|---|
| 1MetH* | 547.06914 | 54.18248 | 4.03786 | 10 | A exp(-Bx) | 27140.4 | 0.00183 | 1084.28479 | 0.000179298 | 1.83 | 0.179298 |
| 2GluH | 540.45368 | 64.73783 | 6.0333 | 10 | A exp(-Bx) | 33728.8 | 0.00185 | 1625.73047 | 0.000218544 | 1.85 | 0.218544 |
| 3GluH | 545.02358 | 50.49101 | 6.51471 | 10 | A exp(-Bx) | 46783.5 | 0.00183 | 1751.37463 | 0.00016854 | 1.83 | 0.16854 |
| 5GlnH | 525.01043 | 53.77213 | 5.0302 | 10 | A exp(-Bx) | 32878 | 0.0019 | 1366.31201 | 0.000193079 | 1.9 | 0.193079 |
| 6SerH | 542.31908 | 46.2551 | 3.20384 | 10 | A exp(-Bx) | 24972.5 | 0.00184 | 862.48822 | 0.000156144 | 1.84 | 0.156144 |
| 7AspH | 510.79951 | 33.37579 | 2.8202 | 10 | A exp(-Bx) | 28781 | 0.00196 | 770.99768 | 0.000127376 | 1.96 | 0.127376 |
| 9SerH | 453.39658 | 25.74414 | 2.72379 | 10 | A exp(-Bx) | 32477.9 | 0.00221 | 770.76868 | 0.000124833 | 2.21 | 0.124833 |
| 10ValH | 504.08527 | 36.48244 | 7.28233 | 10 | A exp(-Bx) | 67344.2 | 0.00198 | 2000.44104 | 0.00014283 | 1.98 | 0.14283 |
| 11GluH | 560.92524 | 50.70489 | 4.76321 | 10 | A exp(-Bx) | 34926.6 | 0.00178 | 1270.47009 | 0.000159858 | 1.78 | 0.159858 |
| 14LeuH | 605.95864 | 56.6958 | 5.19857 | 10 | A exp(-Bx) | 36541.6 | 0.00165 | 1357.80994 | 0.000153078 | 1.65 | 0.153078 |
| 15SerH | 533.37693 | 33.22122 | 2.51299 | 10 | A exp(-Bx) | 26785 | 0.00187 | 679.60254 | 0.000116325 | 1.87 | 0.116325 |
| 16GlnH | 615.36587 | 76.25253 | 9.85755 | 10 | A exp(-Bx) | 52569.3 | 0.00163 | 2563.99854 | 0.000198366 | 1.63 | 0.198366 |
| 17GluH | 497.50928 | 43.69549 | 4.49116 | 10 | A exp(-Bx) | 34363.3 | 0.00201 | 1238.24207 | 0.000175195 | 2.01 | 0.175195 |
| 18ThrH | 514.51686 | 34.05908 | 3.38334 | 10 | A exp(-Bx) | 34093.6 | 0.00194 | 924.12689 | 0.000128098 | 1.94 | 0.128098 |
| 19PheH | 464.79322 | 38.02673 | 3.48659 | 10 | A exp(-Bx) | 28854.3 | 0.00215 | 979.64221 | 0.00017486 | 2.15 | 0.17486 |
| 20SerH | 441.9434 | 26.5702 | 2.49743 | 10 | A exp(-Bx) | 28231.7 | 0.00226 | 711.85321 | 0.00013555 | 2.26 | 0.13555 |
| 21AspH | 451.95391 | 21.44603 | 2.056 | 10 | A exp(-Bx) | 29316.1 | 0.00221 | 582.26569 | 0.000104757 | 2.21 | 0.104757 |
| 22LeuH | 522.75547 | 42.71309 | 5.10914 | 10 | A exp(-Bx) | 41725.7 | 0.00191 | 1389.40649 | 0.000155272 | 1.91 | 0.155272 |
| 23TrpH | 501.47557 | 32.85673 | 3.5151 | 10 | A exp(-Bx) | 35895.9 | 0.00199 | 966.98596 | 0.000130099 | 1.99 | 0.130099 |
| 24AsnH | 394.54055 | 29.04628 | 2.93354 | 10 | A exp(-Bx) | 27565.8 | 0.00253 | 863.6972 | 0.000185598 | 2.53 | 0.185598 |
| 26LeuH | 578.00617 | 48.72791 | 5.07829 | 10 | A exp(-Bx) | 39753.5 | 0.00173 | 1343.47314 | 0.00014483 | 1.73 | 0.14483 |
| 28GluH | 488.63984 | 34.45981 | 4.24994 | 10 | A exp(-Bx) | 40479.6 | 0.00205 | 1177.62781 | 0.000143612 | 2.05 | 0.143612 |
| 29AsnH | 476.16251 | 33.68981 | 2.41848 | 10 | A exp(-Bx) | 23036.3 | 0.0021 | 675.01202 | 0.000147853 | 2.1 | 0.147853 |
| 30AsnH | 429.05347 | 26.0081 | 2.02159 | 10 | A exp(-Bx) | 22503.8 | 0.00233 | 574.57922 | 0.000140766 | 2.33 | 0.140766 |
| 31ValH | 538.72463 | 39.42739 | 4.45619 | 10 | A exp(-Bx) | 40434 | 0.00186 | 1201.82373 | 0.000135132 | 1.86 | 0.135132 |
| 32LeuH | 525.60841 | 46.38074 | 4.03699 | 10 | A exp(-Bx) | 30541.2 | 0.0019 | 1096.17847 | 0.000166598 | 1.9 | 0.166598 |
| 33SerH | 545.69279 | 48.25049 | 3.58501 | 10 | A exp(-Bx) | 26949.6 | 0.00183 | 963.45813 | 0.000160786 | 1.83 | 0.160786 |
| 35LeuH | 558.23538 | 50.80131 | 5.82405 | 10 | A exp(-Bx) | 42442.7 | 0.00179 | 1555.29236 | 0.000161692 | 1.79 | 0.161692 |
| 37SerH | 512.36727 | 47.6237 | 3.37412 | 10 | A exp(-Bx) | 24328.7 | 0.00195 | 922.69025 | 0.000179869 | 1.95 | 0.179869 |
| 38GlnH | 458.49229 | 21.24087 | 1.74958 | 10 | A exp(-Bx) | 25501.1 | 0.00218 | 493.48233 | 0.000100828 | 2.18 | 0.100828 |
| 39AlaH | 548.2018 | 46.43483 | 4.31353 | 10 | A exp(-Bx) | 33809.8 | 0.00182 | 1157.78467 | 0.000153419 | 1.82 | 0.153419 |
| 40MetH | 529.57551 | 42.47851 | 4.00291 | 10 | A exp(-Bx) | 33241.1 | 0.00189 | 1084.67004 | 0.000150503 | 1.89 | 0.150503 |
| 41AspH | 531.10405 | 39.82164 | 4.00277 | 10 | A exp(-Bx) | 35519.6 | 0.00188 | 1083.76904 | 0.000140391 | 1.88 | 0.140391 |
| 42AspH | 540.64686 | 45.68949 | 5.6525 | 10 | A exp(-Bx) | 44476.2 | 0.00185 | 1522.95386 | 0.00015521 | 1.85 | 0.15521 |
| 43LeuH | 594.21858 | 58.10668 | 5.64474 | 10 | A exp(-Bx) | 38072.5 | 0.00168 | 1482.16138 | 0.000163019 | 1.68 | 0.163019 |
| 44MetH | 554.73709 | 52.84353 | 6.31818 | 10 | A exp(-Bx) | 44057.1 | 0.0018 | 1690.34363 | 0.000170188 | 1.8 | 0.170188 |
| 45LeuH | 534.40999 | 51.41271 | 5.47272 | 10 | A exp(-Bx) | 37536.2 | 0.00187 | 1462.86841 | 0.000178384 | 1.87 | 0.178384 |
| 46SerH | 552.55 | 47.21307 | 4.54203 | 10 | A exp(-Bx) | 35263.7 | 0.00181 | 1216.46277 | 0.000153526 | 1.81 | 0.153526 |
| 48AspH | 533.13358 | 42.56285 | 4.33045 | 10 | A exp(-Bx) | 36096.7 | 0.00188 | 1171.13843 | 0.000148805 | 1.88 | 0.148805 |
| 49AspH | 563.28434 | 50.11067 | 5.15648 | 10 | A exp(-Bx) | 38391.9 | 0.00178 | 1373.79016 | 0.000156703 | 1.78 | 0.156703 |
| 50IleH | 565.38211 | 47.00129 | 4.88157 | 10 | A exp(-Bx) | 38839.5 | 0.00177 | 1299.22839 | 0.000146034 | 1.77 | 0.146034 |
| 51GluH | 513.57505 | 46.08536 | 5.58279 | 10 | A exp(-Bx) | 41660 | 0.00195 | 1525.67871 | 0.00017334 | 1.95 | 0.17334 |
| 52GlnH | 510.22499 | 38.76602 | 4.5207 | 10 | A exp(-Bx) | 39784.8 | 0.00196 | 1237.67126 | 0.000148061 | 1.96 | 0.148061 |
| 53TrpH | 524.29341 | 40.07616 | 7.63661 | 10 | A exp(-Bx) | 66581 | 0.00191 | 2074.81885 | 0.000144951 | 1.91 | 0.144951 |
| 55ThrH | 489.29058 | 28.94915 | 2.95452 | 10 | A exp(-Bx) | 33450.2 | 0.00204 | 817.4455 | 0.000120501 | 2.04 | 0.120501 |
| 56GluH | 527.61185 | 39.99741 | 4.0111 | 10 | A exp(-Bx) | 35235.9 | 0.0019 | 1088.00476 | 0.000142866 | 1.9 | 0.142866 |
| 59GlyH | 541.56424 | 46.69497 | 4.67118 | 10 | A exp(-Bx) | 36028 | 0.00185 | 1257.98486 | 0.000158044 | 1.85 | 0.158044 |
| 61AspH | 492.97958 | 34.41928 | 3.60252 | 10 | A exp(-Bx) | 34617 | 0.00203 | 995.77454 | 0.000140943 | 2.03 | 0.140943 |
| 62GluH | 559.84674 | 50.52135 | 4.78664 | 10 | A exp(-Bx) | 35164.5 | 0.00179 | 1277.37744 | 0.000159898 | 1.79 | 0.159898 |
| 63AlaH | 516.99728 | 36.53268 | 4.32535 | 10 | A exp(-Bx) | 40786.2 | 0.00193 | 1178.53674 | 0.000136004 | 1.93 | 0.136004 |
| 65ArgH | 564.43647 | 53.68244 | 4.96217 | 10 | A exp(-Bx) | 34587.4 | 0.00177 | 1321.26782 | 0.000167004 | 1.77 | 0.167004 |
| 66MetH | 574.99287 | 61.99575 | 4.04544 | 10 | A exp(-Bx) | 24881.6 | 0.00174 | 1071.63306 | 0.000185385 | 1.74 | 0.185385 |
| 68GluH | 521.17454 | 46.03009 | 5.02901 | 10 | A exp(-Bx) | 38051.9 | 0.00192 | 1368.76147 | 0.000168162 | 1.92 | 0.168162 |
| 69AlaH | 523.66324 | 98.29397 | 10.39626 | 10 | A exp(-Bx) | 37958.2 | 0.00191 | 2825.82739 | 0.000346635 | 1.91 | 0.346635 |
| 70AlaH | 588.84074 | 66.65063 | 7.55337 | 10 | A exp(-Bx) | 44194.6 | 0.0017 | 1988.20679 | 0.000189823 | 1.7 | 0.189823 |
| 73ValH | 794.06881 | 214.48802 | 21.03358 | 10 | A exp(-Bx) | 53071.6 | 0.00126 | 5120.26904 | 0.000318416 | 1.26 | 0.318416 |

#### Table S8. $T_1$ relaxation for p53-TAD$^{N29K/N30D}$.

| Assign F1 | Time Constant | TC Error | Fit Error | Num Peaks | Function | Fit Param A | Fit Param B | Param Error A | Param Error B | (1/s) R1 | (1/s) R1 error |
|---|---|---|---|---|---|---|---|---|---|---|---|
| 1MetH* | 546.13389 | 14.90432 | 1.14828 | 10 | A exp(-Bx) | 3.92E+04 | 0.00183 | 435.71307 | 4.99E-05 | 1.83 | 4.99E-02 |
| 2GluH | 580.47841 | 15.40035 | 1.36107 | 10 | A exp(-Bx) | 4.75E+04 | 0.00172 | 507.92233 | 4.57E-05 | 1.72 | 4.57E-02 |
| 3GluH | 651.75959 | 20.34815 | 2.00367 | 10 | A exp(-Bx) | 5.88E+04 | 0.00153 | 725.61273 | 4.79E-05 | 1.53 | 4.79E-02 |
| 5GlnH | 544.39941 | 23.35897 | 2.2337 | 10 | A exp(-Bx) | 4.86E+04 | 0.00184 | 849.17615 | 7.87E-05 | 1.84 | 7.87E-02 |
| 6SerH | 539.00179 | 16.04444 | 1.14534 | 10 | A exp(-Bx) | 3.60E+04 | 0.00186 | 436.66174 | 5.52E-05 | 1.86 | 5.52E-02 |
| 7AspH | 595.1324 | 29.05969 | 1.82731 | 10 | A exp(-Bx) | 3.46E+04 | 0.00168 | 678.07642 | 8.19E-05 | 1.68 | 8.19E-02 |
| 9SerH | 470.46671 | 10.73999 | 1.02343 | 10 | A exp(-Bx) | 4.25E+04 | 0.00213 | 404.76096 | 4.85E-05 | 2.13 | 4.85E-02 |
| 10ValH | 591.78978 | 16.82152 | 1.47264 | 10 | A exp(-Bx) | 4.79E+04 | 0.00169 | 547.23822 | 4.80E-05 | 1.69 | 4.80E-02 |
| 11GluH | 599.02291 | 18.60029 | 1.60863 | 10 | A exp(-Bx) | 4.79E+04 | 0.00167 | 595.8797 | 5.18E-05 | 1.67 | 5.18E-02 |
| 14LeuH | 603.31967 | 13.54836 | 1.18812 | 10 | A exp(-Bx) | 4.88E+04 | 0.00166 | 439.25732 | 3.72E-05 | 1.66 | 3.72E-02 |
| 15SerH | 527.13435 | 17.05979 | 1.21434 | 10 | A exp(-Bx) | 3.52E+04 | 0.0019 | 465.81763 | 6.13E-05 | 1.9 | 6.13E-02 |
| 18ThrH | 484.21321 | 17.35513 | 1.67344 | 10 | A exp(-Bx) | 4.42E+04 | 0.00207 | 657.25812 | 7.39E-05 | 2.07 | 7.39E-02 |
| 19PheH | 451.00086 | 11.06902 | 0.93937 | 10 | A exp(-Bx) | 3.62E+04 | 0.00222 | 372.18317 | 5.44E-05 | 2.22 | 5.44E-02 |
| 20SerH | 465.58002 | 7.54244 | 0.48198 | 10 | A exp(-Bx) | 2.80E+04 | 0.00215 | 189.2616 | 3.48E-05 | 2.15 | 3.48E-02 |
| 21AspH | 406.484 | 25.26432 | 1.80798 | 10 | A exp(-Bx) | 2.83E+04 | 0.00246 | 746.14154 | 1.52E-04 | 2.46 | 1.52E-01 |
| 22LeuH | 505.90825 | 13.67335 | 1.76387 | 10 | A exp(-Bx) | 6.15E+04 | 0.00198 | 684.35645 | 5.34E-05 | 1.98 | 5.34E-02 |
| 23TrpH | 480.57256 | 10.8251 | 1.11849 | 10 | A exp(-Bx) | 4.70E+04 | 0.00208 | 440.17465 | 4.68E-05 | 2.08 | 4.68E-02 |
| 24LysH | 469.6814 | 22.68529 | 2.67271 | 10 | A exp(-Bx) | 5.26E+04 | 0.00213 | 1057.53162 | 1.03E-04 | 2.13 | 1.03E-01 |
| 25LeuH | 517.7273 | 11.90932 | 1.24534 | 10 | A exp(-Bx) | 5.08E+04 | 0.00193 | 480.04224 | 4.44E-05 | 1.93 | 4.44E-02 |
| 26LeuH | 554.14303 | 9.34476 | 0.8933 | 10 | A exp(-Bx) | 4.93E+04 | 0.0018 | 337.9902 | 3.04E-05 | 1.8 | 3.04E-02 |
| 28GluH | 562.3781 | 21.33476 | 2.02485 | 10 | A exp(-Bx) | 4.97E+04 | 0.00178 | 763.04004 | 6.74E-05 | 1.78 | 6.74E-02 |
| 29LysH | 545.29874 | 9.64671 | 0.74778 | 10 | A exp(-Bx) | 3.94E+04 | 0.00183 | 283.86389 | 3.24E-05 | 1.83 | 3.24E-02 |
| 30AspH | 528.3923 | 17.18486 | 1.636 | 10 | A exp(-Bx) | 4.71E+04 | 0.00189 | 626.44781 | 6.15E-05 | 1.89 | 6.15E-02 |
| 31ValH | 596.83305 | 22.07434 | 2.23501 | 10 | A exp(-Bx) | 5.58E+04 | 0.00168 | 827.79962 | 6.19E-05 | 1.68 | 6.19E-02 |
| 32LeuH | 546.69593 | 14.91832 | 2.5585 | 10 | A exp(-Bx) | 8.75E+04 | 0.00183 | 971.53143 | 4.99E-05 | 1.83 | 4.99E-02 |
| 33SerH | 550.47289 | 24.5519 | 1.89148 | 10 | A exp(-Bx) | 3.96E+04 | 0.00182 | 716.96954 | 8.09E-05 | 1.82 | 8.09E-02 |
| 35LeuH | 665.59462 | 24.76056 | 1.47153 | 10 | A exp(-Bx) | 3.62E+04 | 0.0015 | 529.8949 | 5.58E-05 | 1.5 | 5.58E-02 |
| 37SerH | 535.35932 | 19.23392 | 1.24658 | 10 | A exp(-Bx) | 3.25E+04 | 0.00187 | 476.14374 | 6.70E-05 | 1.87 | 6.70E-02 |
| 38GlnH | 476.68423 | 11.68203 | 0.75794 | 10 | A exp(-Bx) | 2.93E+04 | 0.0021 | 298.99371 | 5.14E-05 | 2.1 | 5.14E-02 |
| 39AlaH | 586.22313 | 9.36245 | 1.16979 | 10 | A exp(-Bx) | 6.77E+04 | 0.00171 | 435.37677 | 2.72E-05 | 1.71 | 2.72E-02 |
| 40MetH | 568.18287 | 14.64128 | 1.11765 | 10 | A exp(-Bx) | 3.98E+04 | 0.00176 | 415.3613 | 4.53E-05 | 1.76 | 4.53E-02 |
| 41AspH | 547.86163 | 12.99415 | 1.17859 | 10 | A exp(-Bx) | 4.63E+04 | 0.00183 | 446.82852 | 4.33E-05 | 1.83 | 4.33E-02 |
| 42AspH | 495.84048 | 23.59982 | 3.53525 | 10 | A exp(-Bx) | 7.01E+04 | 0.00202 | 1377.776 | 9.58E-05 | 2.02 | 9.58E-02 |
| 43LeuH | 613.61843 | 21.17703 | 1.87882 | 10 | A exp(-Bx) | 5.02E+04 | 0.00163 | 691.4588 | 5.62E-05 | 1.63 | 5.62E-02 |
| 44MetH | 601.96584 | 16.22091 | 1.86854 | 10 | A exp(-Bx) | 6.40E+04 | 0.00166 | 691.16333 | 4.47E-05 | 1.66 | 4.47E-02 |
| 45LeuH | 601.74336 | 11.41742 | 1.07072 | 10 | A exp(-Bx) | 5.20E+04 | 0.00166 | 395.69504 | 3.15E-05 | 1.66 | 3.15E-02 |
| 46SerH | 576.67822 | 15.44901 | 1.47277 | 10 | A exp(-Bx) | 5.09E+04 | 0.00173 | 550.58832 | 4.64E-05 | 1.73 | 4.64E-02 |
| 48AspH | 566.76612 | 16.00619 | 1.45822 | 10 | A exp(-Bx) | 4.80E+04 | 0.00176 | 548.34216 | 4.98E-05 | 1.76 | 4.98E-02 |
| 49AspH | 582.00973 | 20.52884 | 1.87867 | 10 | A exp(-Bx) | 4.93E+04 | 0.00172 | 700.58002 | 6.05E-05 | 1.72 | 6.05E-02 |
| 50IleH | 632.32229 | 23.38862 | 2.21706 | 10 | A exp(-Bx) | 5.51E+04 | 0.00158 | 809.39087 | 5.84E-05 | 1.58 | 5.84E-02 |
| 51GluH | 572.68239 | 19.52352 | 1.74231 | 10 | A exp(-Bx) | 4.74E+04 | 0.00175 | 652.5899 | 5.95E-05 | 1.75 | 5.95E-02 |
| 52GlnH | 556.84714 | 8.8327 | 0.83671 | 10 | A exp(-Bx) | 4.90E+04 | 0.0018 | 315.80191 | 2.85E-05 | 1.8 | 2.85E-02 |
| 53TrpH | 534.91936 | 11.80736 | 2.281 | 10 | A exp(-Bx) | 9.65E+04 | 0.00187 | 870.47687 | 4.12E-05 | 1.87 | 4.12E-02 |
| 54PheH | 534.91936 | 11.80736 | 2.281 | 10 | A exp(-Bx) | 9.65E+04 | 0.00187 | 870.47687 | 4.12E-05 | 1.87 | 4.12E-02 |
| 55ThrH | 515.88844 | 10.8446 | 0.91917 | 10 | A exp(-Bx) | 4.11E+04 | 0.00194 | 354.66223 | 4.07E-05 | 1.94 | 4.07E-02 |
| 56GluH | 517.49827 | 13.33932 | 1.37483 | 10 | A exp(-Bx) | 5.00E+04 | 0.00193 | 529.48315 | 4.98E-05 | 1.93 | 4.98E-02 |
| 57AspH | 588.86767 | 34.10853 | 3.17228 | 10 | A exp(-Bx) | 5.08E+04 | 0.0017 | 1180.53528 | 9.80E-05 | 1.7 | 9.80E-02 |
| 59GlyH | 587.46931 | 19.01171 | 1.73301 | 10 | A exp(-Bx) | 4.96E+04 | 0.0017 | 645.27454 | 5.50E-05 | 1.7 | 5.50E-02 |
| 61AspH | 569.6347 | 21.58714 | 1.78164 | 10 | A exp(-Bx) | 4.37E+04 | 0.00176 | 669.04242 | 6.64E-05 | 1.76 | 6.64E-02 |
| 62GluH | 586.75302 | 23.93751 | 1.83407 | 10 | A exp(-Bx) | 4.17E+04 | 0.0017 | 683.1969 | 6.94E-05 | 1.7 | 6.94E-02 |
| 63AlaH | 585.31642 | 18.57846 | 1.74197 | 10 | A exp(-Bx) | 5.08E+04 | 0.00171 | 649.32489 | 5.42E-05 | 1.71 | 5.42E-02 |
| 65ArgH | 521.83125 | 15.85914 | 2.4116 | 10 | A exp(-Bx) | 7.45E+04 | 0.00192 | 927.66364 | 5.82E-05 | 1.92 | 5.82E-02 |
| 66MetH | 552.02787 | 34.46209 | 2.30034 | 10 | A exp(-Bx) | 3.44E+04 | 0.00181 | 871.26703 | 1.13E-04 | 1.81 | 1.13E-01 |
| 68GluH | 542.22371 | 13.70823 | 1.40169 | 10 | A exp(-Bx) | 5.18E+04 | 0.00184 | 533.46191 | 4.66E-05 | 1.84 | 4.66E-02 |
| 69AlaH | 516.6322 | 25.67052 | 4.617 | 10 | A exp(-Bx) | 8.74E+04 | 0.00194 | 1780.75745 | 9.59E-05 | 1.94 | 9.59E-02 |
| 70AlaH | 698.87868 | 22.73309 | 2.01135 | 10 | A exp(-Bx) | 5.63E+04 | 0.00143 | 715.07428 | 4.65E-05 | 1.43 | 4.65E-02 |
| 72ArgH | 631.29798 | 16.13939 | 1.77537 | 10 | A exp(-Bx) | 6.37E+04 | 0.00158 | 647.70551 | 4.05E-05 | 1.58 | 4.05E-02 |
| 73ValH | 1050.86193 | 49.26951 | 3.98679 | 10 | A exp(-Bx) | 7.68E+04 | 9.52E-04 | 1284.28198 | 4.45E-05 | 0.9516 | 4.45E-02 |

#### Table S9. $T_1$ relaxation for p53-TAD$^{D49Y}$.

| Assign F1 | Time Constant | TC Error | Fit Error | Num Peaks | Function | Fit Param A | Fit Param B | Param Error A | Param Error B | (1/s) R1 | (1/s) R1 Error |
|---|---|---|---|---|---|---|---|---|---|---|---|
| 1MetH* | 627.93173 | 57.91619 | 0.99128 | 10 | A exp(-Bx) | 2.25E+04 | 0.00159 | 821.02649 | 1.46E-04 | 1.59 | 1.46E-01 |
| 2GluH | 614.62136 | 47.6759 | 1.18757 | 10 | A exp(-Bx) | 3.21E+04 | 0.00163 | 989.16693 | 1.25E-04 | 1.63 | 1.25E-01 |
| 3GluH | 649.17002 | 47.9095 | 1.34464 | 10 | A exp(-Bx) | 3.80E+04 | 0.00154 | 1103.85242 | 1.13E-04 | 1.54 | 1.13E-01 |
| 5GlnH | 543.20059 | 40.39784 | 0.99814 | 10 | A exp(-Bx) | 2.85E+04 | 0.00184 | 859.79492 | 1.36E-04 | 1.84 | 1.36E-01 |
| 6SerH | 571.56471 | 53.99666 | 0.95137 | 10 | A exp(-Bx) | 2.13E+04 | 0.00175 | 808.24615 | 1.64E-04 | 1.75 | 1.64E-01 |
| 7AspH | 606.21142 | 54.94246 | 1.15858 | 10 | A exp(-Bx) | 2.69E+04 | 0.00165 | 968.70496 | 1.48E-04 | 1.65 | 1.48E-01 |
| 9SerH | 495.81215 | 21.1151 | 0.51354 | 10 | A exp(-Bx) | 2.58E+04 | 0.00202 | 453.68045 | 8.57E-05 | 2.02 | 8.57E-02 |
| 10ValH | 568.62706 | 37.99284 | 0.96997 | 10 | A exp(-Bx) | 3.06E+04 | 0.00176 | 825.21753 | 1.17E-04 | 1.76 | 1.17E-01 |
| 11GluH | 665.91216 | 78.65261 | 1.56777 | 10 | A exp(-Bx) | 2.78E+04 | 0.0015 | 1278.37927 | 1.75E-04 | 1.5 | 1.75E-01 |
| 14LeuH | 547.01441 | 24.50971 | 0.82879 | 10 | A exp(-Bx) | 3.91E+04 | 0.00183 | 711.82178 | 8.17E-05 | 1.83 | 8.17E-02 |
| 15SerH | 606.84281 | 37.45033 | 0.58612 | 10 | A exp(-Bx) | 1.99E+04 | 0.00165 | 489.92944 | 1.01E-04 | 1.65 | 1.01E-01 |
| 18ThrH | 517.39615 | 31.10759 | 0.69985 | 10 | A exp(-Bx) | 2.48E+04 | 0.00193 | 611.06915 | 1.16E-04 | 1.93 | 1.16E-01 |
| 19PheH | 504.00033 | 24.52099 | 0.44018 | 10 | A exp(-Bx) | 1.91E+04 | 0.00198 | 382.84009 | 9.63E-05 | 1.98 | 9.63E-02 |
| 20SerH | 508.11527 | 30.5775 | 0.44559 | 10 | A exp(-Bx) | 1.58E+04 | 0.00197 | 391.02142 | 1.18E-04 | 1.97 | 1.18E-01 |
| 21AspH | 473.90233 | 33.429 | 0.57963 | 10 | A exp(-Bx) | 1.77E+04 | 0.00211 | 518.63556 | 1.48E-04 | 2.11 | 1.48E-01 |
| 22LeuH | 504.71057 | 23.23343 | 0.68732 | 10 | A exp(-Bx) | 3.19E+04 | 0.00198 | 603.5871 | 9.10E-05 | 1.98 | 9.10E-02 |
| 23TrpH | 474.83581 | 21.19411 | 0.64756 | 10 | A exp(-Bx) | 3.12E+04 | 0.00211 | 579.03314 | 9.38E-05 | 2.11 | 9.38E-02 |
| 24LysH | 457.20904 | 28.1792 | 0.85654 | 10 | A exp(-Bx) | 3.01E+04 | 0.00219 | 773.29926 | 1.34E-04 | 2.19 | 1.34E-01 |
| 25LeuH | 498.39258 | 22.0495 | 0.61665 | 10 | A exp(-Bx) | 2.98E+04 | 0.00201 | 544.04309 | 8.86E-05 | 2.01 | 8.86E-02 |
| 26LeuH | 503.36357 | 22.47058 | 0.66953 | 10 | A exp(-Bx) | 3.20E+04 | 0.00199 | 589.00354 | 8.85E-05 | 1.99 | 8.85E-02 |
| 28GluH | 515.92503 | 33.4562 | 0.88045 | 10 | A exp(-Bx) | 2.90E+04 | 0.00194 | 769.3653 | 1.25E-04 | 1.94 | 1.25E-01 |
| 29AsnH | 530.86553 | 45.15549 | 0.7113 | 10 | A exp(-Bx) | 1.76E+04 | 0.00188 | 609.83801 | 1.59E-04 | 1.88 | 1.59E-01 |
| 30AsnH | 545.5395 | 53.81132 | 0.59391 | 10 | A exp(-Bx) | 1.28E+04 | 0.00183 | 511.04407 | 1.79E-04 | 1.83 | 1.79E-01 |
| 31ValH | 545.40649 | 35.24278 | 0.98933 | 10 | A exp(-Bx) | 3.24E+04 | 0.00183 | 850.39056 | 1.18E-04 | 1.83 | 1.18E-01 |
| 32LeuH | 540.59126 | 34.8028 | 0.8623 | 10 | A exp(-Bx) | 2.84E+04 | 0.00185 | 743.84027 | 1.19E-04 | 1.85 | 1.19E-01 |
| 33SerH | 543.18998 | 32.90386 | 0.79127 | 10 | A exp(-Bx) | 2.77E+04 | 0.00184 | 681.59918 | 1.11E-04 | 1.84 | 1.11E-01 |
| 35LeuH | 652.07911 | 49.44602 | 1.02453 | 10 | A exp(-Bx) | 2.82E+04 | 0.00153 | 840.07324 | 1.16E-04 | 1.53 | 1.16E-01 |
| 37SerH | 556.82621 | 36.05858 | 0.56263 | 10 | A exp(-Bx) | 1.84E+04 | 0.0018 | 481.42294 | 1.16E-04 | 1.8 | 1.16E-01 |
| 38GlnH | 556.52519 | 46.29832 | 0.64651 | 10 | A exp(-Bx) | 1.65E+04 | 0.0018 | 552.65021 | 1.48E-04 | 1.8 | 1.48E-01 |
| 39AlaH | 565.63255 | 26.16479 | 0.54667 | 10 | A exp(-Bx) | 2.49E+04 | 0.00177 | 465.76025 | 8.16E-05 | 1.77 | 8.16E-02 |
| 40MetH | 572.72294 | 45.5397 | 1.04393 | 10 | A exp(-Bx) | 2.77E+04 | 0.00175 | 885.40912 | 1.38E-04 | 1.75 | 1.38E-01 |
| 41AspH | 523.80751 | 24.69817 | 0.67104 | 10 | A exp(-Bx) | 3.03E+04 | 0.00191 | 583.91949 | 8.98E-05 | 1.91 | 8.98E-02 |
| 42AspH | 504.96466 | 24.12772 | 0.88777 | 10 | A exp(-Bx) | 3.97E+04 | 0.00198 | 780.39203 | 9.44E-05 | 1.98 | 9.44E-02 |
| 43LeuH | 575.61899 | 25.0412 | 0.64068 | 10 | A exp(-Bx) | 3.10E+04 | 0.00174 | 543.25458 | 7.54E-05 | 1.74 | 7.54E-02 |
| 44MetH | 511.24653 | 24.74973 | 0.74033 | 10 | A exp(-Bx) | 3.26E+04 | 0.00196 | 648.48065 | 9.45E-05 | 1.96 | 9.45E-02 |
| 45LeuH | 518.85989 | 32.57167 | 0.92821 | 10 | A exp(-Bx) | 3.15E+04 | 0.00193 | 809.82751 | 1.21E-04 | 1.93 | 1.21E-01 |
| 46SerH | 495.2453 | 25.72284 | 0.7837 | 10 | A exp(-Bx) | 3.23E+04 | 0.00202 | 692.56726 | 1.05E-04 | 2.02 | 1.05E-01 |
| 48AspH | 533.74086 | 46.7978 | 1.34367 | 10 | A exp(-Bx) | 3.27E+04 | 0.00187 | 1163.17664 | 1.63E-04 | 1.87 | 1.63E-01 |
| 49TyrH | 541.06971 | 23.68384 | 0.61471 | 10 | A exp(-Bx) | 2.97E+04 | 0.00185 | 530.14001 | 8.07E-05 | 1.85 | 8.07E-02 |
| 50IleH | 568.26462 | 42.50459 | 0.935 | 10 | A exp(-Bx) | 2.64E+04 | 0.00176 | 795.59827 | 1.31E-04 | 1.76 | 1.31E-01 |
| 51GluH | 581.11564 | 58.5938 | 1.15461 | 10 | A exp(-Bx) | 2.43E+04 | 0.00172 | 976.51019 | 1.72E-04 | 1.72 | 1.72E-01 |
| 52GlnH | 555.9332 | 28.03075 | 0.86485 | 10 | A exp(-Bx) | 3.62E+04 | 0.0018 | 740.34576 | 9.05E-05 | 1.8 | 9.05E-02 |
| 53TrpH | 485.56705 | 25.70476 | 0.89892 | 10 | A exp(-Bx) | 3.61E+04 | 0.00206 | 789.9718 | 1.09E-04 | 2.06 | 1.09E-01 |
| 54PheH | 495.78411 | 28.58508 | 1.09915 | 10 | A exp(-Bx) | 4.09E+04 | 0.00202 | 971.14874 | 1.16E-04 | 2.02 | 1.16E-01 |
| 55ThrH | 488.60776 | 28.55337 | 0.63659 | 10 | A exp(-Bx) | 2.34E+04 | 0.00205 | 564.7417 | 1.19E-04 | 2.05 | 1.19E-01 |
| 56GluH | 588.39779 | 33.7703 | 0.68387 | 10 | A exp(-Bx) | 2.50E+04 | 0.0017 | 576.42706 | 9.72E-05 | 1.7 | 9.72E-02 |
| 57AspH | 568.91532 | 34.16347 | 0.85626 | 10 | A exp(-Bx) | 3.00E+04 | 0.00176 | 727.5578 | 1.05E-04 | 1.76 | 1.05E-01 |
| 59GlyH | 566.44497 | 30.17734 | 0.80072 | 10 | A exp(-Bx) | 3.17E+04 | 0.00177 | 681.86884 | 9.38E-05 | 1.77 | 9.38E-02 |
| 61AspH | 633.83377 | 65.52355 | 1.10049 | 10 | A exp(-Bx) | 2.24E+04 | 0.00158 | 909.20172 | 1.61E-04 | 1.58 | 1.61E-01 |
| 62GluH | 581.01859 | 28.52778 | 1.04817 | 10 | A exp(-Bx) | 4.49E+04 | 0.00172 | 886.4303 | 8.43E-05 | 1.72 | 8.43E-02 |
| 63AlaH | 622.23408 | 68.43105 | 1.41353 | 10 | A exp(-Bx) | 2.71E+04 | 0.00161 | 1173.62488 | 1.75E-04 | 1.61 | 1.75E-01 |
| 65ArgH | 545.22578 | 27.54325 | 1.11749 | 10 | A exp(-Bx) | 4.68E+04 | 0.00183 | 961.71271 | 9.24E-05 | 1.83 | 9.24E-02 |
| 66MetH | 609.86037 | 66.89551 | 1.03241 | 10 | A exp(-Bx) | 1.99E+04 | 0.00164 | 861.82965 | 1.78E-04 | 1.64 | 1.78E-01 |
| 68GluH | 577.3521 | 30.56491 | 0.8555 | 10 | A exp(-Bx) | 3.40E+04 | 0.00173 | 724.82056 | 9.14E-05 | 1.73 | 9.14E-02 |
| 69AlaH | 576.05125 | 48.25707 | 1.1419 | 10 | A exp(-Bx) | 2.88E+04 | 0.00174 | 968.052 | 1.44E-04 | 1.74 | 1.44E-01 |
| 70AlaH | 724.20964 | 67.2149 | 1.45805 | 10 | A exp(-Bx) | 3.25E+04 | 0.00138 | 1163.18005 | 1.27E-04 | 1.38 | 1.27E-01 |
| 72ArgH | 647.92404 | 58.80923 | 1.79537 | 10 | A exp(-Bx) | 4.14E+04 | 0.00154 | 1474.61841 | 1.39E-04 | 1.54 | 1.39E-01 |
| 73ValH | 855.36526 | 131.72907 | 3.83388 | 10 | A exp(-Bx) | 5.19E+04 | 0.00117 | 2934.10181 | 1.76E-04 | 1.17 | 1.76E-01 |

#### Table S10. $T_1$ relaxation for p53-TAD$^{W53G}$.

| Assign F1 | Assign F2 | Time Constant | TC Error | Fit Error | Num Peaks | Function | Fit Param A | Fit Param B | Param Error A | Param Error B | (1/s) R1 | (1/s) R1 Error |
|---|---|---|---|---|---|---|---|---|---|---|---|---|
| 1MetH* | 1MetN | 617.21977 | 48.46708 | 3.60561 | 10 | A exp(-Bx) | 29658.9 | 0.00162 | 924.43207 | 0.000126449 | 1.62 | 0.126449 |
| 2GluH | 2GluN | 674.16097 | 28.41354 | 2.31976 | 10 | A exp(-Bx) | 35101.8 | 0.00148 | 580.95203 | 6.24063E-05 | 1.48 | 0.0624063 |
| 3GluH | 3GluN | 676.09268 | 15.661 | 1.77622 | 10 | A exp(-Bx) | 48297.6 | 0.00148 | 439.64926 | 3.42432E-05 | 1.48 | 0.0342432 |
| 5GlnH | 5GlnN | 647.12563 | 33.58933 | 3.12269 | 10 | A exp(-Bx) | 38547 | 0.00155 | 790.55579 | 7.99942E-05 | 1.55 | 0.0799942 |
| 6SerH | 6SerN | 565.10229 | 37.72297 | 2.76716 | 10 | A exp(-Bx) | 26984 | 0.00177 | 726.62878 | 0.000117606 | 1.77 | 0.117606 |
| 7AspH | 7AspN | 561.20086 | 19.69532 | 2.31078 | 10 | A exp(-Bx) | 42752.8 | 0.00178 | 607.86951 | 6.24586E-05 | 1.78 | 0.0624586 |
| 9SerH | 9SerN | 499.47305 | 30.05643 | 3.26224 | 10 | A exp(-Bx) | 35776.4 | 0.002 | 886.29889 | 0.000120046 | 2 | 0.120046 |
| 10ValH | 10ValN | 537.83939 | 11.2037 | 1.49357 | 10 | A exp(-Bx) | 46750 | 0.00186 | 397.54517 | 0.000038714 | 1.86 | 0.038714 |
| 11GluH | 11GluN | 644.63179 | 13.31635 | 1.41743 | 10 | A exp(-Bx) | 43876.9 | 0.00155 | 359.17374 | 3.20315E-05 | 1.55 | 0.0320315 |
| 14LeuH | 14LeuN | 644.66377 | 18.3663 | 2.25714 | 10 | A exp(-Bx) | 50685.6 | 0.00155 | 572.00293 | 4.41574E-05 | 1.55 | 0.0441574 |
| 15SerH | 15SerN | 541.42842 | 38.15728 | 3.37528 | 10 | A exp(-Bx) | 31344.5 | 0.00185 | 896.77209 | 0.000129525 | 1.85 | 0.129525 |
| 16GlnH | 16GlnN | 700.79816 | 35.53004 | 4.83154 | 10 | A exp(-Bx) | 60625.5 | 0.00143 | 1197.78601 | 7.21602E-05 | 1.43 | 0.0721602 |
| 17GluH | 17GluN | 536.86769 | 25.83493 | 5.16274 | 10 | A exp(-Bx) | 70099.1 | 0.00186 | 1374.86108 | 8.94274E-05 | 1.86 | 0.0894274 |
| 18ThrH | 18ThrN | 490.71281 | 20.8609 | 2.24178 | 10 | A exp(-Bx) | 34811 | 0.00204 | 611.9967 | 0.000086476 | 2.04 | 0.086476 |
| 19PheH | 19PheN | 493.30199 | 27.13383 | 2.31625 | 10 | A exp(-Bx) | 27816.8 | 0.00203 | 631.47198 | 0.000111167 | 2.03 | 0.111167 |
| 20SerH | 20SerN | 486.31407 | 27.8582 | 2.04128 | 10 | A exp(-Bx) | 23587 | 0.00206 | 558.72943 | 0.000117409 | 2.06 | 0.117409 |
| 21AspH | 21AspN | 441.37379 | 23.91221 | 2.04373 | 10 | A exp(-Bx) | 25278.8 | 0.00227 | 574.87268 | 0.000122388 | 2.27 | 0.122388 |
| 22LeuH | 22LeuN | 505.53043 | 9.83401 | 1.4516 | 10 | A exp(-Bx) | 48959.1 | 0.00198 | 392.61115 | 3.84656E-05 | 1.98 | 0.0384656 |
| 23TrpH | 23TrpN | 495.20961 | 7.2852 | 1.02011 | 10 | A exp(-Bx) | 45650.1 | 0.00202 | 277.77783 | 2.97009E-05 | 2.02 | 0.0297009 |
| 24LysH | 24LysN | 489.22264 | 9.99496 | 1.38797 | 10 | A exp(-Bx) | 44806.9 | 0.00204 | 379.27136 | 4.17433E-05 | 2.04 | 0.0417433 |
| 25LeuH | 25LeuN | 535.3284 | 20.90148 | 2.3235 | 10 | A exp(-Bx) | 38822 | 0.00187 | 618.55731 | 7.28242E-05 | 1.87 | 0.0728242 |
| 26LeuH | 26LeuN | 592.13706 | 16.08585 | 2.16726 | 10 | A exp(-Bx) | 51477 | 0.00169 | 561.90552 | 4.58437E-05 | 1.69 | 0.0458437 |
| 28GluH | 28GluN | 523.74661 | 11.73271 | 1.65472 | 10 | A exp(-Bx) | 48317.8 | 0.00191 | 443.68213 | 4.27502E-05 | 1.91 | 0.0427502 |
| 29AsnH | 29AsnN | 520.10217 | 33.47926 | 2.29319 | 10 | A exp(-Bx) | 23407 | 0.00192 | 616.06421 | 0.000123257 | 1.92 | 0.123257 |
| 30AsnH | 30AsnN | 492.07916 | 29.67995 | 1.89757 | 10 | A exp(-Bx) | 20800.9 | 0.00203 | 517.68091 | 0.00012213 | 2.03 | 0.12213 |
| 31ValH | 31ValN | 563.03372 | 19.80823 | 2.28683 | 10 | A exp(-Bx) | 42195.4 | 0.00178 | 601.10022 | 0.000062408 | 1.78 | 0.062408 |
| 32LeuH | 32LeuN | 587.95011 | 17.45719 | 1.87458 | 10 | A exp(-Bx) | 40770.1 | 0.0017 | 486.90833 | 5.04558E-05 | 1.7 | 0.0504558 |
| 33SerH | 33SerN | 579.11367 | 30.7109 | 2.64141 | 10 | A exp(-Bx) | 32283.2 | 0.00173 | 688.99426 | 9.13164E-05 | 1.73 | 0.0913164 |
| 35LeuH | 35LeuN | 639.69267 | 14.70706 | 1.7152 | 10 | A exp(-Bx) | 47750.6 | 0.00156 | 435.56595 | 3.59215E-05 | 1.56 | 0.0359215 |
| 37SerH | 37SerN | 516.01882 | 31.69426 | 2.61239 | 10 | A exp(-Bx) | 27962 | 0.00194 | 703.35034 | 0.000118582 | 1.94 | 0.118582 |
| 38GlnH | 38GlnN | 509.46561 | 39.63071 | 2.57896 | 10 | A exp(-Bx) | 21878.7 | 0.00196 | 696.81671 | 0.000151774 | 1.96 | 0.151774 |
| 39AlaH | 39AlaN | 585.72818 | 35.3594 | 4.11592 | 10 | A exp(-Bx) | 44172.2 | 0.00171 | 1070.30139 | 0.000102692 | 1.71 | 0.102692 |
| 40MetH | 40MetN | 575.52269 | 32.06898 | 3.21767 | 10 | A exp(-Bx) | 37459.3 | 0.00174 | 840.63312 | 9.65202E-05 | 1.74 | 0.0965202 |
| 41AspH | 41AspN | 625.9393 | 24.33737 | 2.68822 | 10 | A exp(-Bx) | 44386.1 | 0.0016 | 686.56024 | 6.20232E-05 | 1.6 | 0.0620232 |
| 42AspH | 42AspN | 611.89685 | 21.03872 | 2.63753 | 10 | A exp(-Bx) | 49349.6 | 0.00163 | 677.80688 | 5.61242E-05 | 1.63 | 0.0561242 |
| 43LeuH | 43LeuN | 603.75441 | 9.32218 | 1.35937 | 10 | A exp(-Bx) | 56599.8 | 0.00166 | 350.20886 | 2.55678E-05 | 1.66 | 0.0255678 |
| 44MetH | 44MetN | 622.82455 | 15.37326 | 2.21219 | 10 | A exp(-Bx) | 57513.7 | 0.00161 | 565.73901 | 3.96068E-05 | 1.61 | 0.0396068 |
| 45LeuH | 45LeuN | 640.07963 | 13.49213 | 1.76727 | 10 | A exp(-Bx) | 53595.6 | 0.00156 | 448.21643 | 0.000032917 | 1.56 | 0.032917 |
| 46SerH | 46SerN | 589.05399 | 20.25366 | 2.36757 | 10 | A exp(-Bx) | 44474.8 | 0.0017 | 614.71082 | 5.83016E-05 | 1.7 | 0.0583016 |
| 48AspH | 48AspN | 607.15614 | 20.67675 | 2.73713 | 10 | A exp(-Bx) | 51746.3 | 0.00165 | 704.87604 | 5.60246E-05 | 1.65 | 0.0560246 |
| 49AspH | 49AspN | 556.34608 | 20.38625 | 5.7754 | 10 | A exp(-Bx) | 102461 | 0.0018 | 1523.03687 | 6.57757E-05 | 1.8 | 0.0657757 |
| 50IleH | 50IleN | 649.85159 | 17.21241 | 3.17855 | 10 | A exp(-Bx) | 76707.5 | 0.00154 | 803.79028 | 4.07295E-05 | 1.54 | 0.0407295 |
| 51GluH | 51GluN | 587.24505 | 11.45632 | 1.85852 | 10 | A exp(-Bx) | 61496.8 | 0.0017 | 482.89441 | 3.32079E-05 | 1.7 | 0.0332079 |
| 52GlnH | 52GlnN | 603.60787 | 22.64001 | 2.69952 | 10 | A exp(-Bx) | 46376.5 | 0.00166 | 696.28302 | 6.20522E-05 | 1.66 | 0.0620522 |
| 53GlyH | 53GlyN | 579.80441 | 28.90321 | 3.04872 | 10 | A exp(-Bx) | 39620.4 | 0.00172 | 794.97083 | 8.57646E-05 | 1.72 | 0.0857646 |
| 55ThrH | 55ThrN | 566.84829 | 37.92342 | 3.46878 | 10 | A exp(-Bx) | 33740 | 0.00176 | 910.09912 | 0.000117501 | 1.76 | 0.117501 |
| 56GluH | 56GluN | 631.89584 | 19.60504 | 2.11483 | 10 | A exp(-Bx) | 43699.7 | 0.00158 | 538.80994 | 4.90523E-05 | 1.58 | 0.0490523 |
| 57AspH | 57AspN | 581.85996 | 25.76966 | 3.38382 | 10 | A exp(-Bx) | 49447.8 | 0.00172 | 881.4137 | 7.59665E-05 | 1.72 | 0.0759665 |
| 59GlyH | 59GlyN | 617.88209 | 12.10845 | 1.63209 | 10 | A exp(-Bx) | 53481.8 | 0.00162 | 418.3222 | 3.17037E-05 | 1.62 | 0.0317037 |
| 61AspH | 61AspN | 611.24103 | 31.79847 | 3.8247 | 10 | A exp(-Bx) | 47373.6 | 0.00164 | 983.1731 | 0.000084881 | 1.64 | 0.084881 |
| 62GluH | 62GluN | 671.31846 | 18.72849 | 1.75302 | 10 | A exp(-Bx) | 40045 | 0.00149 | 439.46487 | 4.15248E-05 | 1.49 | 0.0415248 |
| 63AlaH | 63AlaN | 611.40835 | 23.10273 | 2.97606 | 10 | A exp(-Bx) | 50679.2 | 0.00164 | 764.8819 | 6.17137E-05 | 1.64 | 0.0617137 |
| 65ArgH | 65ArgN | 600.63357 | 38.50251 | 3.4873 | 10 | A exp(-Bx) | 35167.1 | 0.00166 | 900.68573 | 0.000106291 | 1.66 | 0.106291 |
| 66MetH | 66MetN | 576.03294 | 24.31359 | 2.00514 | 10 | A exp(-Bx) | 30776.6 | 0.00174 | 523.78674 | 7.31447E-05 | 1.74 | 0.0731447 |
| 68GluH | 68GluN | 603.68109 | 31.11443 | 3.76554 | 10 | A exp(-Bx) | 47134.7 | 0.00166 | 971.21021 | 8.51525E-05 | 1.66 | 0.0851525 |
| 69AlaH | 69AlaN | 598.5452 | 36.70602 | 4.01064 | 10 | A exp(-Bx) | 42277.6 | 0.00167 | 1036.82654 | 0.000102075 | 1.67 | 0.102075 |
| 70AlaH | 70AlaN | 742.65962 | 31.85268 | 3.10504 | 10 | A exp(-Bx) | 45820.2 | 0.00135 | 758.26733 | 5.76461E-05 | 1.35 | 0.0576461 |
| 72ArgH | 72ArgN | 743.96997 | 54.12589 | 5.42956 | 10 | A exp(-Bx) | 47396 | 0.00134 | 1325.47571 | 9.72778E-05 | 1.34 | 0.0972778 |
| 73ValH | 73ValN | 1236.76589 | 27.582 | 2.62959 | 10 | A exp(-Bx) | 75489.9 | 0.00080856 | 571.87378 | 1.80233E-05 | 0.80856 | 0.0180233 |

### Table S11. $T_2$ relaxation for p53-TAD$^{WT}$.

| Assign F2 | Time Constant | TC error | Fit error | Number Peaks | Function | Fit Parameter A | Fit parameter B | Error Paramater A | Error Parameter B | (1/s) R2 | (1/s) R2 Error |
|---|---|---|---|---|---|---|---|---|---|---|---|
| 1MetN | 196.26235 | 12.6342 | 0.6281 | 10 | A exp(-Bx) | 39772.2 | 0.0051 | 1271.75452 | 0.000326652 | 5.1 | 0.326652 |
| 2GluN | 200.54205 | 7.53118 | 0.51354 | 10 | A exp(-Bx) | 55887.1 | 0.00499 | 1034.44141 | 0.000187 | 4.99 | 0.187 |
| 3GluN | 215.42221 | 8.35836 | 0.80402 | 10 | A exp(-Bx) | 86463.4 | 0.00464 | 1593.93494 | 0.000179841 | 4.64 | 0.179841 |
| 5GlnN | 192.71679 | 5.71117 | 0.39685 | 10 | A exp(-Bx) | 54084.4 | 0.00519 | 805.67572 | 0.00015364 | 5.19 | 0.15364 |
| 6SerN | 161.7585 | 5.2277 | 0.34992 | 10 | A exp(-Bx) | 42586.2 | 0.00618 | 744.4599 | 0.000199583 | 6.18 | 0.199583 |
| 7AspN | 175.79719 | 4.18892 | 0.3188 | 10 | A exp(-Bx) | 53143 | 0.00569 | 663.24286 | 0.000135467 | 5.69 | 0.135467 |
| 9SerN | 170.2932 | 4.75459 | 0.36499 | 10 | A exp(-Bx) | 51698.2 | 0.00587 | 765.72626 | 0.000163825 | 5.87 | 0.163825 |
| 10ValN | 191.06927 | 11.27752 | 0.95275 | 10 | A exp(-Bx) | 65342.3 | 0.00523 | 1941.42249 | 0.000307841 | 5.23 | 0.307841 |
| 11GluN | 237.09586 | 13.76894 | 0.77389 | 10 | A exp(-Bx) | 57591.2 | 0.00422 | 1504.08423 | 0.000244115 | 4.22 | 0.244115 |
| 14LeuN | 194.15883 | 6.1496 | 0.52467 | 10 | A exp(-Bx) | 67138.2 | 0.00515 | 1065.02441 | 0.000162966 | 5.15 | 0.162966 |
| 15SerN | 169.19081 | 5.16942 | 0.33177 | 10 | A exp(-Bx) | 42905.3 | 0.00591 | 697.12573 | 0.00018042 | 5.91 | 0.18042 |
| 16GlnN | 184.77315 | 4.7202 | 0.53431 | 10 | A exp(-Bx) | 83831.4 | 0.00541 | 1097.7074 | 0.000138166 | 5.41 | 0.138166 |
| 17GluN | 170.60256 | 4.55037 | 0.6391 | 10 | A exp(-Bx) | 94758.5 | 0.00586 | 1339.91382 | 0.000156231 | 5.86 | 0.156231 |
| 18ThrN | 165.49307 | 4.95298 | 0.39269 | 10 | A exp(-Bx) | 51715.9 | 0.00604 | 830.21124 | 0.000180684 | 6.04 | 0.180684 |
| 19PheN | 150.71945 | 2.10094 | 0.14629 | 10 | A exp(-Bx) | 41070.3 | 0.00663 | 317.61966 | 9.24679E-05 | 6.63 | 0.0924679 |
| 20SerN | 148.31342 | 3.39276 | 0.1898 | 10 | A exp(-Bx) | 32465.5 | 0.00674 | 414.02292 | 0.000154158 | 6.74 | 0.154158 |
| 21AspN | 145.10121 | 2.90164 | 0.18233 | 10 | A exp(-Bx) | 35668.6 | 0.00689 | 400.38416 | 0.000137762 | 6.89 | 0.137762 |
| 22LeuN | 157.66985 | 4.61822 | 0.55448 | 10 | A exp(-Bx) | 74292.2 | 0.00634 | 1188.18323 | 0.000185612 | 6.34 | 0.185612 |
| 23TrpN | 159.62282 | 2.6912 | 0.2524 | 10 | A exp(-Bx) | 58761.9 | 0.00626 | 538.89673 | 0.000105592 | 6.26 | 0.105592 |
| 24LysN | 157.23106 | 3.88016 | 0.40637 | 10 | A exp(-Bx) | 64597.2 | 0.00636 | 871.51038 | 0.000156859 | 6.36 | 0.156859 |
| 25LeuN | 183.89139 | 9.60158 | 0.63541 | 10 | A exp(-Bx) | 48822.6 | 0.00544 | 1306.73804 | 0.000283166 | 5.44 | 0.283166 |
| 26LeuN | 200.07167 | 8.89268 | 0.74627 | 10 | A exp(-Bx) | 68626.2 | 0.005 | 1504.28601 | 0.000221721 | 5 | 0.221721 |
| 28GluN | 164.57252 | 5.46285 | 0.53896 | 10 | A exp(-Bx) | 63961.2 | 0.00608 | 1141.00537 | 0.000201478 | 6.08 | 0.201478 |
| 29AsnN | 148.16128 | 3.21172 | 0.1968 | 10 | A exp(-Bx) | 35002 | 0.00675 | 423.14502 | 0.000146239 | 6.75 | 0.146239 |
| 30AsnN | 157.18501 | 5.29863 | 0.24052 | 10 | A exp(-Bx) | 27999.5 | 0.00636 | 515.78326 | 0.000214215 | 6.36 | 0.214215 |
| 31ValN | 191.40029 | 4.98873 | 0.3927 | 10 | A exp(-Bx) | 60842.6 | 0.00522 | 799.86768 | 0.000136085 | 5.22 | 0.136085 |
| 32LeuN | 213.19981 | 7.52299 | 0.48472 | 10 | A exp(-Bx) | 57127.1 | 0.00469 | 963.25397 | 0.000165302 | 4.69 | 0.165302 |
| 33SerN | 174.85157 | 5.27706 | 0.31795 | 10 | A exp(-Bx) | 41819.1 | 0.00572 | 662.3067 | 0.000172448 | 5.72 | 0.172448 |
| 35LeuN | 220.52305 | 7.86983 | 0.59299 | 10 | A exp(-Bx) | 69732.8 | 0.00453 | 1167.91125 | 0.000161624 | 4.53 | 0.161624 |
| 37SerN | 181.87913 | 4.2466 | 0.20942 | 10 | A exp(-Bx) | 35274.9 | 0.0055 | 425.16446 | 0.000128304 | 5.5 | 0.128304 |
| 38GlnN | 154.81091 | 3.74414 | 0.21814 | 10 | A exp(-Bx) | 35349.7 | 0.00646 | 469.92194 | 0.000156133 | 6.46 | 0.156133 |
| 39AlaN | 189.74602 | 7.3127 | 0.49438 | 10 | A exp(-Bx) | 51743.7 | 0.00527 | 1009.07855 | 0.00020281 | 5.27 | 0.20281 |
| 40MetN | 179.34766 | 3.75168 | 0.26456 | 10 | A exp(-Bx) | 50389.2 | 0.00558 | 547.50024 | 0.000116585 | 5.58 | 0.116585 |
| 41AspN | 183.29736 | 4.44342 | 0.36949 | 10 | A exp(-Bx) | 60990.1 | 0.00546 | 760.59283 | 0.000132175 | 5.46 | 0.132175 |
| 42AspN | 204.23292 | 3.74461 | 0.33259 | 10 | A exp(-Bx) | 74431.1 | 0.0049 | 667.26898 | 8.97447E-05 | 4.9 | 0.0897447 |
| 43LeuN | 197.33806 | 7.48654 | 0.58297 | 10 | A exp(-Bx) | 62555.5 | 0.00507 | 1178.88208 | 0.000191971 | 5.07 | 0.191971 |
| 44MetN | 194.27471 | 4.88274 | 0.48896 | 10 | A exp(-Bx) | 78703.7 | 0.00515 | 990.78845 | 0.000129288 | 5.15 | 0.129288 |
| 45LeuN | 190.90076 | 6.55824 | 0.52332 | 10 | A exp(-Bx) | 60530 | 0.00524 | 1049.60229 | 0.000179746 | 5.24 | 0.179746 |
| 46SerN | 172.75358 | 5.46258 | 0.52032 | 10 | A exp(-Bx) | 64210.6 | 0.00579 | 1070.66675 | 0.000182856 | 5.79 | 0.182856 |
| 48AspN | 174.31585 | 3.66417 | 0.32104 | 10 | A exp(-Bx) | 59641.6 | 0.00574 | 659.0271 | 0.000120534 | 5.74 | 0.120534 |
| 49AspN | 206.60606 | 5.00895 | 0.36423 | 10 | A exp(-Bx) | 61756.5 | 0.00484 | 727.62982 | 0.000117275 | 4.84 | 0.117275 |
| 50IleN | 218.15873 | 3.23665 | 0.25629 | 10 | A exp(-Bx) | 71084.7 | 0.00458 | 498.35733 | 6.79916E-05 | 4.58 | 0.0679916 |
| 51GluN | 199.35414 | 6.08642 | 0.52086 | 10 | A exp(-Bx) | 69479.2 | 0.00502 | 1049.05957 | 0.000153005 | 5.02 | 0.153005 |
| 52GlnN | 177.58248 | 4.51084 | 0.40078 | 10 | A exp(-Bx) | 62770.3 | 0.00563 | 831.513 | 0.000142948 | 5.63 | 0.142948 |
| 53TrpN | 175.26185 | 6.85036 | 1.08987 | 10 | A exp(-Bx) | 110635 | 0.00571 | 2265.63257 | 0.000222678 | 5.71 | 0.222678 |
| 55ThrN | 181.91199 | 9.36436 | 0.64424 | 10 | A exp(-Bx) | 50109.1 | 0.0055 | 1328.68799 | 0.000282234 | 5.5 | 0.282234 |
| 56GluN | 195.4492 | 10.44721 | 0.81806 | 10 | A exp(-Bx) | 62241.8 | 0.00512 | 1657.97607 | 0.000272707 | 5.12 | 0.272707 |
| 57AspN | 195.8 | 6.7182 | 0.62624 | 10 | A exp(-Bx) | 74136.2 | 0.00511 | 1268.68372 | 0.000175032 | 5.11 | 0.175032 |
| 59GlyN | 203.85023 | 11.43823 | 0.90791 | 10 | A exp(-Bx) | 66543.9 | 0.00491 | 1822.28113 | 0.000274395 | 4.91 | 0.274395 |
| 61AspN | 176.58928 | 4.74423 | 0.39166 | 10 | A exp(-Bx) | 57949.3 | 0.00566 | 813.77051 | 0.000152028 | 5.66 | 0.152028 |
| 62GluN | 196.92984 | 6.43884 | 0.46656 | 10 | A exp(-Bx) | 58038.2 | 0.00508 | 943.92822 | 0.000165852 | 5.08 | 0.165852 |
| 63AlaN | 224.96883 | 8.84174 | 0.57268 | 10 | A exp(-Bx) | 61579 | 0.00445 | 1123.18628 | 0.000174431 | 4.45 | 0.174431 |
| 65ArgN | 170.60256 | 4.55037 | 0.6391 | 10 | A exp(-Bx) | 94758.5 | 0.00586 | 1339.91382 | 0.000156231 | 5.86 | 0.156231 |
| 66MetN | 182.81843 | 3.93075 | 0.23257 | 10 | A exp(-Bx) | 43249.6 | 0.00547 | 478.98846 | 0.000117553 | 5.47 | 0.117553 |
| 68GluN | 196.53539 | 7.00997 | 0.49571 | 10 | A exp(-Bx) | 56509.9 | 0.00509 | 1003.36853 | 0.000181252 | 5.09 | 0.181252 |
| 69AlaN | 222.60828 | 8.22696 | 0.38461 | 10 | A exp(-Bx) | 43889.3 | 0.00449 | 757.27563 | 0.000165793 | 4.49 | 0.165793 |
| 70AlaN | 264.3274 | 12.87312 | 0.66746 | 10 | A exp(-Bx) | 60800 | 0.00378 | 1248.4613 | 0.000183812 | 3.78 | 0.183812 |
| 72ArgN | 184.77315 | 4.7202 | 0.53431 | 10 | A exp(-Bx) | 83831.4 | 0.00541 | 1097.7074 | 0.000138166 | 5.41 | 0.138166 |
| 73ValN | 363.83678 | 17.48287 | 0.87643 | 10 | A exp(-Bx) | 97602.8 | 0.00275 | 1587.44995 | 0.000131765 | 2.75 | 0.131765 |

#### Table S12. $T_2$ relaxation for p53-TAD$^{K24N}$.

| Assign F1 | Time Constant | TC Error | Fit Error | Num Peaks | Function | Fit Param A | Fit Param B | Param Error A | Param Error B | (1/s) R2 | (1/s) R2 Error |
|---|---|---|---|---|---|---|---|---|---|---|---|
| 1MetH* | 239.34926 | 24.16485 | 4.20401 | 10 | A exp(-Bx) | 27561 | 0.00418 | 1236.2373 | 0.000417599 | 4.18 | 0.417599 |
| 2GluH | 234.88884 | 23.47937 | 5.62286 | 10 | A exp(-Bx) | 36960.7 | 0.00426 | 1659.66736 | 0.000421392 | 4.26 | 0.421392 |
| 3GluH | 261.08903 | 21.09561 | 5.67767 | 10 | A exp(-Bx) | 48037.8 | 0.00383 | 1642.38257 | 0.000307473 | 3.83 | 0.307473 |
| 5GlnH | 226.88679 | 25.56757 | 5.87029 | 10 | A exp(-Bx) | 33890 | 0.00441 | 1744.89429 | 0.000490522 | 4.41 | 0.490522 |
| 6SerH | 215.09355 | 23.86761 | 4.60107 | 10 | A exp(-Bx) | 26488.2 | 0.00465 | 1382.90662 | 0.000509686 | 4.65 | 0.509686 |
| 7AspH | 231.01272 | 8.20179 | 1.57389 | 10 | A exp(-Bx) | 28702.1 | 0.00433 | 466.10577 | 0.000153494 | 4.33 | 0.153494 |
| 9SerH | 230.58538 | 23.56518 | 5.03915 | 10 | A exp(-Bx) | 32193.4 | 0.00434 | 1492.92371 | 0.000438673 | 4.34 | 0.438673 |
| 10ValH | 197.01397 | 12.95245 | 7.59773 | 10 | A exp(-Bx) | 71359.5 | 0.00508 | 2325.65137 | 0.000332271 | 5.08 | 0.332271 |
| 11GluH | 259.12089 | 14.64657 | 2.8358 | 10 | A exp(-Bx) | 34072.4 | 0.00386 | 821.45953 | 0.000217445 | 3.86 | 0.217445 |
| 14LeuH | 248.91392 | 20.22572 | 4.36794 | 10 | A exp(-Bx) | 35948.9 | 0.00402 | 1272.50647 | 0.000324314 | 4.02 | 0.324314 |
| 15SerH | 238.23159 | 21.91268 | 3.94039 | 10 | A exp(-Bx) | 28256.7 | 0.0042 | 1159.80408 | 0.000382885 | 4.2 | 0.382885 |
| 16GlnH | 245.68744 | 16.01726 | 5.48122 | 10 | A exp(-Bx) | 55899.2 | 0.00407 | 1603.68433 | 0.000264234 | 4.07 | 0.264234 |
| 17GluH | 208.79873 | 11.89072 | 3.10759 | 10 | A exp(-Bx) | 34190.8 | 0.00479 | 938.6955 | 0.000271864 | 4.79 | 0.271864 |
| 18ThrH | 228.66993 | 19.7922 | 4.60853 | 10 | A exp(-Bx) | 34559.9 | 0.00437 | 1367.64697 | 0.000375715 | 4.37 | 0.375715 |
| 19PheH | 236.27128 | 24.14933 | 4.87493 | 10 | A exp(-Bx) | 31416.1 | 0.00423 | 1436.97302 | 0.000428169 | 4.23 | 0.428169 |
| 20SerH | 188.23377 | 10.31277 | 2.67863 | 10 | A exp(-Bx) | 29883 | 0.00531 | 830.24408 | 0.00029019 | 5.31 | 0.29019 |
| 21AspH | 216.25937 | 13.79829 | 3.20508 | 10 | A exp(-Bx) | 31362.1 | 0.00462 | 946.3772 | 0.000293845 | 4.62 | 0.293845 |
| 22LeuH | 216.8402 | 11.85792 | 3.89804 | 10 | A exp(-Bx) | 45245.8 | 0.00461 | 1169.76538 | 0.000251441 | 4.61 | 0.251441 |
| 23TrpH | 198.5362 | 8.90682 | 2.72248 | 10 | A exp(-Bx) | 37514.3 | 0.00504 | 833.09021 | 0.000225513 | 5.04 | 0.225513 |
| 24AsnH | 175.01073 | 7.79041 | 2.15639 | 10 | A exp(-Bx) | 29138.2 | 0.00571 | 679.6637 | 0.000253848 | 5.71 | 0.253848 |
| 26LeuH | 262.51494 | 23.20561 | 5.2212 | 10 | A exp(-Bx) | 40453 | 0.00381 | 1506.11951 | 0.000334142 | 3.81 | 0.334142 |
| 28GluH | 184.47093 | 13.12346 | 4.86261 | 10 | A exp(-Bx) | 41689.5 | 0.00542 | 1514.69055 | 0.000383717 | 5.42 | 0.383717 |
| 29AsnH | 207.92858 | 14.54133 | 2.72129 | 10 | A exp(-Bx) | 24428.1 | 0.00481 | 824.01849 | 0.000334709 | 4.81 | 0.334709 |
| 30AsnH | 213.67384 | 21.53734 | 3.96332 | 10 | A exp(-Bx) | 25018.9 | 0.00468 | 1193.13794 | 0.000467028 | 4.68 | 0.467028 |
| 31ValH | 208.29467 | 19.63539 | 6.58713 | 10 | A exp(-Bx) | 44069.1 | 0.0048 | 1994.16797 | 0.000448616 | 4.8 | 0.448616 |
| 32LeuH | 258.40716 | 18.30865 | 3.1324 | 10 | A exp(-Bx) | 30037.9 | 0.00387 | 907.672 | 0.000272825 | 3.87 | 0.272825 |
| 33SerH | 218.22639 | 16.33064 | 3.22142 | 10 | A exp(-Bx) | 27451.1 | 0.00458 | 965.40186 | 0.000341018 | 4.58 | 0.341018 |
| 35LeuH | 245.60885 | 12.0449 | 3.21817 | 10 | A exp(-Bx) | 43544.7 | 0.00407 | 941.62671 | 0.000199193 | 4.07 | 0.199193 |
| 37SerH | 218.00824 | 10.50399 | 1.90364 | 10 | A exp(-Bx) | 25062.3 | 0.00459 | 569.63281 | 0.000220498 | 4.59 | 0.220498 |
| 38GlnH | 192.53935 | 14.1564 | 3.12694 | 10 | A exp(-Bx) | 26187.2 | 0.00519 | 963.93091 | 0.000379826 | 5.19 | 0.379826 |
| 39AlaH | 263.53263 | 9.11151 | 1.78832 | 10 | A exp(-Bx) | 35318.6 | 0.00379 | 516.43524 | 0.00013104 | 3.79 | 0.13104 |
| 40MetH | 204.15043 | 15.19747 | 4.07046 | 10 | A exp(-Bx) | 34171.2 | 0.0049 | 1237.64185 | 0.000362647 | 4.9 | 0.362647 |
| 41AspH | 214.47536 | 13.54596 | 3.63691 | 10 | A exp(-Bx) | 36461.1 | 0.00466 | 1093.98523 | 0.000293314 | 4.66 | 0.293314 |
| 42AspH | 203.41399 | 16.39869 | 5.87551 | 10 | A exp(-Bx) | 45551.2 | 0.00492 | 1788.20874 | 0.000393778 | 4.92 | 0.393778 |
| 43LeuH | 239.29015 | 14.5903 | 3.63971 | 10 | A exp(-Bx) | 39188.1 | 0.00418 | 1068.47778 | 0.000253868 | 4.18 | 0.253868 |
| 44MetH | 242.63929 | 23.05489 | 5.67596 | 10 | A exp(-Bx) | 39634.7 | 0.00412 | 1661.71277 | 0.000388125 | 4.12 | 0.388125 |
| 45LeuH | 231.99932 | 15.66374 | 3.92653 | 10 | A exp(-Bx) | 37770.6 | 0.00431 | 1159.82715 | 0.000289705 | 4.31 | 0.289705 |
| 46SerH | 204.95475 | 10.60381 | 3.05537 | 10 | A exp(-Bx) | 36847.3 | 0.00488 | 928.31158 | 0.000251761 | 4.88 | 0.251761 |
| 48AspH | 221.63018 | 21.36503 | 5.60859 | 10 | A exp(-Bx) | 37424.6 | 0.00451 | 1675.26379 | 0.000430988 | 4.51 | 0.430988 |
| 49AspH | 221.58572 | 17.42797 | 4.87206 | 10 | A exp(-Bx) | 39655 | 0.00451 | 1452.84753 | 0.000352778 | 4.51 | 0.352778 |
| 50IleH | 231.601 | 17.63286 | 4.81046 | 10 | A exp(-Bx) | 41124.4 | 0.00432 | 1423.65637 | 0.000326848 | 4.32 | 0.326848 |
| 51GluH | 187.73046 | 15.35048 | 6.03192 | 10 | A exp(-Bx) | 45226.7 | 0.00533 | 1870.84937 | 0.00043269 | 5.33 | 0.43269 |
| 52GlnH | 198.5085 | 19.55506 | 6.48511 | 10 | A exp(-Bx) | 41010.2 | 0.00504 | 1984.86511 | 0.000491526 | 5.04 | 0.491526 |
| 53TrpH | 206.12666 | 10.50043 | 5.78982 | 10 | A exp(-Bx) | 71022.8 | 0.00485 | 1756.86609 | 0.000246499 | 4.85 | 0.246499 |
| 55ThrH | 216.574 | 11.96751 | 3.07681 | 10 | A exp(-Bx) | 35331.3 | 0.00462 | 923.56372 | 0.000254373 | 4.62 | 0.254373 |
| 56GluH | 244.69129 | 13.6624 | 3.08724 | 10 | A exp(-Bx) | 36660.9 | 0.00409 | 903.96674 | 0.00022748 | 4.09 | 0.22748 |
| 59GlyH | 218.31247 | 10.59311 | 2.83486 | 10 | A exp(-Bx) | 37140.6 | 0.00458 | 849.46857 | 0.000221742 | 4.58 | 0.221742 |
| 61AspH | 191.59836 | 10.32621 | 3.09821 | 10 | A exp(-Bx) | 35269.9 | 0.00522 | 956.19379 | 0.00028048 | 5.22 | 0.28048 |
| 62GluH | 246.07671 | 17.47901 | 3.74126 | 10 | A exp(-Bx) | 35006.4 | 0.00406 | 1092.32812 | 0.000287211 | 4.06 | 0.287211 |
| 63AlaH | 237.90494 | 13.46753 | 3.44147 | 10 | A exp(-Bx) | 39871.9 | 0.0042 | 1013.2287 | 0.00023719 | 4.2 | 0.23719 |
| 65ArgH | 230.41895 | 19.53167 | 4.58261 | 10 | A exp(-Bx) | 35176.2 | 0.00434 | 1357.86829 | 0.000365271 | 4.34 | 0.365271 |
| 66MetH | 233.71532 | 17.66131 | 2.85561 | 10 | A exp(-Bx) | 24679.6 | 0.00428 | 843.7052 | 0.000321506 | 4.28 | 0.321506 |
| 68GluH | 205.36487 | 15.07223 | 4.73909 | 10 | A exp(-Bx) | 40420.6 | 0.00487 | 1439.25464 | 0.000355471 | 4.87 | 0.355471 |
| 69AlaH | 287.23195 | 13.53524 | 2.279 | 10 | A exp(-Bx) | 34357.5 | 0.00348 | 647.17084 | 0.000163697 | 3.48 | 0.163697 |
| 70AlaH | 261.84891 | 12.14438 | 3.14513 | 10 | A exp(-Bx) | 46210.2 | 0.00382 | 909.16437 | 0.000176743 | 3.82 | 0.176743 |
| 72ArgH | 466.55034 | 49.31039 | 5.45361 | 10 | A exp(-Bx) | 49686.1 | 0.00214 | 1452.81372 | 0.000224063 | 2.14 | 0.224063 |

#### Table S13. $T_2$ relaxation for p53-TAD$^{N29K/N30D}$.

| Assign F1 | Time Constant | TC Error | Fit Error | Num Peaks | Function | Fit Param A | Fit Param B | Param Error A | Param Error B | (1/s) R2 | (1/s) R2 error |
|---|---|---|---|---|---|---|---|---|---|---|---|
| 1MetH* | 245.38251 | 17.96961 | 3.34935 | 10 | A exp(-Bx) | 4.29E+04 | 0.00408 | 1383.33765 | 2.97E-04 | 4.08 | 2.97E-01 |
| 2GluH | 255.326 | 27.84731 | 6.49738 | 10 | A exp(-Bx) | 5.63E+04 | 0.00392 | 2621.28394 | 4.22E-04 | 3.92 | 4.22E-01 |
| 3GluH | 276.50955 | 22.86657 | 5.53212 | 10 | A exp(-Bx) | 6.64E+04 | 0.00362 | 2239.67456 | 2.97E-04 | 3.62 | 2.97E-01 |
| 5GlnH | 230.63843 | 19.18614 | 5.01032 | 10 | A exp(-Bx) | 5.54E+04 | 0.00434 | 2098.56836 | 3.58E-04 | 4.34 | 3.58E-01 |
| 6SerH | 230.63925 | 19.85394 | 3.93135 | 10 | A exp(-Bx) | 4.20E+04 | 0.00434 | 1643.76941 | 3.71E-04 | 4.34 | 3.71E-01 |
| 7AspH | 232.66557 | 15.90152 | 3.11331 | 10 | A exp(-Bx) | 4.19E+04 | 0.0043 | 1299.42407 | 2.92E-04 | 4.3 | 2.92E-01 |
| 9SerH | 215.89175 | 14.50497 | 3.55621 | 10 | A exp(-Bx) | 4.75E+04 | 0.00463 | 1510.21338 | 3.10E-04 | 4.63 | 3.10E-01 |
| 10ValH | 225.39168 | 17.09151 | 4.60182 | 10 | A exp(-Bx) | 5.53E+04 | 0.00444 | 1936.5863 | 3.35E-04 | 4.44 | 3.35E-01 |
| 11GluH | 243.71883 | 11.77412 | 2.80223 | 10 | A exp(-Bx) | 5.42E+04 | 0.0041 | 1158.89197 | 1.98E-04 | 4.1 | 1.98E-01 |
| 14LeuH | 238.71043 | 16.74149 | 4.17709 | 10 | A exp(-Bx) | 5.53E+04 | 0.00419 | 1734.51025 | 2.92E-04 | 4.19 | 2.92E-01 |
| 15SerH | 225.31955 | 16.81248 | 3.30654 | 10 | A exp(-Bx) | 4.04E+04 | 0.00444 | 1391.56226 | 3.29E-04 | 4.44 | 3.29E-01 |
| 18ThrH | 227.74669 | 14.54633 | 3.53386 | 10 | A exp(-Bx) | 5.05E+04 | 0.00439 | 1483.94763 | 2.79E-04 | 4.39 | 2.79E-01 |
| 19PheH | 222.97468 | 17.27488 | 3.37534 | 10 | A exp(-Bx) | 3.96E+04 | 0.00448 | 1423.61438 | 3.45E-04 | 4.48 | 3.45E-01 |
| 20SerH | 211.64276 | 15.82194 | 2.62407 | 10 | A exp(-Bx) | 3.13E+04 | 0.00472 | 1117.30823 | 3.51E-04 | 4.72 | 3.51E-01 |
| 21AspH | 188.21023 | 10.95598 | 2.09732 | 10 | A exp(-Bx) | 3.11E+04 | 0.00531 | 917.63312 | 3.08E-04 | 5.31 | 3.08E-01 |
| 22LeuH | 192.96747 | 12.8983 | 5.26034 | 10 | A exp(-Bx) | 6.85E+04 | 0.00518 | 2291.4729 | 3.45E-04 | 5.18 | 3.45E-01 |
| 23TrpH | 176.40449 | 10.75936 | 3.91046 | 10 | A exp(-Bx) | 5.48E+04 | 0.00567 | 1741.76147 | 3.44E-04 | 5.67 | 3.44E-01 |
| 24LysH | 187.34369 | 12.23356 | 4.2852 | 10 | A exp(-Bx) | 5.67E+04 | 0.00534 | 1880.0249 | 3.47E-04 | 5.34 | 3.47E-01 |
| 25LeuH | 205.91569 | 16.94385 | 5.33549 | 10 | A exp(-Bx) | 5.75E+04 | 0.00486 | 2289.52002 | 3.97E-04 | 4.86 | 3.97E-01 |
| 26LeuH | 216.34663 | 13.36222 | 3.89381 | 10 | A exp(-Bx) | 5.65E+04 | 0.00462 | 1650.04236 | 2.84E-04 | 4.62 | 2.84E-01 |
| 28GluH | 210.96517 | 12.92591 | 3.9638 | 10 | A exp(-Bx) | 5.75E+04 | 0.00474 | 1688.95093 | 2.89E-04 | 4.74 | 2.89E-01 |
| 29LysH | 212.25553 | 11.93689 | 2.84001 | 10 | A exp(-Bx) | 4.50E+04 | 0.00471 | 1210.53748 | 2.64E-04 | 4.71 | 2.64E-01 |
| 30AspH | 204.33318 | 11.95442 | 3.47559 | 10 | A exp(-Bx) | 5.24E+04 | 0.00489 | 1493.78638 | 2.85E-04 | 4.89 | 2.85E-01 |
| 31ValH | 218.50875 | 13.24141 | 4.17225 | 10 | A exp(-Bx) | 6.20E+04 | 0.00458 | 1766.98657 | 2.76E-04 | 4.58 | 2.76E-01 |
| 32LeuH | 259.07448 | 16.97088 | 6.43012 | 10 | A exp(-Bx) | 9.43E+04 | 0.00386 | 2633.54688 | 2.52E-04 | 3.86 | 2.52E-01 |
| 33SerH | 199.66798 | 11.5419 | 2.95263 | 10 | A exp(-Bx) | 4.47E+04 | 0.00501 | 1273.83618 | 2.89E-04 | 5.01 | 2.89E-01 |
| 35LeuH | 312.38784 | 33.98574 | 3.74694 | 10 | A exp(-Bx) | 3.65E+04 | 0.0032 | 1486.69617 | 3.44E-04 | 3.2 | 3.44E-01 |
| 37SerH | 233.96802 | 12.36086 | 2.06325 | 10 | A exp(-Bx) | 3.60E+04 | 0.00427 | 861.67969 | 2.25E-04 | 4.27 | 2.25E-01 |
| 38GlnH | 216.72201 | 14.73724 | 2.47413 | 10 | A exp(-Bx) | 3.26E+04 | 0.00461 | 1048.04883 | 3.12E-04 | 4.61 | 3.12E-01 |
| 39AlaH | 260.33795 | 14.64216 | 4.38084 | 10 | A exp(-Bx) | 7.49E+04 | 0.00384 | 1792.60852 | 2.15E-04 | 3.84 | 2.15E-01 |
| 40MetH | 233.41899 | 17.47942 | 3.59248 | 10 | A exp(-Bx) | 4.43E+04 | 0.00428 | 1501.06848 | 3.19E-04 | 4.28 | 3.19E-01 |
| 41AspH | 233.53408 | 17.25379 | 4.19067 | 10 | A exp(-Bx) | 5.23E+04 | 0.00428 | 1750.8147 | 3.15E-04 | 4.28 | 3.15E-01 |
| 42AspH | 221.46155 | 11.62547 | 4.50417 | 10 | A exp(-Bx) | 7.75E+04 | 0.00452 | 1902.14075 | 2.36E-04 | 4.52 | 2.36E-01 |
| 43LeuH | 234.89225 | 17.99152 | 4.56573 | 10 | A exp(-Bx) | 5.51E+04 | 0.00426 | 1901.98157 | 3.24E-04 | 4.26 | 3.24E-01 |
| 44MetH | 221.46468 | 13.2705 | 4.76595 | 10 | A exp(-Bx) | 7.18E+04 | 0.00452 | 2009.57422 | 2.70E-04 | 4.52 | 2.70E-01 |
| 45LeuH | 245.5063 | 19.67325 | 4.90743 | 10 | A exp(-Bx) | 5.77E+04 | 0.00407 | 2030.27197 | 3.24E-04 | 4.07 | 3.24E-01 |
| 46SerH | 214.2397 | 17.89945 | 5.47009 | 10 | A exp(-Bx) | 5.88E+04 | 0.00467 | 2326.48169 | 3.87E-04 | 4.67 | 3.87E-01 |
| 48AspH | 224.01455 | 9.66963 | 2.53422 | 10 | A exp(-Bx) | 5.32E+04 | 0.00446 | 1067.83545 | 1.92E-04 | 4.46 | 1.92E-01 |
| 49AspH | 215.25915 | 8.91242 | 2.65422 | 10 | A exp(-Bx) | 5.73E+04 | 0.00465 | 1127.87891 | 1.92E-04 | 4.65 | 1.92E-01 |
| 50IleH | 243.52879 | 9.48117 | 2.57725 | 10 | A exp(-Bx) | 6.18E+04 | 0.00411 | 1067.89136 | 1.60E-04 | 4.11 | 1.60E-01 |
| 51GluH | 219.14485 | 12.08761 | 3.13239 | 10 | A exp(-Bx) | 5.11E+04 | 0.00456 | 1325.98059 | 2.51E-04 | 4.56 | 2.51E-01 |
| 52GlnH | 204.57521 | 11.45935 | 3.55687 | 10 | A exp(-Bx) | 5.60E+04 | 0.00489 | 1528.54297 | 2.73E-04 | 4.89 | 2.73E-01 |
| 53TrpH | 206.39397 | 12.94653 | 7.56462 | 10 | A exp(-Bx) | 1.07E+05 | 0.00485 | 3244.42969 | 3.03E-04 | 4.85 | 3.03E-01 |
| 54PheH | 206.39397 | 12.94653 | 7.56462 | 10 | A exp(-Bx) | 1.07E+05 | 0.00485 | 3244.42969 | 3.03E-04 | 4.85 | 3.03E-01 |
| 55ThrH | 222.62751 | 15.61277 | 3.58396 | 10 | A exp(-Bx) | 4.63E+04 | 0.00449 | 1511.86292 | 3.13E-04 | 4.49 | 3.13E-01 |
| 56GluH | 230.91279 | 20.02485 | 5.27718 | 10 | A exp(-Bx) | 5.60E+04 | 0.00433 | 2209.80786 | 3.73E-04 | 4.33 | 3.73E-01 |
| 57AspH | 232.58486 | 17.04169 | 4.78656 | 10 | A exp(-Bx) | 6.01E+04 | 0.0043 | 1997.94019 | 3.13E-04 | 4.3 | 3.13E-01 |
| 59GlyH | 241.0229 | 19.64926 | 4.73705 | 10 | A exp(-Bx) | 5.42E+04 | 0.00415 | 1963.30627 | 3.36E-04 | 4.15 | 3.36E-01 |
| 61AspH | 205.98449 | 14.56577 | 4.23031 | 10 | A exp(-Bx) | 5.21E+04 | 0.00485 | 1785.50391 | 3.42E-04 | 4.85 | 3.42E-01 |
| 62GluH | 231.27796 | 15.91861 | 3.5725 | 10 | A exp(-Bx) | 4.76E+04 | 0.00432 | 1492.88794 | 2.96E-04 | 4.32 | 2.96E-01 |
| 63AlaH | 274.47525 | 17.44101 | 3.49972 | 10 | A exp(-Bx) | 5.43E+04 | 0.00364 | 1418.41296 | 2.31E-04 | 3.64 | 2.31E-01 |
| 65ArgH | 218.91843 | 15.60085 | 6.85694 | 10 | A exp(-Bx) | 8.68E+04 | 0.00457 | 2903.26831 | 3.24E-04 | 4.57 | 3.24E-01 |
| 66MetH | 235.1849 | 15.37745 | 2.69934 | 10 | A exp(-Bx) | 3.82E+04 | 0.00425 | 1126.18457 | 2.77E-04 | 4.25 | 2.77E-01 |
| 68GluH | 217.53521 | 17.42971 | 5.23643 | 10 | A exp(-Bx) | 5.88E+04 | 0.0046 | 2216.3772 | 3.66E-04 | 4.6 | 3.66E-01 |
| 69AlaH | 290.07033 | 47.32348 | 13.30983 | 10 | A exp(-Bx) | 8.45E+04 | 0.00345 | 5345.0708 | 5.48E-04 | 3.45 | 5.48E-01 |
| 70AlaH | 315.74921 | 27.33071 | 5.10124 | 10 | A exp(-Bx) | 6.26E+04 | 0.00317 | 2021.10986 | 2.72E-04 | 3.17 | 2.72E-01 |
| 72ArgH | 247.0044 | 17.29565 | 5.36223 | 10 | A exp(-Bx) | 7.21E+04 | 0.00405 | 2211.8916 | 2.82E-04 | 4.05 | 2.82E-01 |
| 73ValH | 386.31526 | 34.02149 | 6.17744 | 10 | A exp(-Bx) | 8.39E+04 | 0.00259 | 2378.79248 | 2.26E-04 | 2.59 | 2.26E-01 |

#### Table S14. $T_2$ relaxation for p53-TAD$^{D49Y}$.

| Assign F1 | Time Constant | TC Error | Fit Error | Num Peaks | Function | Fit Param A | Fit Param B | Param Error A | Param Error B | (1/s) R2 | (1/s) R2 Error |
|---|---|---|---|---|---|---|---|---|---|---|---|
| 1MetH* | 201.91336 | 9.80916 | 0.61796 | 10 | A exp(-Bx) | 2.53E+04 | 0.00495 | 603.15265 | 2.40E-04 | 4.95 | 2.40E-01 |
| 2GluH | 209.25276 | 13.45119 | 1.15705 | 10 | A exp(-Bx) | 3.62E+04 | 0.00478 | 1120.29382 | 3.06E-04 | 4.78 | 3.06E-01 |
| 3GluH | 221.76927 | 17.18033 | 1.57907 | 10 | A exp(-Bx) | 4.19E+04 | 0.00451 | 1509.88354 | 3.47E-04 | 4.51 | 3.47E-01 |
| 5GlnH | 199.94079 | 12.15473 | 0.95635 | 10 | A exp(-Bx) | 3.12E+04 | 0.005 | 934.0127 | 3.03E-04 | 5 | 3.03E-01 |
| 6SerH | 178.90992 | 12.01124 | 0.82045 | 10 | A exp(-Bx) | 2.37E+04 | 0.00559 | 824.53796 | 3.74E-04 | 5.59 | 3.74E-01 |
| 7AspH | 208.07452 | 10.45341 | 0.70428 | 10 | A exp(-Bx) | 2.81E+04 | 0.00481 | 681.62402 | 2.41E-04 | 4.81 | 2.41E-01 |
| 9SerH | 205.25958 | 9.42126 | 0.63539 | 10 | A exp(-Bx) | 2.76E+04 | 0.00487 | 617.85437 | 2.23E-04 | 4.87 | 2.23E-01 |
| 10ValH | 181.25668 | 8.99469 | 0.85531 | 10 | A exp(-Bx) | 3.34E+04 | 0.00552 | 855.36481 | 2.73E-04 | 5.52 | 2.73E-01 |
| 11GluH | 228.84372 | 8.45491 | 0.57949 | 10 | A exp(-Bx) | 3.24E+04 | 0.00437 | 549.55048 | 1.61E-04 | 4.37 | 1.61E-01 |
| 14LeuH | 195.1727 | 9.97722 | 1.08175 | 10 | A exp(-Bx) | 4.17E+04 | 0.00512 | 1062.45569 | 2.61E-04 | 5.12 | 2.61E-01 |
| 15SerH | 188.00555 | 7.71609 | 0.47095 | 10 | A exp(-Bx) | 2.24E+04 | 0.00532 | 466.72507 | 2.18E-04 | 5.32 | 2.18E-01 |
| 18ThrH | 188.79606 | 6.09401 | 0.45239 | 10 | A exp(-Bx) | 2.73E+04 | 0.0053 | 447.87103 | 1.71E-04 | 5.3 | 1.71E-01 |
| 19PheH | 172.43668 | 9.26003 | 0.60046 | 10 | A exp(-Bx) | 2.15E+04 | 0.0058 | 609.23669 | 3.11E-04 | 5.8 | 3.11E-01 |
| 20SerH | 164.54319 | 5.7391 | 0.32206 | 10 | A exp(-Bx) | 1.77E+04 | 0.00608 | 330.8584 | 2.12E-04 | 6.08 | 2.12E-01 |
| 21AspH | 160.9516 | 11.19644 | 0.73241 | 10 | A exp(-Bx) | 2.02E+04 | 0.00621 | 757.14111 | 4.30E-04 | 6.21 | 4.30E-01 |
| 22LeuH | 155.46141 | 6.75431 | 0.83484 | 10 | A exp(-Bx) | 3.66E+04 | 0.00643 | 871.61572 | 2.79E-04 | 6.43 | 2.79E-01 |
| 23TrpH | 156.85869 | 8.0657 | 0.93074 | 10 | A exp(-Bx) | 3.45E+04 | 0.00638 | 967.74799 | 3.27E-04 | 6.38 | 3.27E-01 |
| 24LysH | 144.91832 | 5.42147 | 0.68799 | 10 | A exp(-Bx) | 3.49E+04 | 0.0069 | 733.47498 | 2.58E-04 | 6.9 | 2.58E-01 |
| 25LeuH | 162.22236 | 10.77361 | 1.10622 | 10 | A exp(-Bx) | 3.19E+04 | 0.00616 | 1141.06909 | 4.08E-04 | 6.16 | 4.08E-01 |
| 26LeuH | 184.90326 | 12.96969 | 1.19644 | 10 | A exp(-Bx) | 3.33E+04 | 0.00541 | 1192.30249 | 3.78E-04 | 5.41 | 3.78E-01 |
| 28GluH | 158.43491 | 3.84216 | 0.42323 | 10 | A exp(-Bx) | 3.32E+04 | 0.00631 | 439.46954 | 1.53E-04 | 6.31 | 1.53E-01 |
| 29AsnH | 177.21487 | 11.79349 | 0.64934 | 10 | A exp(-Bx) | 1.89E+04 | 0.00564 | 654.16003 | 3.74E-04 | 5.64 | 3.74E-01 |
| 30AsnH | 189.04393 | 8.44065 | 0.32394 | 10 | A exp(-Bx) | 1.42E+04 | 0.00529 | 321.12189 | 2.36E-04 | 5.29 | 2.36E-01 |
| 31ValH | 190.67305 | 11.24465 | 1.02906 | 10 | A exp(-Bx) | 3.43E+04 | 0.00524 | 1017.84241 | 3.08E-04 | 5.24 | 3.08E-01 |
| 32LeuH | 252.61426 | 15.30807 | 0.77059 | 10 | A exp(-Bx) | 2.74E+04 | 0.00396 | 718.02948 | 2.39E-04 | 3.96 | 2.39E-01 |
| 33SerH | 181.78749 | 8.90264 | 0.76014 | 10 | A exp(-Bx) | 3.01E+04 | 0.0055 | 760.84814 | 2.69E-04 | 5.5 | 2.69E-01 |
| 35LeuH | 220.95349 | 15.81428 | 1.0388 | 10 | A exp(-Bx) | 2.97E+04 | 0.00453 | 994.05304 | 3.22E-04 | 4.53 | 3.22E-01 |
| 37SerH | 206.00383 | 9.74086 | 0.45133 | 10 | A exp(-Bx) | 1.91E+04 | 0.00485 | 438.5264 | 2.29E-04 | 4.85 | 2.29E-01 |
| 38GlnH | 175.45304 | 17.94481 | 0.92132 | 10 | A exp(-Bx) | 1.75E+04 | 0.0057 | 930.55945 | 5.77E-04 | 5.7 | 5.77E-01 |
| 39AlaH | 218.92571 | 8.45262 | 0.52233 | 10 | A exp(-Bx) | 2.75E+04 | 0.00457 | 500.72217 | 1.76E-04 | 4.57 | 1.76E-01 |
| 40MetH | 174.45541 | 6.69595 | 0.63235 | 10 | A exp(-Bx) | 3.17E+04 | 0.00573 | 638.62634 | 2.20E-04 | 5.73 | 2.20E-01 |
| 41AspH | 183.26259 | 6.48125 | 0.60921 | 10 | A exp(-Bx) | 3.35E+04 | 0.00546 | 608.54639 | 1.93E-04 | 5.46 | 1.93E-01 |
| 42AspH | 175.8989 | 10.22846 | 1.25286 | 10 | A exp(-Bx) | 4.16E+04 | 0.00569 | 1264.39819 | 3.29E-04 | 5.69 | 3.29E-01 |
| 43LeuH | 180.93963 | 9.31577 | 0.96108 | 10 | A exp(-Bx) | 3.62E+04 | 0.00553 | 963.10004 | 2.84E-04 | 5.53 | 2.84E-01 |
| 44MetH | 168.42187 | 7.56647 | 0.87044 | 10 | A exp(-Bx) | 3.72E+04 | 0.00594 | 888.57666 | 2.66E-04 | 5.94 | 2.66E-01 |
| 45LeuH | 158.84627 | 6.23796 | 0.70043 | 10 | A exp(-Bx) | 3.40E+04 | 0.0063 | 725.66681 | 2.47E-04 | 6.3 | 2.47E-01 |
| 46SerH | 168.74633 | 7.54367 | 0.81533 | 10 | A exp(-Bx) | 3.50E+04 | 0.00593 | 832.00952 | 2.64E-04 | 5.93 | 2.64E-01 |
| 48AspH | 167.6425 | 7.21933 | 0.83131 | 10 | A exp(-Bx) | 3.70E+04 | 0.00597 | 849.81183 | 2.56E-04 | 5.97 | 2.56E-01 |
| 49TyrH | 165.38924 | 7.85994 | 0.82603 | 10 | A exp(-Bx) | 3.33E+04 | 0.00605 | 847.40479 | 2.87E-04 | 6.05 | 2.87E-01 |
| 50IleH | 186.2492 | 10.03343 | 0.82925 | 10 | A exp(-Bx) | 3.00E+04 | 0.00537 | 824.91003 | 2.88E-04 | 5.37 | 2.88E-01 |
| 51GluH | 176.25793 | 6.54999 | 0.53287 | 10 | A exp(-Bx) | 2.77E+04 | 0.00567 | 537.56812 | 2.11E-04 | 5.67 | 2.11E-01 |
| 52GlnH | 174.69905 | 9.91827 | 1.18725 | 10 | A exp(-Bx) | 4.03E+04 | 0.00572 | 1198.59412 | 3.24E-04 | 5.72 | 3.24E-01 |
| 53TrpH | 169.58101 | 9.45755 | 1.15103 | 10 | A exp(-Bx) | 3.97E+04 | 0.0059 | 1172.85657 | 3.28E-04 | 5.9 | 3.28E-01 |
| 54PheH | 165.72939 | 6.80111 | 0.97304 | 10 | A exp(-Bx) | 4.54E+04 | 0.00603 | 997.8017 | 2.47E-04 | 6.03 | 2.47E-01 |
| 55ThrH | 185.11796 | 7.36622 | 0.52601 | 10 | A exp(-Bx) | 2.57E+04 | 0.0054 | 524.1217 | 2.15E-04 | 5.4 | 2.15E-01 |
| 56GluH | 191.36004 | 7.56505 | 0.58099 | 10 | A exp(-Bx) | 2.88E+04 | 0.00523 | 574.16492 | 2.06E-04 | 5.23 | 2.06E-01 |
| 57AspH | 208.21384 | 9.50333 | 0.75493 | 10 | A exp(-Bx) | 3.31E+04 | 0.0048 | 730.53546 | 2.19E-04 | 4.8 | 2.19E-01 |
| 59GlyH | 218.43134 | 20.80291 | 1.4187 | 10 | A exp(-Bx) | 3.05E+04 | 0.00458 | 1360.90552 | 4.32E-04 | 4.58 | 4.32E-01 |
| 61AspH | 185.57142 | 12.23238 | 0.93098 | 10 | A exp(-Bx) | 2.76E+04 | 0.00539 | 927.07111 | 3.54E-04 | 5.39 | 3.54E-01 |
| 62GluH | 191.07731 | 10.64702 | 1.35681 | 10 | A exp(-Bx) | 4.78E+04 | 0.00523 | 1341.33459 | 2.91E-04 | 5.23 | 2.91E-01 |
| 63AlaH | 229.84556 | 11.05835 | 0.72444 | 10 | A exp(-Bx) | 3.12E+04 | 0.00435 | 687.57971 | 2.09E-04 | 4.35 | 2.09E-01 |
| 65ArgH | 185.23438 | 7.08022 | 0.98857 | 10 | A exp(-Bx) | 5.04E+04 | 0.0054 | 984.86377 | 2.06E-04 | 5.4 | 2.06E-01 |
| 66MetH | 201.76571 | 10.25935 | 0.58635 | 10 | A exp(-Bx) | 2.29E+04 | 0.00496 | 572.40771 | 2.51E-04 | 4.96 | 2.51E-01 |
| 68GluH | 197.71922 | 11.24816 | 1.02927 | 10 | A exp(-Bx) | 3.58E+04 | 0.00506 | 1009.49365 | 2.87E-04 | 5.06 | 2.87E-01 |
| 69AlaH | 193.02455 | 9.15225 | 0.87791 | 10 | A exp(-Bx) | 3.64E+04 | 0.00518 | 865.93793 | 2.45E-04 | 5.18 | 2.45E-01 |
| 70AlaH | 269.20972 | 14.8328 | 0.84834 | 10 | A exp(-Bx) | 3.40E+04 | 0.00371 | 781.37585 | 2.04E-04 | 3.71 | 2.04E-01 |
| 72ArgH | 216.26782 | 10.47035 | 1.03518 | 10 | A exp(-Bx) | 4.27E+04 | 0.00462 | 978.58807 | 2.23E-04 | 4.62 | 2.23E-01 |
| 73ValH | 334.57403 | 21.95869 | 1.44975 | 10 | A exp(-Bx) | 5.47E+04 | 0.00299 | 1289.2395 | 1.95E-04 | 2.99 | 1.95E-01 |

#### Table S15. $T_2$ relaxation for p53-TAD$^{W53G}$.

| Assign F1 | Assign F2 | Time Constant | TC Error | Fit Error | Num Peaks | Function | Fit Param A | Fit Param B | Param Error A | Param Error B | (1/s) R2 | (1/s) R2 Error |
|---|---|---|---|---|---|---|---|---|---|---|---|---|
| 1MetH* | 1MetN | 185.28338 | 3.84804 | 1.14089 | 8 | A exp(-Bx) | 34498.1 | 0.0054 | 405.8779 | 0.000112042 | 5.4 | 0.112042 |
| 2GluH | 2GluN | 195.22836 | 6.55074 | 2.09718 | 8 | A exp(-Bx) | 40364 | 0.00512 | 749.41827 | 0.000171679 | 5.12 | 0.171679 |
| 3GluH | 3GluN | 214.66299 | 7.30526 | 2.80124 | 8 | A exp(-Bx) | 54544.2 | 0.00466 | 980.88025 | 0.00015835 | 4.66 | 0.15835 |
| 5GlnH | 5GlnN | 187.33003 | 7.64833 | 2.86867 | 8 | A exp(-Bx) | 44896.6 | 0.00534 | 1032.33154 | 0.000217585 | 5.34 | 0.217585 |
| 6SerH | 6SerN | 170.38926 | 3.83427 | 1.10386 | 8 | A exp(-Bx) | 30847 | 0.00587 | 406.23911 | 0.000132001 | 5.87 | 0.132001 |
| 7AspH | 7AspN | 196.93052 | 8.97067 | 3.23506 | 8 | A exp(-Bx) | 45935.5 | 0.00508 | 1152.14807 | 0.000230834 | 5.08 | 0.230834 |
| 9SerH | 9SerN | 182.71083 | 2.84437 | 0.95771 | 8 | A exp(-Bx) | 39109.9 | 0.00547 | 346.97803 | 8.51829E-05 | 5.47 | 0.0851829 |
| 10ValH | 10ValN | 178.37624 | 6.32351 | 2.91766 | 8 | A exp(-Bx) | 52153.7 | 0.00561 | 1062.76538 | 0.00019849 | 5.61 | 0.19849 |
| 11GluH | 11GluN | 216.87314 | 5.22047 | 1.787 | 8 | A exp(-Bx) | 48596.6 | 0.00461 | 615.24512 | 0.000110929 | 4.61 | 0.110929 |
| 14LeuH | 14LeuN | 190.84482 | 8.38931 | 3.75478 | 8 | A exp(-Bx) | 54907.1 | 0.00524 | 1348.07275 | 0.000229895 | 5.24 | 0.229895 |
| 15SerH | 15SerN | 172.33888 | 5.80176 | 1.85174 | 8 | A exp(-Bx) | 34659.7 | 0.0058 | 679.61633 | 0.00019512 | 5.8 | 0.19512 |
| 16GlnH | 16GlnN | 191.63512 | 5.89109 | 3.25582 | 8 | A exp(-Bx) | 68079 | 0.00522 | 1167.90808 | 0.000160264 | 5.22 | 0.160264 |
| 17GluH | 17GluN | 168.755 | 4.25615 | 3.23644 | 8 | A exp(-Bx) | 80488.4 | 0.00593 | 1191.93555 | 0.000149358 | 5.93 | 0.149358 |
| 18ThrH | 18ThrN | 171.96011 | 3.54857 | 1.25937 | 8 | A exp(-Bx) | 38415.6 | 0.00582 | 462.43649 | 0.000119954 | 5.82 | 0.119954 |
| 19PheH | 19PheN | 157.56432 | 6.02949 | 1.96835 | 8 | A exp(-Bx) | 32141.6 | 0.00635 | 737.91187 | 0.00024251 | 6.35 | 0.24251 |
| 20SerH | 20SerN | 167.83705 | 6.52909 | 1.59404 | 8 | A exp(-Bx) | 25745.8 | 0.00596 | 588.68011 | 0.000231431 | 5.96 | 0.231431 |
| 21AspH | 21AspN | 148.5782 | 4.04835 | 1.2534 | 8 | A exp(-Bx) | 28677.7 | 0.00673 | 476.82962 | 0.000183251 | 6.73 | 0.183251 |
| 22LeuH | 22LeuN | 154.64785 | 4.9975 | 2.83741 | 8 | A exp(-Bx) | 54794.6 | 0.00647 | 1068.60767 | 0.000208743 | 6.47 | 0.208743 |
| 23TrpH | 23TrpN | 157.34966 | 5.37459 | 2.76946 | 8 | A exp(-Bx) | 50645.4 | 0.00636 | 1038.58069 | 0.000216824 | 6.36 | 0.216824 |
| 24LysH | 24LysN | 154.19067 | 3.029 | 1.55124 | 8 | A exp(-Bx) | 49171.1 | 0.00649 | 583.81311 | 0.000127355 | 6.49 | 0.127355 |
| 25LeuH | 25LeuN | 162.96187 | 5.77033 | 2.5622 | 8 | A exp(-Bx) | 45315.2 | 0.00614 | 952.77161 | 0.000217013 | 6.14 | 0.217013 |
| 26LeuH | 26LeuN | 180.20562 | 6.14435 | 3.06332 | 8 | A exp(-Bx) | 57030.2 | 0.00555 | 1113.34766 | 0.000188988 | 5.55 | 0.188988 |
| 28GluH | 28GluN | 160.08336 | 6.47423 | 3.51516 | 8 | A exp(-Bx) | 54296.2 | 0.00625 | 1310.8241 | 0.000252224 | 6.25 | 0.252224 |
| 29AsnH | 29AsnN | 166.89369 | 8.14634 | 2.03707 | 8 | A exp(-Bx) | 26188.5 | 0.00599 | 752.16089 | 0.000291777 | 5.99 | 0.291777 |
| 30AsnH | 30AsnN | 168.14413 | 3.64762 | 0.83148 | 8 | A exp(-Bx) | 23712.1 | 0.00595 | 302.4696 | 0.000128956 | 5.95 | 0.128956 |
| 31ValH | 31ValN | 185.14705 | 4.65692 | 1.87831 | 8 | A exp(-Bx) | 47551.4 | 0.0054 | 677.62531 | 0.000135766 | 5.4 | 0.135766 |
| 32LeuH | 32LeuN | 217.3834 | 5.90973 | 1.70824 | 8 | A exp(-Bx) | 41857 | 0.0046 | 597.65051 | 0.000124967 | 4.6 | 0.124967 |
| 33SerH | 33SerN | 174.03719 | 4.18205 | 1.38503 | 8 | A exp(-Bx) | 35817.8 | 0.00575 | 499.76981 | 0.000137992 | 5.75 | 0.137992 |
| 35LeuH | 35LeuN | 204.52568 | 9.24652 | 3.64108 | 8 | A exp(-Bx) | 52629.7 | 0.00489 | 1286.95081 | 0.000220596 | 4.89 | 0.220596 |
| 37SerH | 37SerN | 162.91432 | 4.34159 | 1.40103 | 8 | A exp(-Bx) | 32856.7 | 0.00614 | 520.25977 | 0.000163464 | 6.14 | 0.163464 |
| 38GlnH | 38GlnN | 169.06791 | 6.67394 | 1.49148 | 8 | A exp(-Bx) | 23414.4 | 0.00591 | 541.85168 | 0.000233123 | 5.91 | 0.233123 |
| 39AlaH | 39AlaN | 202.77089 | 6.50698 | 2.38588 | 8 | A exp(-Bx) | 47722.4 | 0.00493 | 832.6026 | 0.000158096 | 4.93 | 0.158096 |
| 40MetH | 40MetN | 177.63682 | 3.81967 | 1.45641 | 8 | A exp(-Bx) | 42215.9 | 0.00563 | 523.05762 | 0.000120993 | 5.63 | 0.120993 |
| 41AspH | 41AspN | 188.15031 | 6.63899 | 2.70303 | 8 | A exp(-Bx) | 48974.9 | 0.00531 | 971.82422 | 0.000187306 | 5.31 | 0.187306 |
| 42AspH | 42AspN | 193.9699 | 4.76873 | 2.12511 | 8 | A exp(-Bx) | 55617.2 | 0.00516 | 759.19849 | 0.000126669 | 5.16 | 0.126669 |
| 43LeuH | 43LeuN | 177.70911 | 4.00952 | 2.30138 | 8 | A exp(-Bx) | 64538.2 | 0.00563 | 838.86188 | 0.000126897 | 5.63 | 0.126897 |
| 44MetH | 44MetN | 194.2965 | 8.9535 | 4.39151 | 8 | A exp(-Bx) | 61435.4 | 0.00515 | 1568.33057 | 0.00023667 | 5.15 | 0.23667 |
| 45LeuH | 45LeuN | 204.05091 | 11.41632 | 4.84413 | 8 | A exp(-Bx) | 56695.2 | 0.0049 | 1715.74988 | 0.000273335 | 4.9 | 0.273335 |
| 46SerH | 46SerN | 175.99183 | 3.78794 | 1.67171 | 8 | A exp(-Bx) | 48995.3 | 0.00568 | 609.82233 | 0.000122241 | 5.68 | 0.122241 |
| 48AspH | 48AspN | 186.99264 | 5.62427 | 2.69008 | 8 | A exp(-Bx) | 57165.3 | 0.00535 | 969.80359 | 0.000160703 | 5.35 | 0.160703 |
| 49AspH | 49AspN | 184.55946 | 3.87209 | 3.85955 | 8 | A exp(-Bx) | 117223 | 0.00542 | 1395.49207 | 0.000113627 | 5.42 | 0.113627 |
| 50IleH | 50IleN | 201.00695 | 11.18939 | 7.12719 | 8 | A exp(-Bx) | 83471 | 0.00497 | 2531.54443 | 0.000276086 | 4.97 | 0.276086 |
| 51GluH | 51GluN | 232.71877 | 10.05916 | 3.99723 | 8 | A exp(-Bx) | 63102.5 | 0.0043 | 1379.07605 | 0.000185391 | 4.3 | 0.185391 |
| 52GlnH | 52GlnN | 183.67528 | 4.93188 | 2.25924 | 8 | A exp(-Bx) | 53496.2 | 0.00544 | 816.45203 | 0.000146083 | 5.44 | 0.146083 |
| 53GlyH | 53GlyN | 187.15446 | 7.40034 | 2.74242 | 8 | A exp(-Bx) | 44303.9 | 0.00534 | 987.09393 | 0.000210947 | 5.34 | 0.210947 |
| 55ThrH | 55ThrN | 175.78041 | 5.7019 | 1.97997 | 8 | A exp(-Bx) | 38572.8 | 0.00569 | 723.45599 | 0.000184341 | 5.69 | 0.184341 |
| 56GluH | 56GluN | 205.45956 | 7.12837 | 2.45325 | 8 | A exp(-Bx) | 46305 | 0.00487 | 867.7486 | 0.000168661 | 4.87 | 0.168661 |
| 57AspH | 57AspN | 194.1845 | 8.81868 | 3.79457 | 8 | A exp(-Bx) | 53854 | 0.00515 | 1355.30615 | 0.00023339 | 5.15 | 0.23339 |
| 59GlyH | 59GlyN | 186.97868 | 12.24136 | 5.92133 | 8 | A exp(-Bx) | 58009.9 | 0.00535 | 2135.04126 | 0.000348655 | 5.35 | 0.348655 |
| 61AspH | 61AspN | 189.54989 | 4.96831 | 2.11509 | 8 | A exp(-Bx) | 50928.6 | 0.00528 | 748.71442 | 0.000138186 | 5.28 | 0.138186 |
| 62GluH | 62GluN | 206.53977 | 8.43453 | 2.72197 | 8 | A exp(-Bx) | 43664.9 | 0.00484 | 960.23511 | 0.000197393 | 4.84 | 0.197393 |
| 63AlaH | 63AlaN | 237.92191 | 8.31334 | 2.73862 | 8 | A exp(-Bx) | 53998.6 | 0.0042 | 942.6889 | 0.000146682 | 4.2 | 0.146682 |
| 65ArgH | 65ArgN | 259.24269 | 126.91035 | 23.56236 | 8 | A exp(-Bx) | 41230.8 | 0.00386 | 7996.43066 | 0.00157 | 3.86 | 1.57 |
| 66MetH | 66MetN | 188.68543 | 6.39632 | 1.84731 | 8 | A exp(-Bx) | 34912.6 | 0.0053 | 664.81604 | 0.000179455 | 5.3 | 0.179455 |
| 68GluH | 68GluN | 195.74396 | 9.47195 | 3.7718 | 8 | A exp(-Bx) | 50352.9 | 0.00511 | 1344.96497 | 0.000246632 | 5.11 | 0.246632 |
| 69AlaH | 69AlaN | 245.64436 | 5.48172 | 1.34915 | 8 | A exp(-Bx) | 41437 | 0.00407 | 453.82339 | 9.08003E-05 | 4.07 | 0.0908003 |
| 70AlaH | 70AlaN | 263.07886 | 5.07533 | 1.34674 | 8 | A exp(-Bx) | 50200.1 | 0.0038 | 455.96994 | 7.33045E-05 | 3.8 | 0.0733045 |
| 72ArgH | 72ArgN | 198.46944 | 6.94604 | 2.9362 | 8 | A exp(-Bx) | 54416.1 | 0.00504 | 1045.73938 | 0.000176124 | 5.04 | 0.176124 |
| 73ValH | 73ValN | 342.37308 | 8.73464 | 2.50357 | 8 | A exp(-Bx) | 81167 | 0.00292 | 814.5694 | 7.44669E-05 | 2.92 | 0.0744669 |

**Table S16. NHNOE/NONOE ratios of p53-TAD$^{WT}$.** Values of NONOE and NHNOE intensity represent the integrated peak heights at those assigned positions within their respective spectra for p53-TAD$^{WT}$.

| Residue | NONOE Intensity | NHNOE Intensity | Intensity Ratio | Residue | NONOE Intensity | NHNOE Intensity | Intensity Ratio |
|---|---|---|---|---|---|---|---|
| 1 | -25523 | 21541 | -1.18 | 38 | -9648 | 16992 | -0.57 |
| 2 | -33492 | 32001 | -1.05 | 39 | -27634 | 36023 | -0.77 |
| 3 | -23470 | 37819 | -0.62 | 40 | -17458 | 24543 | -0.71 |
| 5 | -15200 | 20229 | -0.75 | 41 | -17262 | 30362 | -0.57 |
| 6 | -18038 | 20173 | -0.89 | 42 | -22754 | 49485 | -0.46 |
| 7 | -12933 | 30748 | -0.42 | 43 | -8100 | 36906 | -0.22 |
| 9 | -5266 | 25832 | -0.20 | 44 | -18773 | 40369 | -0.47 |
| 10 | -7593 | 14845 | -0.51 | 45 | -8191 | 32310 | -0.25 |
| 11 | -18148 | 32064 | -0.57 | 46 | -6462 | 25574 | -0.25 |
| 14 | -24774 | 39634 | -0.63 | 48 | -8504 | 33162 | -0.26 |
| 15 | -18545 | 25291 | -0.73 | 49 | -4530 | 32676 | -0.14 |
| 16 | -37597 | 36099 | -1.04 | 50 | -9623 | 27067 | -0.36 |
| 17 | -10911 | 43156 | -0.25 | 51 | -11298 | 38600 | -0.29 |
| 18 | -5646 | 17706 | -0.32 | 52 | -1822 | 33568 | -0.05 |
| 19 | -6842 | 15440 | -0.44 | 53 | -17203 | 54572 | -0.32 |
| 20 | -2394 | 14991 | -0.16 | 54 | -17203 | 54572 | -0.32 |
| 21 | -5308 | 19345 | -0.27 | 55 | -10606 | 23110 | -0.46 |
| 22 | -5222 | 20053 | -0.26 | 56 | -18583 | 39803 | -0.47 |
| 23 | -1622 | 14600 | -0.11 | 57 | -10505 | 43458 | -0.24 |
| 24 | -2038 | 14461 | -0.14 | 59 | -9940 | 38857 | -0.26 |
| 25 | -867 | 9374 | -0.09 | 61 | -9856 | 37046 | -0.27 |
| 26 | -2215 | 12248 | -0.18 | 62 | -4575 | 30764 | -0.15 |
| 28 | -4842 | 13550 | -0.36 | 63 | -23026 | 40489 | -0.57 |
| 29 | -4971 | 10399 | -0.48 | 65 | -10911 | 43156 | -0.25 |
| 30 | -6464 | 12385 | -0.52 | 66 | -6841 | 25645 | -0.27 |
| 31 | -4335 | 13496 | -0.32 | 68 | -13955 | 32268 | -0.43 |
| 32 | -12434 | 21355 | -0.58 | 69 | -37322 | 43141 | -0.87 |
| 33 | -11295 | 16824 | -0.67 | 70 | -38143 | 37412 | -1.02 |
| 35 | -10835 | 21307 | -0.51 | 72 | -37597 | 36099 | -1.04 |
| 37 | -9908 | 14468 | -0.68 | 73 | -19235 | 19421 | -0.99 |

**Table S17. NHNOE/NONOE ratios of p53-TAD$^{K24N}$.** Values of NONOE and NHNOE intensity represent the integrated peak heights at those assigned positions within their respective spectra for p53-TAD$^{K24N}$.

| Residue | NONOE Intensity | NHNOE Intensity | Intensity Ratio | Residue | NONOE Intensity | NHNOE Intensity | Intensity Ratio |
|---|---|---|---|---|---|---|---|
| 1 | -5.15E+05 | 6.32E+05 | -0.81419 | 38 | -1.61E+05 | 7.05E+05 | -0.22872 |
| 2 | -4.60E+05 | 8.33E+05 | -0.55213 | 39 | -3.28E+05 | 1.12E+06 | -0.29403 |
| 3 | -3.35E+05 | 1.06E+06 | -0.3152 | 40 | -3.04E+05 | 7.21E+05 | -0.42112 |
| 5 | -3.98E+05 | 1.13E+06 | -0.35226 | 41 | -1.76E+05 | 1.02E+06 | -0.17292 |
| 6 | -2.38E+05 | 1.03E+06 | -0.23193 | 42 | -1.63E+05 | 1.29E+06 | -0.12662 |
| 7 | -3.53E+05 | 9.70E+05 | -0.36375 | 43 | -2.54E+05 | 7.51E+05 | -0.33769 |
| 9 | -1.20E+05 | 1.06E+06 | -0.11348 | 44 | -1.79E+05 | 9.86E+05 | -0.18097 |
| 10 | -1.24E+05 | 2.02E+06 | -0.06124 | 45 | -2.21E+05 | 6.62E+05 | -0.33398 |
| 11 | -2.59E+05 | 1.03E+06 | -0.25215 | 46 | -1.59E+05 | 8.09E+05 | -0.1961 |
| 14 | -3.60E+05 | 1.03E+06 | -0.34833 | 48 | -1.44E+04 | 1.10E+06 | -0.01309 |
| 15 | -1.94E+05 | 1.02E+06 | -0.18958 | 49 | -8.01E+04 | 1.18E+06 | -0.06765 |
| 16 | -1.24E+06 | 1.47E+06 | -0.84456 | 50 | -1.21E+05 | 1.18E+06 | -0.10252 |
| 17 | -1.38E+05 | 1.20E+06 | -0.11477 | 51 | -1.55E+05 | 1.10E+06 | -0.1407 |
| 18 | -2.04E+05 | 9.35E+05 | -0.21819 | 52 | -6281.29639 | 9.95E+05 | -0.00631 |
| 19 | -1.37E+05 | 7.99E+05 | -0.17114 | 53 | -2.50E+04 | 1.92E+06 | -0.01303 |
| 20 | -1.36E+05 | 6.57E+05 | -0.20691 | 54 | - | - | - |
| 21 | -2.95E+04 | 8.39E+05 | -0.03517 | 55 | -7.50E+04 | 5.68E+05 | -0.13206 |
| 22 | 7.09E+04 | 1.32E+06 | 0.05356 | 56 | -1.14E+05 | 1.04E+06 | -0.10939 |
| 23 | 4.58E+04 | 1.25E+06 | 0.03665 | 59 | -2.06E+05 | 1.15E+06 | -0.17923 |
| 24 | 5.75E+04 | 6.22E+05 | 0.09253 | 61 | -1.12E+05 | 1.18E+06 | -0.09502 |
| 26 | -2.06E+05 | 9.63E+05 | -0.21427 | 62 | -1.59E+05 | 1.05E+06 | -0.15169 |
| 28 | -1.43E+05 | 8.75E+05 | -0.16382 | 63 | -2.48E+05 | 1.10E+06 | -0.22474 |
| 29 | -6.55E+04 | 8.22E+05 | -0.07974 | 65 | -2.63E+05 | 1.21E+06 | -0.21685 |
| 30 | -1.09E+05 | 5.53E+05 | -0.19792 | 66 | -1.72E+05 | 7.33E+05 | -0.2352 |
| 31 | -3.16E+05 | 1.11E+06 | -0.28536 | 68 | -3.19E+05 | 9.39E+05 | -0.33985 |
| 32 | -2.65E+05 | 8.35E+05 | -0.31729 | 69 | -5.00E+05 | 1.02E+06 | -0.48811 |
| 33 | -2.07E+05 | 9.80E+05 | -0.2107 | 70 | -6.51E+05 | 1.09E+06 | -0.59531 |
| 35 | -3.72E+05 | 1.09E+06 | -0.3419 | 72 | -1.24E+06 | 1.47E+06 | -0.84456 |
| 37 | -1.99E+05 | 4.77E+05 | -0.41657 | 73 | -1.88E+06 | 1.27E+06 | -1.48255 |

**Table S18. NHNOE/NONOE ratios of p53-TAD$^{N29K/N30D}$.** Values of NONOE and NHNOE intensity represent the integrated peak heights at those assigned positions within their respective spectra for p53-TAD$^{N29K/N30D}$.

| Residue | NONOE Intensity | NHNOE Intensity | Intensity Ratio | Residue | NONOE Intensity | NHNOE Intensity | Intensity Ratio |
|---|---|---|---|---|---|---|---|
| 1 | -1.37E+05 | 3.00E+05 | -0.45735 | 38 | -1.07E+05 | 7.81E+05 | -0.13757 |
| 2 | -1.73E+05 | 3.21E+05 | -0.5393 | 39 | -4.51E+05 | 1.44E+06 | -0.31387 |
| 3 | -8.71E+04 | 2.68E+05 | -0.32533 | 40 | -2.34E+05 | 8.74E+05 | -0.26813 |
| 5 | -5.82E+05 | 7.16E+05 | -0.81242 | 41 | -3.77E+05 | 9.96E+05 | -0.37832 |
| 6 | -3.38E+05 | 6.78E+05 | -0.49934 | 42 | -5.00E+05 | 1.49E+06 | -0.33473 |
| 7 | -2.30E+05 | 8.59E+05 | -0.26755 | 43 | -3.79E+04 | 1.04E+06 | -0.03632 |
| 9 | -1.50E+05 | 8.40E+05 | -0.17873 | 44 | -4.11E+05 | 1.11E+06 | -0.36994 |
| 10 | -2.10E+05 | 7.87E+05 | -0.26702 | 45 | -4.93E+04 | 9.66E+05 | -0.05104 |
| 11 | -1.94E+05 | 9.10E+05 | -0.21283 | 46 | -1.70E+05 | 8.80E+05 | -0.19281 |
| 14 | -1.55E+05 | 1.05E+06 | -0.14728 | 48 | -1.52E+05 | 1.06E+06 | -0.14333 |
| 15 | -1.66E+05 | 8.12E+05 | -0.20424 | 49 | | | |
| 16 | -6.69E+05 | 1.26E+06 | -0.52874 | 50 | -1.11E+05 | 1.06E+06 | -0.10425 |
| 17 | -7.14E+04 | 1.41E+06 | -0.05072 | 51 | 413.19424 | 1.05E+06 | 3.95E-04 |
| 18 | -8.11E+04 | 8.42E+05 | -0.09632 | 52 | 2.41E+04 | 9.09E+05 | 0.02655 |
| 19 | -1.12E+05 | 9.18E+05 | -0.12214 | 53 | -9.52E+04 | 1.84E+06 | -0.05187 |
| 20 | 7251.70557 | 7.17E+05 | 0.01012 | 54 | | | |
| 21 | 1.87E+04 | 8.24E+05 | 0.02276 | 55 | -9.22E+04 | 7.88E+05 | -0.11704 |
| 22 | 4.83E+04 | 1.12E+06 | 0.04293 | 56 | -1.99E+05 | 9.79E+05 | -0.20362 |
| 23 | 1.27E+05 | 9.78E+05 | 0.1298 | 57 | 3.39E+04 | 9.50E+05 | 0.03572 |
| 24 | 2.19E+04 | 8.93E+05 | 0.02454 | 59 | 3.04E+04 | 8.32E+05 | 0.03651 |
| 25 | -5.99E+04 | 9.19E+05 | -0.06516 | 61 | -7.65E+04 | 9.23E+05 | -0.0829 |
| 26 | 3.13E+04 | 9.25E+05 | 0.03388 | 62 | -3.38E+04 | 8.06E+05 | -0.0419 |
| 28 | 3.24E+04 | 9.64E+05 | 0.0336 | 63 | -3.27E+05 | 8.98E+05 | -0.36362 |
| 29 | -4.26E+04 | 9.38E+05 | -0.04541 | 65 | -7.14E+04 | 1.41E+06 | -0.05072 |
| 30 | -1.66E+05 | 9.56E+05 | -0.17388 | 66 | -9.87E+04 | 7.63E+05 | -0.12928 |
| 31 | -8.23E+04 | 8.95E+05 | -0.09195 | 68 | -4.10E+05 | 9.35E+05 | -0.43845 |
| 32 | -5.51E+05 | 1.12E+06 | -0.49217 | 69 | -6.07E+05 | 1.74E+06 | -0.34951 |
| 33 | -2.26E+05 | 8.07E+05 | -0.27951 | 70 | -6.79E+05 | 9.59E+05 | -0.70759 |
| 35 | -2.49E+05 | 9.47E+05 | -0.26318 | 72 | -6.69E+05 | 1.26E+06 | -0.52874 |
| 37 | -1.68E+05 | 7.16E+05 | -0.2346 | 73 | -1.07E+06 | 1.06E+06 | -1.00985 |

**Table S19. NHNOE/NONOE ratios of p53-TAD$^{D49Y}$.** Values of NONOE and NHNOE intensity represent the integrated peak heights at those assigned positions within their respective spectra for p53-TAD$^{D49Y}$.

| Residue | NONOE Intensity | NHNOE Intensity | Intensity Ratio | Residue | NONOE Intensity | NHNOE Intensity | Intensity Ratio |
|---|---|---|---|---|---|---|---|
| 1 | 1.05E+05 | -6.15E+04 | -1.705844222 | 38 | 4.18E+04 | -7.63E+04 | -0.548272667 |
| 2 | 1.52E+05 | -1.04E+05 | -1.454968718 | 39 | 6.51E+04 | -9.08E+04 | -0.716553093 |
| 3 | 1.33E+05 | -1.53E+05 | -0.86871916 | 40 | 6.90E+04 | -8.84E+04 | -0.780201604 |
| 5 | 4.18E+04 | -6.35E+04 | -0.657933693 | 41 | 4.08E+04 | -1.01E+05 | -0.405845803 |
| 6 | 7.13E+04 | -6.30E+04 | -1.131785051 | 42 | 1.31E+04 | -1.04E+05 | -0.126119786 |
| 7 | 8.41E+04 | -1.44E+05 | -0.585517803 | 43 | 6144.44189 | -1.04E+05 | -0.059216553 |
| 9 | 8472.04102 | -7.83E+04 | -0.108258265 | 44 | -1.58E+04 | -9.62E+04 | 0.164610408 |
| 10 | 2.35E+04 | -4.79E+04 | -0.490018327 | 45 | -2.30E+04 | -8.58E+04 | 0.268656155 |
| 11 | 8.68E+04 | -1.11E+05 | -0.78255834 | 46 | 2420.35571 | -1.08E+05 | -0.022463778 |
| 14 | 1.14E+05 | -1.38E+05 | -0.830771787 | 48 | -4826.18066 | -9.97E+04 | 0.04841567 |
| 15 | 6.28E+04 | -8.77E+04 | -0.715804242 | 49 | 4.02E+04 | -5.66E+04 | -0.709929788 |
| 16 | 1.31E+05 | -9.98E+04 | -1.311871122 | 50 | 3761.53174 | -6.94E+04 | -0.054196665 |
| 17 | 5.26E+04 | -1.39E+05 | -0.37822484 | 51 | 3501.30615 | -1.11E+05 | -0.031608789 |
| 18 | 1.70E+04 | -6.84E+04 | -0.248691262 | 52 | -1.19E+04 | -1.40E+05 | 0.084984656 |
| 19 | 2.75E+04 | -7.94E+04 | -0.345768314 | 53 | 3.86E+04 | -5.52E+04 | -0.698158259 |
| 20 | 6692.53076 | -5.65E+04 | -0.118520345 | 54 | 3.86E+04 | -5.52E+04 | -0.698158259 |
| 21 | 1.96E+04 | -8.60E+04 | -0.228079299 | 55 | 2.86E+04 | -5.11E+04 | -0.560632393 |
| 22 | 3.80E+04 | -6.42E+04 | -0.591631958 | 56 | 4.81E+04 | -9.95E+04 | -0.483292576 |
| 23 | 3352.58276 | -6.95E+04 | -0.048259567 | 57 | 4.55E+04 | -1.24E+05 | -0.367348588 |
| 24 | 2.53E+04 | -8.80E+04 | -0.287913119 | 59 | 5.20E+04 | -1.08E+05 | -0.480235892 |
| 25 | -5795.16895 | -5.77E+04 | 0.100422276 | 61 | 3.60E+04 | -9.22E+04 | -0.390854016 |
| 26 | 2.23E+04 | -5.75E+04 | -0.387473749 | 62 | -1.19E+04 | -1.40E+05 | 0.084984656 |
| 28 | 2.58E+04 | -6.36E+04 | -0.406195291 | 63 | 6.53E+04 | -1.04E+05 | -0.625880144 |
| 29 | 2.19E+04 | -6.01E+04 | -0.363816287 | 65 | 5.26E+04 | -1.39E+05 | -0.37822484 |
| 30 | 3.10E+04 | -5.65E+04 | -0.548609549 | 66 | 2.09E+04 | -6.52E+04 | -0.320831595 |
| 31 | 2.11E+04 | -6.12E+04 | -0.344370405 | 68 | 2.90E+04 | -7.93E+04 | -0.365992021 |
| 32 | 6.01E+04 | -8.74E+04 | -0.68763022 | 69 | 1.16E+05 | -1.11E+05 | -1.044026602 |
| 33 | 2.83E+04 | -7.43E+04 | -0.380659454 | 70 | 9.56E+04 | -9.39E+04 | -1.017542432 |
| 35 | 7.61E+04 | -1.16E+05 | -0.658280177 | 72 | 1.31E+05 | -9.98E+04 | -1.311871122 |
| 37 | 5.27E+04 | -6.94E+04 | -0.759982368 | 73 | 6.98E+04 | -9.02E+04 | -0.773850059 |

**Table S20. NHNOE/NONOE ratios of p53-TAD$^{W53G}$.** Values of NONOE and NHNOE intensity represent the integrated peak heights at those assigned positions within their respective spectra for p53-TAD$^{W53G}$.

| Residue | Intensity A | Intensity B | Intensity Ratio | Residue | Intensity A | Intensity B | Intensity Ratio |
|---|---|---|---|---|---|---|---|
| 1 | 1.70E+05 | -1.01E+05 | -1.6868 | 38 | 7.34E+04 | -1.11E+05 | -0.6591 |
| 2 | 1.94E+05 | -1.62E+05 | -1.19825 | 39 | 1.24E+05 | -1.58E+05 | -0.78666 |
| 3 | 1.36E+05 | -1.65E+05 | -0.82407 | 40 | 1.07E+05 | -1.18E+05 | -0.91188 |
| 5 | 1.05E+05 | -1.16E+05 | -0.90777 | 41 | 1.16E+05 | -1.74E+05 | -0.66866 |
| 6 | 9.71E+04 | -1.05E+05 | -0.92769 | 42 | 1.11E+05 | -1.96E+05 | -0.56593 |
| 7 | 7.47E+04 | -1.31E+05 | -0.57038 | 43 | 6.26E+04 | -1.71E+05 | -0.36649 |
| 9 | 4.41E+04 | -1.36E+05 | -0.32343 | 44 | 1.70E+05 | -2.87E+05 | -0.59231 |
| 10 | 6.32E+04 | -1.46E+05 | -0.432 | 45 | 8.13E+04 | -1.31E+05 | -0.61833 |
| 11 | 8.94E+04 | -1.57E+05 | -0.5697 | 46 | 7.26E+04 | -1.23E+05 | -0.59125 |
| 14 | 1.22E+05 | -2.10E+05 | -0.57907 | 48 | 9.36E+04 | -1.76E+05 | -0.53247 |
| 15 | 8.34E+04 | -1.38E+05 | -0.60342 | 49 | 1.18E+05 | -2.29E+05 | -0.5161 |
| 16 | 2.26E+05 | -1.49E+05 | -1.52028 | 50 | 1.13E+05 | -1.66E+05 | -0.67916 |
| 17 | 1.03E+05 | -2.41E+05 | -0.42604 | 51 | 1.09E+05 | -2.17E+05 | -0.50481 |
| 18 | 4.16E+04 | -1.11E+05 | -0.37502 | 52 | 1.30E+05 | -1.51E+05 | -0.86407 |
| 19 | 4.91E+04 | -1.24E+05 | -0.3961 | 53 | 1.39E+05 | -1.37E+05 | -1.01398 |
| 20 | 2.70E+04 | -1.12E+05 | -0.24064 | 53 | - | - | - |
| 21 | 3.16E+04 | -1.25E+05 | -0.25206 | 55 | 9.12E+04 | -1.35E+05 | -0.67344 |
| 22 | 3.43E+04 | -1.80E+05 | -0.19015 | 56 | 1.47E+05 | -1.90E+05 | -0.77082 |
| 23 | 2.05E+04 | -1.65E+05 | -0.12432 | 57 | 6.84E+04 | -1.70E+05 | -0.40335 |
| 24 | 1.61E+04 | -1.43E+05 | -0.11275 | 59 | 6.90E+04 | -1.96E+05 | -0.35164 |
| 25 | 1.87E+04 | -1.19E+05 | -0.15697 | 61 | 6.37E+04 | -1.33E+05 | -0.47726 |
| 26 | 4.21E+04 | -1.49E+05 | -0.28324 | 62 | 5.00E+04 | -1.64E+05 | -0.30518 |
| 28 | 5.56E+04 | -1.51E+05 | -0.36722 | 63 | 1.32E+05 | -1.97E+05 | -0.6683 |
| 29 | 4.28E+04 | -1.06E+05 | -0.40292 | 65 | 1.02E+05 | -2.41E+05 | -0.42266 |
| 30 | 5.25E+04 | -9.86E+04 | -0.53245 | 66 | 6.60E+04 | -1.26E+05 | -0.52155 |
| 31 | 4.69E+04 | -1.50E+05 | -0.3114 | 68 | 1.09E+05 | -1.64E+05 | -0.6663 |
| 32 | 1.05E+05 | -1.51E+05 | -0.7 | 69 | 1.92E+05 | -1.94E+05 | -0.99162 |
| 33 | 7.97E+04 | -1.30E+05 | -0.6154 | 70 | 2.16E+05 | -1.55E+05 | -1.391 |
| 35 | 1.05E+05 | -1.88E+05 | -0.5594 | 72 | 2.56E+05 | -1.49E+05 | -1.72488 |
| 37 | 8.29E+04 | -1.11E+05 | -0.74882 | 73 | 3.25E+05 | -1.87E+05 | -1.73187 |

**Table S21. Statistics for relaxation and reduced spectral density mapping for p53-TAD variants.**

| P53TAD Variant | R1 (Hz) | R2 (Hz) | NOE | $J(\omega_H)$ (ps/rad) | $J(\omega_N)$ (ps/rad) | J(0) (ns/rad) |
|---|---|---|---|---|---|---|
| WT | 1.95±0.31 | 5.41±0.74 | -0.47±0.27 | 49.3±09.1 | 456±080 | 1.85±0.25 |
| K24N | 1.91±0.19 | 4.46±0.56 | -0.24±0.21 | 41.1±05.4 | 460±056 | 1.46±0.21 |
| N29K/N30D | 1.81±0.22 | 4.38±0.53 | -0.21±0.23 | 37.6±06.1 | 439±060 | 1.45±0.19 |
| D49Y | 1.80±0.19 | 5.32±0.70 | -0.46±0.34 | 45.4±09.2 | 422±058 | 1.85±0.27 |
| W53G | 1.72±0.22 | 5.34±0.70 | -0.60±0.27 | 47.4±06.8 | 393±062 | 1.87±0.26 |
